# A genetic compensatory mechanism regulated by c-Jun and Mef2d modulates the expression of distinct class IIa HDACs to ensure peripheral nerve myelination and repair

**DOI:** 10.1101/2021.09.20.461026

**Authors:** Sergio Velasco-Aviles, Nikiben Patel, Angeles Casillas-Bajo, Laura Frutos-Rincón, Enrique Velasco-Serna, Juana Gallar, Peter Arthur-Farraj, Jose A. Gomez-Sanchez, Hugo Cabedo

## Abstract

The class IIa histone-deacetylases (HDACs) have pivotal roles in the development of different tissues. Of this family, Schwann cells express HDAC4, 5 and 7 but not HDCA9. Here we show that a transcription factor regulated genetic compensatory mechanism within this family of proteins, blocks negative regulators of myelination ensuring peripheral nerve developmental myelination and remyelination after injury. Thus, when HDAC4 and 5 are knocked-out from Schwann cells, a c-Jun dependent mechanism induces the compensatory overexpression of HDAC7 permitting, although with a delay, the formation of a myelin sheath. When HDAC4,5 and 7 are simultaneously removed, the Myocyte-specific enhancer-factor d (Mef2d) binds to the promoter and induces the *de novo* expression of HDAC9, and although several melanocytic- lineage genes are mis- expressed and Remak bundle structure is disrupted, myelination proceeds after a long delay. Thus, our data unveil a finely tuned compensatory mechanism within the class IIa HDAC family, coordinated by distinct transcription factors, that guarantees the ability of Schwann cells to myelinate during development and remyelinate after nerve injury.

## INTRODUCTION

During the postnatal development of the peripheral nervous system (PNS), immature Schwann cells ensheath large caliber axons of sensory and motor neurons and differentiate, forming myelin, a highly specialized plasma membrane that increases nerve impulse velocity by allowing saltatory conduction (Kristjan R. Jessen & Mirsky, 2005). Immature Schwann cells downregulate the transcription factor *c-Jun* (which negatively regulates myelination) and upregulate the expression of transcriptional regulators of myelination such as *Krox20* and *Yy1* (Fazal et al., 2017; Monk et al., 2015; Parkinson et al., 2008). *c-Jun* is strongly re-expressed after nerve injury enabling trans-differentiation of Schwann cells into a repair phenotype that promotes axon regeneration and functional nerve repair (Arthur-Farraj et al., 2012; Gomez-Sanchez et al., 2015). After axon regeneration Schwann cells reestablish contact with them and downregulate *c-Jun*. This allows re-expression of *Krox20* and the consequent reactivation of a gene expression program aimed at remyelination of axons and reestablishment of nerve function (Stassart & Woodhoo, 2020). Activation of Gpr126, a G-protein–coupled receptor that increases intracellular levels of cAMP, is required for Schwann cell myelination and remyelination (Monk et al., 2009, 2011). We have recently shown that the pro-differentiating activity of cAMP is in part mediated by its ability to shuttle HDAC4 into the nucleus of Schwann cells (Gomis-Coloma et al., 2018). Nuclear HDAC4 recruits the complex NcoR1/HDAC3 and deacetylates histone 3 on the promoter of *c-Jun*, repressing its expression. At the same time HDAC4 promotes *Krox20* expression and activation of the myelination program (Velasco-Aviles et al., 2018). *In vivo*, HDAC5 is able to compensate for the loss of HDAC4 expression in Schwann cells and only the removal of both HDAC4 and HDAC5 from Schwann cells leads to myelination defects. Surprisingly by postnatal day 8, myelination in HDAC4/5 double knockout mice proceeds at the same pace as in wild-type nerves, suggesting that there is an additional compensatory mechanism permitting nerve myelination (Gomis-Coloma et al., 2018). Here we show that the *in vivo* elimination of HDAC4 and HDAC5 from Schwann cells induces the overexpression of HDAC7 through a mechanism mediated by the transcription factor c-Jun. Notably, the removal of HDAC7 from Schwann cells in the absence of HDAC4 and HDAC5 produces a much longer delay in myelin development. This demonstrates that overexpressed HDAC7 can partially compensate for the absence of both HDAC4 and HDAC5 in myelinating Schwann cells. Interestingly, non-myelin forming Schwann cells in these triple knock-outs (KOs) mis-express melanocytic lineage genes and fail to properly segregate small caliber axons in the Remak bundles. We show that genetic compensation also plays a pivotal role during remyelination after nerve injury. Thus, and akin to what happens during development, remyelination is delayed when HDAC4 and HDAC5 are removed from Schwann cells. This delay is longer when HDAC7 is also removed, which has a profound impact on nerve impulse conduction during nerve regeneration. Importantly, remyelination in the HDAC4/5/7 triple KO also catches up, supporting the idea that an additional mechanism compensates for the absence of class IIa HDACs. Strikingly, HDAC9, the only class IIa HDAC that is not normally expressed by Schwann cells, is *de novo* expressed in the nerves of the HDAC4/5/7 triple KO mice, induced by the transcription factor Mef2d. These genetic compensatory mechanisms, centering around transcription factors, allow Schwann cells to retain a class IIa HDAC gene dosage high enough to permit eventual myelination during development and remyelination after injury.

## RESULTS

### Upregulation of HDAC7 permits myelination in the absence of HDAC4 and HDAC5

We have previously shown that HDAC4 and HDAC5 redundantly contribute to activate the myelin transcriptional program in Schwann cells *in vivo*. However, although during postnatal development c-Jun levels remain high in the PNS of the HDAC4/5 double conditional knock out mice (*P0-Cre^+/-^; HDAC4^flx/flx^;HDAC5^-/-^*, hereafter called *dKO*), myelination proceeds normally after P8 and adult nerves are morphologically and functionally indistinguishable from those of wild type mice (Gomis-Coloma et al., 2018). In muscle development class IIa HDACs can compensate for each other (Potthoff, Olson, et al., 2007). In addition to HDAC4 and HDAC5, Schwann cells also express HDAC7 (Gomis-Coloma et al., 2018). To test if it can functionally compensate for the absence of HDAC4 and HDAC5, we measured the expression levels of *HDAC7* in the nerves of *dKO.* As shown in Figure 1A, the expression of HDAC7 was substantially induced in the sciatic nerve of the *dKO* mice at P60 (325,1 ± 48,1 %; P=0,0034, n=4), while HDAC9 expression remained residual. This is specific for the *dKO*, as minor or no changes at all were found in the single KOs (Figure S1A and S1B). These results suggest that the simultaneous elimination of HDAC4 and HDAC5 from Schwann cells activates a mechanism aimed to compensate for the drop in the gene dose of class IIa HDACs that upregulates 3-fold the expression of HDCA7. To test whether HDAC7 can functionally compensate to allow myelination in the absence of HDAC4/5, we generated a *HDAC4/5/7* triple Schwann cell specific conditional KO (genotype *P0-Cre^+/-^; HDAC4^flx/flx^;HDAC5^-/-^; HDAC7^flx/flx^*, hereafter called *tKO*; Figures S1C-E). To study myelin development in these mice we evaluated a number of morphological parameters of sciatic nerves at P2, P8 and P21 using transmission electron microscopy (TEM) images. We also quantified the mRNA and protein levels for a number of negative and positive regulators of myelination. We previously separately analyzed Schwann cell gene expression in sciatic nerves of *HDAC4* conditional KO mice (genotype *P0-Cre^+/-^; HDAC4^flx/flx^*, referred to as *cKO4*) mice and global *HDAC5* KO mice (referred to as *KO5*) (Gomis-Coloma et al., 2018). As additional controls, here we also performed a detailed morphological analysis of developing nerves in *cKO4, KO5, dKO* and *HDAC7* Schwann cell specific conditional KO mice (*P0-Cre^+/-^; HDAC7^flx/flx^* referred to as *cKO7*) (Figures S2-S5). Together our data support the view that the individual removal of *HDAC4,5* and *7* has little impact on myelination initiation. By contrast, and in line with our previous results (Gomis-Coloma et al., 2018), the simultaneous elimination of *HDAC4* and *HDAC5* from Schwann cells produced a decrease in the number of myelinated axons at P2 that was almost normalized by P8 (Figure S5). Strikingly, the simultaneous elimination of *HDAC4,5* and *7* from Schwann cells produced a much more pronounced delay in myelin development (Figures 1B-J). Interestingly, expression of the negative regulators of myelination (including *c-Jun*) was notably increased from P2 to P21, which can explain the delay in the expression of myelin genes (Figure 1K-M) and in morphological parameters of myelin development in the *tKO* mice. Thus, our data demonstrates that HDAC7 upregulation can compensate for the absence of HDAC4 and HDAC5 allowing myelination to proceed, although with some delay. Interestingly, and although the coordinated removal of HDAC4, HDAC5 and HDAC7 produces a long delay in myelination, myelin is finally formed and adult *tKO* nerves show almost normal myelination parameters (Figures 2A and 2B).

**Figure 1.**
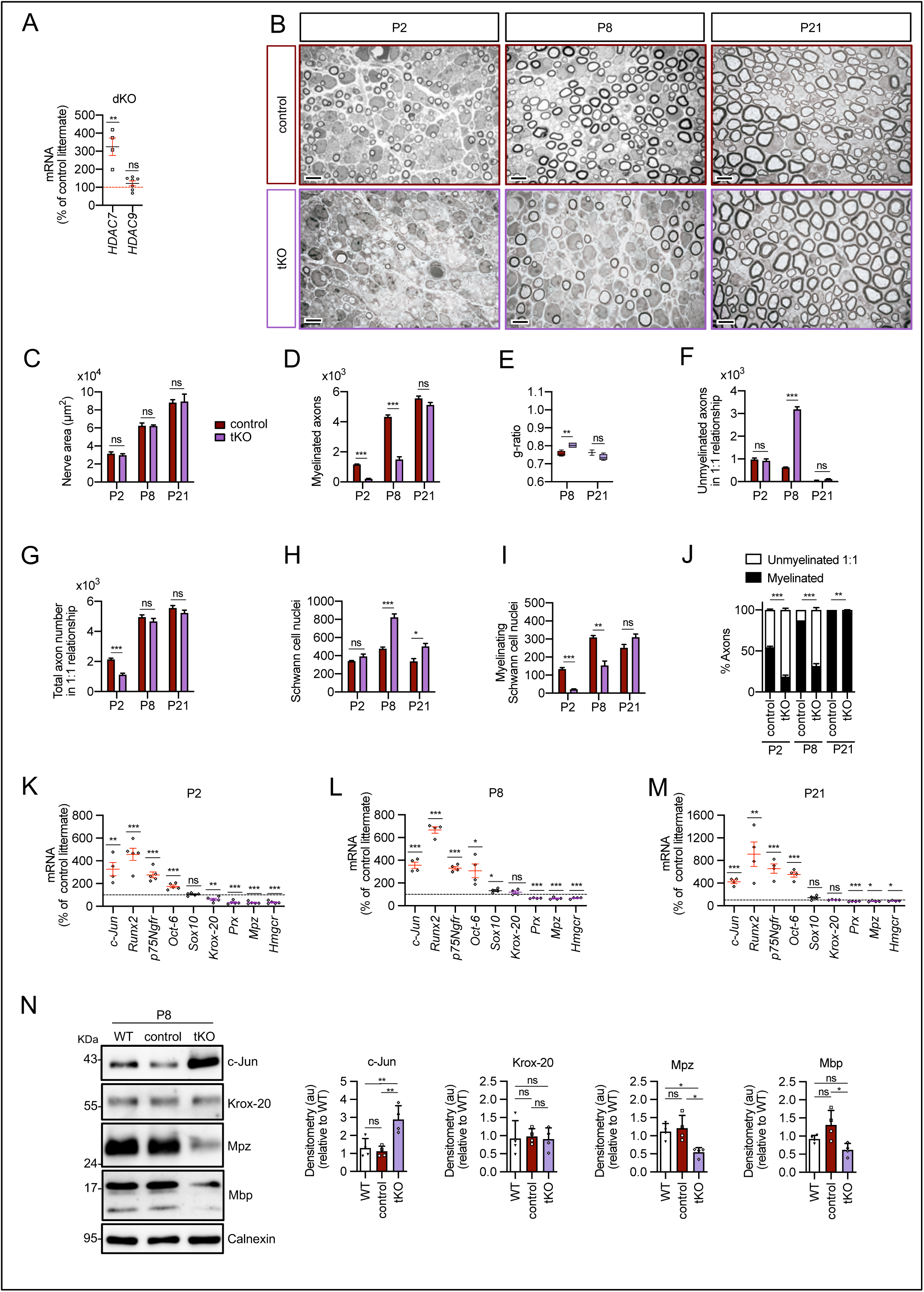
Myelin development is notably delayed in the *tKO* mice. **A)** A 325,1 ± 48,1 % (P=0,0034) increase in the amount of mRNA for HDAC7 was found in the *dKO* nerves. No changes in the expression of HDAC9 were found. RT-qPCR with mouse-specific primers for HDAC7 was performed and normalized to 18S rRNA. The scatter plot, which include also the mean ± SE, shows the fold change in mRNA normalized to control littermates. 4 to 8 mice per genotype were used. Data were analyzed with the unpaired t-test. **B)** Representative transmission TEM mages of P2, P8 and P21 sciatic nerves of *tKO* mice (*P0-Cre^+/−^; HDAC4^flx/flx^; HDAC5^−/−^; HDAC7^flx/flx^*) and the control (*P0- Cre^−/−^*; *HDAC4^flx/flx^; HDAC5^−/−^; HDAC7^flx/flx^*) littermates. Scale bars 5 μm **C)** No statistically significant differences were observed between the area of the *tKO* nerves and control littermates (P2: P=0,5234; P8: P=0,9279; P21: P=0,9009). **D)** The number of myelinated axons is notably decreased at P2 (208 ± 24 in *tKO* versus 1.160 ± 29 in controls; P≤0,0001) and P8 (1.487 ± 179 in *tKO* versus 4.235 ± 129 in controls; P≤0,0001). **E)** *g-ratio* was increased at P8 (0,80 ± 0,01 in the *tKO* versus 0,76 ± 0,01 in control; P=0,0045). **F)** The number of unmyelinated axons in a 1:1 relationship with Schwann cells was notably increased at P8 (3.187 ± 111 in the *tKO* versus 628 ± 21 in controls; P≤0,0001) **G)** The total number of sorted axons in a 1:1 relationship with Schwann cells is decreased at P2 (1.128 ± 90 in the *tKO* versus 2.131 ± 95 in the control; P=0007).**H)** The total number of Schwann cells (counted as nuclei) is increased at P8 (823 ± 37 in the *tKO* versus 476 ± 20 in controls; P≤0,0001) and at P21 (503 ± 31 in the *tKO* versus 337 ± 32 in controls; P≤0,0152). **I)** In contrast, the number of myelinating Schwann cells is decreased at P2 (22 ± 1 in the *tKO* versus 134 ± 8 in controls; P≤0,0001) and at P8 (153 ± 25 in the *tKO* versus 309 ± 11 in controls; P=0,0013). **J)** The percentage of myelinated axons is decreased at P2 (18,5 ± 3,7 % in the *tKO* versus 54,6 ± 1,1 % in controls; P≤0,0001), P8 (31,6 ± 2,9% in the *tKO* versus 54,6 ± 1,1 % in controls; P≤0,0001) and, although much less, at P21 (97,9 ± 0,4 % in the *tKO* versus 99,9 ± 0,0 % in controls; P=0,0135). For these experiments 3 to 4 animals per genotype were used; Unpaired t-test was applied for statistical analysis. **K)** Markers of non-myelin forming Schwann cells are upregulated whereas those of myelin-forming Schwan cells are downregulated in the *tKO*. P2, sciatic nerves were removed and total RNA extracted. RTqPCR with mouse-specific primers for the indicated genes was performed and normalized to 18S rRNA. Graph shows the percentage of mRNA for each gene in the *tKO* normalized to the control littermates. A scatter plot is shown with the results obtained, which include also the mean ± SE. **L)** The same for P8. **M)** The same for P21.For these experiments, 4 to 5 mice per genotype and age were used. Data were analyzed with the unpaired t-test. **N)** A representative WB of protein extracts from *tKO*, control, and wild type P8 nerves is shown. In the quantification, c-Jun protein increased in the *tKO* (2,88 ± 0,19 in the *tKO* versus 1,12 ± 0,071 in the control nerve; P=0,004). Mpz protein was found decreased (0,55 ± 0,03 in the *tKO* versus 1,21 ± 0,09 in the control nerve; P=0,0115) as was Mbp (0,62 ± 0,045 in the *tKO* versus 1,31 ± 0,100 in the control nerve; P=0,012) We couldn’t find changes in Krox20. Densitometric analysis was done for 7 to 9 WB from the same number of mice and normalized to the WT. Data were analyzed with the One-way ANOVA Tukey’s test (* P<0,05; ** P<0,01; *** P<0,001; ns: no significant). See Source data section online (Graphs source data) for more details.

**Figure 2.**
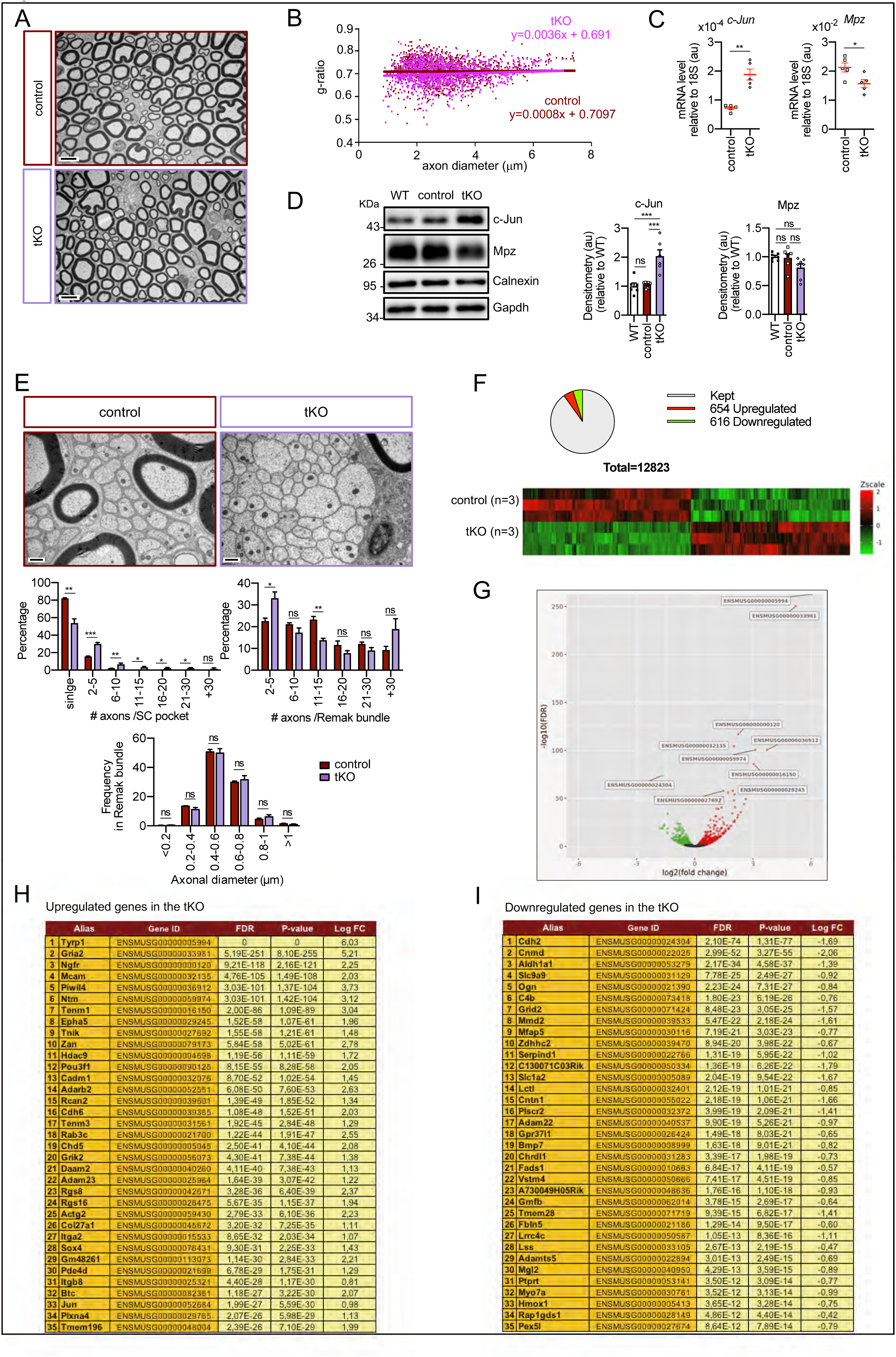
Characterization of the *tKO*. **A)** Representative transmission TEM image of the sciatic nerve of and adult (P60) *tKO* mouse and a control littermate. **B)** Scatter plot of *g-ratio* versus axon diameter. 1100 axons of 4 different mice per genotype were used. No changes in *g-ratio* were detected. **C)** mRNA for *c- Jun* remains increased in the *tKO* by 2,6-fold (1,88 ± 0,19 x10^-4^ au in the *tKO* versus 0,72 ± 0,05 x10^-4^ au in controls; P=0,003) whereas *Mpz* was slightly decreased (1,57 ± 0,13 x10^-2^ au in the *tKO* versus 2,13 ± 0,05 x10^-2^ au in controls; P=0,027). RTqPCR with mouse-specific primers for the indicated genes was performed. Graph shows a scatter plot for the ΔCt (which include also the mean ± SE) of the gene normalized to the housekeeping 18S. 5 mice per genotype and age were used. Data were analyzed with the unpaired t-test).**D)** c-Jun and Mpz protein levels. A representative Western blot of protein extracts from wild type (C57BL/6), control and *tKO* sciatic nerves is shown. The densitometric analysis of 6 to 7 different experiments normalized to WT is also shown. Data were analyzed with the unpaired t-test. Only for c-Jun were detected consistent changes (2,04 ± 0,22 in the *tKO* versus 1,05 ± 0,04 in controls; P=0,0003) at the protein level (*** P<0,001). **E)** Failed segregation of the axons in the Remak bundles of the *tKO.* A representative high power TEM image is shown. Morphometric analysis shows that axon diameter distribution is preserved in the *tKO*, but the number of axons per Remak bundle and the distribution of axon per pocket is changed. 500 axons from 4 animals per genotype were counted. Mixed model ANOVA with Bonferroni post-hoc test was used for comparations. **F)** Pie chart and DEG heatmap of the RNAseq analysis of P60 showing the distribution of changed genes in the *tKO*. **G)** Volcano plot shows that the most robustly changed genes were upregulated. ENSEMBL indemnification numbers for the 10 most robustly changed genes is shown. **H)** List of the 35 most upregulated genes in the adult (P60) *tKO* classified by FDR. **I)** List of the 35 most downregulated genes in the adult (P60) *tKO* classified by FDR. (* P<0,05; ** P<0,01; *** P<0,001; ns: no significant). See Source data section online (Graphs source data) for more details.

### Defects in Remak Schwann cell differentiation in the *tKO*

Despite PNS myelination looks normal in the adult (p60) *tKO* mice we found the Remak bundles profoundly altered in these nerves, with many axons not properly segregated (Figure 2E). Thus, there is a significant increase in the number of pockets with 2-5 axons and, although it is very rare to find pockets with more than 5 axons in the control (2,3%), an important number of axons are grouped together in packs of more than 5 in the *tKO* (16,5%), with some of them being in pockets of more than 30 axons. Importantly, axon diameter distribution is not changed, which points towards an intrinsic defect in axon segregation more than a sorting problem (Figure 2E). Whole genome-wide transcriptome analysis showed 654 upregulated and 616 downregulated in the nerves of adult *tKO* (Figure 2F and Source data section online (RNAseq source data)). Volcano plot shows that genes tended towards being more strongly upregulated than downregulated. Surprisingly, the most robustly upregulated gene is the Tyrosinase-related protein 1 encoding gene (*Tyrp1*; Log FC= 6,03; FDR=0), a melanocyte lineage specific gene (Figures 2G and 2H). Additionally the melanoma cell adhesion molecule *Mcam* and *p75Ngfr* are also highly induced in the sciatic nerves of the *tKO*. RTqPCR confirmed the strong induction of *Tyrp1* and *Mcam* in the sciatic nerves of the *tKO* (P60), but not in the single *KOs* neither *dKO* nerves (Figures 3A and 3B). Interestingly we found also increased the expression of Microphthalmia-associated transcription factor (*Mitf*) and the Endothelin B receptor (*Ednrb*), two other genes of the melanocytic lineage (Figures 3C and 3D). Importantly, the expression of all these genes increased from early in postnatal development (Figures S6B and S6C). Western blot analysis (Figures 3E and 3F) and confocal microscopy confirmed these findings and showed that the mis-expression of melanocytic lineage markers is confined to the non-myelin forming Schwann cells of the Remak bundles (Figures 3G-K). Thus, class IIa HDACs are necessary to allow Schwann cell precursors to differentiate into Remak Shcwann cells and properly segregate small size axons.

**Figure 3.**
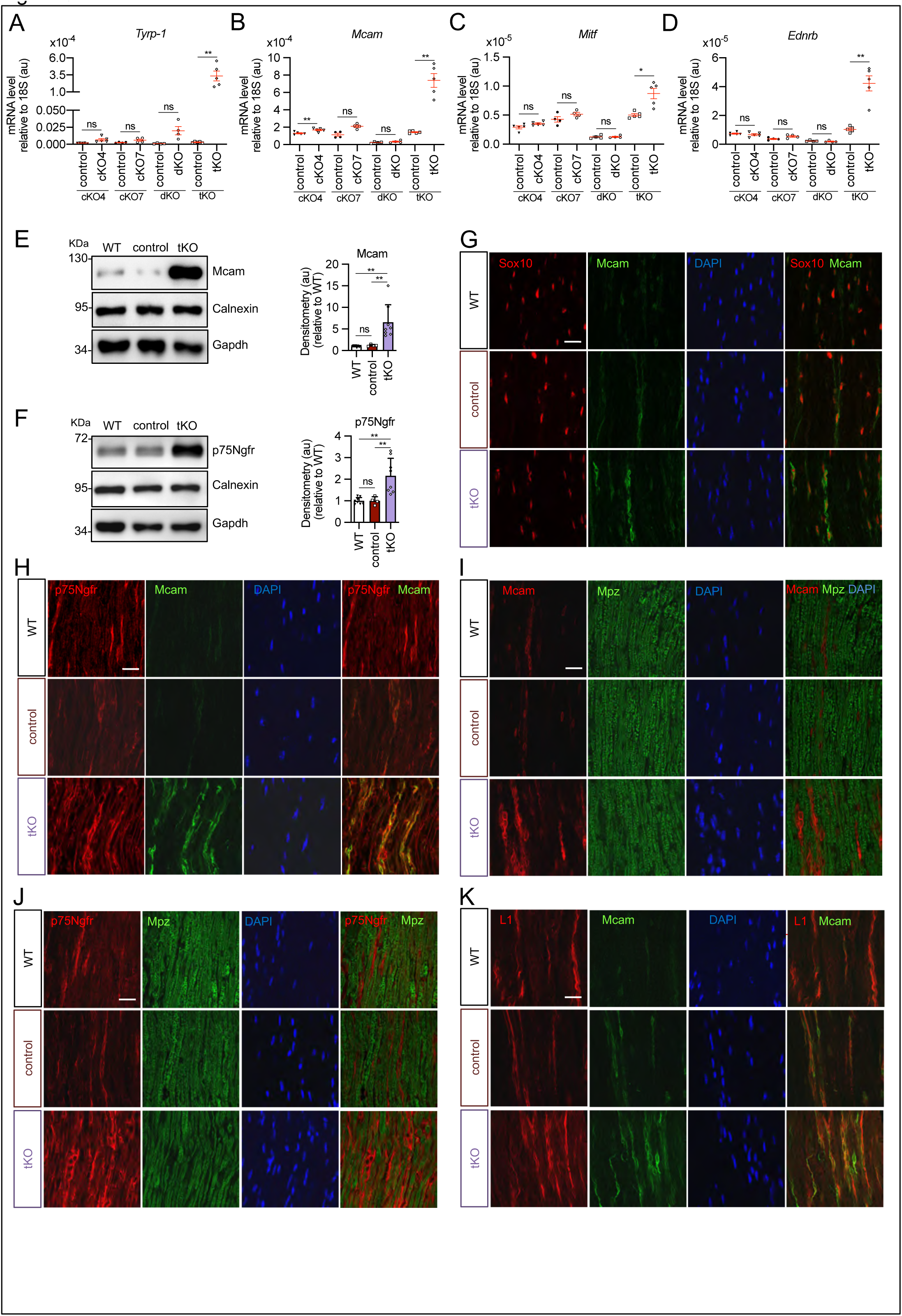
Melanocyte lineage markers are expressed by non-myelinating Schwann cells of the Remak bundles in the sciatic nerves of the *tKO*. A) mRNA for *Tyrp-1* is dramatically increased by 1.081-fold in the *tKO* (3,35 ± 0,71 x10^-4^ au in the *tKO* versus 0,11± 0,05 x10^-6^ au in controls; P=0,0092) whereas no changes were found in the *cKO4*, *cKO7* neither *dKO* sciatic nerves. B) mRNA for *Mcam* is also upregulated (5,13-fold) in the *tKO* (7,39 ± 0,79 x10^-4^ au in the *tKO* versus 1,44 ± 0,06 x10^-4^ au in controls) with only minor o no changes at all for the other genotypes. The same although less marked (1,74-fold) for *Mitf* ( 0,87 ± 0,09 x10^-5^ au in the *tKO* versus 0,50 ± 0,02 x10^-5^ au in controls; P=0,0128) (**C**) and *Ednrb* (4,1-fold; 4,23 ± 0,52 x10^-5^ au in the *tKO* versus 1,04 ± 0,09 x10^-5^ au in controls; P=0,0032) (**D**). RTqPCR with mouse-specific primers for the indicated genes was performed. Graph shows a scatter plot for the ΔCt (which include also the mean ± SE) of the gene normalized to the housekeeping 18S. 4 to 5 mice per genotype were used. Data were analyzed with the unpaired t-test with Welch’s correlation. **E)** Mcam protein levels in the sciatic nerves of the *tKO*. A representative Western blot of protein extracts from wild type (C57BL/6), control and *tKO* sciatic nerves is shown. Mcam protein was increased by 7,6-fold in the *tKO* (9,93 ± 1,75 au in the *tKO* versus 1,30 ± 0,13 in controls; P=0,0003). **F)** p75Ngfr protein was increased by 2,15-fold (2,16 ± 0,29 in the *tKO* versus 1,005 ± 0,09 in controls; P=0,0003).4 to 8 WB of the same number of animals per genotype were quantified. Data were analyzed with the One-way ANOVA Tukey’s test. **G)** Mcam signal colocalizes with Sox10. **H)** Mcam signal colocalizes with p75Ngfr. **I)** Mcam is not expressed by myelin forming Schwann cells (Mpz^+^). **J)** Same happens with p75Ngfr. **K)** Mcam signal colocalizes with L1cam, a marker of the non-myelin forming Schwann cells of the Remak bundles. P60 sciatic nerves were fixed and submitted to immunofluorescence with the indicated antibodies. Nuclei were counterstained with Hoechst. Representative confocal images of sections obtained from the sciatic nerves of wild type (WT), control and *tKO* mice are shown. Scale bars= 20 μm (* P<0,05; ** P<0,01; *** P<0,001; ns: no significant). See Source data section online (Graphs source data) for more details.

### Remyelination kinetics depends on class IIa HDAC gene dose

The molecular mechanisms of Schwann cell remyelination share similarities to myelination during development, however there are also notable differences (Stassart & Woodhoo, 2020). Given the role of class II a HDACs in myelination we asked whether they are also involved in remyelination after nerve injury. To address this, we first performed crush experiments in the sciatic nerves of 8- week-old *cKO4* mice. As controls, we used *P0-Cre^-/-^;HDAC4 ^flx/flx^* littermates (Figures S7A-K) . At 10 days post injury (dpi), we found a small decrease in the percentage of myelinated axons, with an increase in the number of unmyelinated axons with a diameter > 1,5 μm in a 1:1 relationship with the Schwann cells (Figure S7F and S7K). A small increase in the number of Schwann cell nuclei was also found at 20 dpi and 30 dpi (Figure S7I). Thus, HDAC4 removal has, although limited, an impact on remyelination that is compensated for after 10 dpi. Analysis of remyelination in *KO5* mice and wild type littermates (*HDAC5^+/+^)* showed no differences (Fig S8). Also, no relevant differences in Schwann cell and myelin protein gene expression in either *cKO4* or *KO5* injured nerves were found (Figures S7L-N and S8L-M). Myelin clearance was also normal in both genotypes (Figures S7O and S8N). Thus, similar to development, individual removal of HDAC4 or HDAC5 has very little impact on remyelination after nerve injury. To explore whether there is also genetic compensation within class IIa HDACs in Schwann cells after injury, we analysed the *dKO* crushed nerve. In this case, the percentage of myelinated axons at 10 dpi was notably decreased (15,5 ± 2,3 % in the *dKO* versus 60,4 ± 4,8 % in the control; P≤0,0001) (Figures 4A and 4K). Total axon counts were similar between genotypes suggesting that this difference was not due to an axon regeneration defect (Figure 4G). At 20 dpi the difference between both genotypes was reduced and normalized at 30 dpi. A notable increase in the number of unmyelinated axons with a diameter > 1,5 μm in a 1:1 relationship with the Schwann cells was found at 10 dpi and at 20 dpi that almost normalized at 30 dpi (Figure 4F). Furthermore, *g-ratio* was increased at 10 dpi and at 20 dpi but showed no difference between control and *dKO* nerves at 30dpi (Figure 4D). Quantifying gene expression at 10 dpi demonstrated that mRNA for *c-Jun* remains higher in the *dKO* injured nerves, as does *Gdnf* (Figure 4L). In the same line, we found a significant decrease in the mRNA for *Krox20*, *Periaxin*, *Mpz,* and *Mbp*. We did not find changes in the expression of *Runx2* or *Oct6* (Figures 4L and 4M). To substantiate these results, we looked at protein levels, and found c-Jun protein remained higher in the *dKO* at 10 dpi and was unchanged at 21 dpi. Though there was no difference in Krox20, Mpz protein levels were decreased at 21 dpi. Together, our data supports the view that the simultaneous removal of HDAC4 and HDAC5 from Schwann cells produces a moderate delay in remyelination after nerve injury. This remyelination delay could be caused by an intrinsic problem in the capacity of Schwann cells to reactivate the myelination program, but could also be secondary to a failure in the ability of myelinating Schwann cells to acquire the repair phenotype and clear myelin debris in the distal stump. However, we did not favor the second explanation because markers of the Schwann cell repair phenotype, such as *Bdnf* and *Olig 1*, in addition to *c-Jun,* are highly expressed in the *dKO* at 10 dpi (Figure 4L and N). Furthermore, we did not find any change in the number of intact myelin sheaths at 4 days after cut in the *dKO*, or in clearance of Mpz protein, suggesting no effect on the rate of demyelination. Finally, repair program genes were normally upregulated in the dKO (Figures S9A-C).Together, our data show that the delay in remyelination of the *dKO* is due to an intrinsic defect of Schwann cells to activate the myelin transcriptional program and not a consequence of an altered reprograming capacity to the repair phenotype or to delayed myelin clearance.

**Figure 4.**
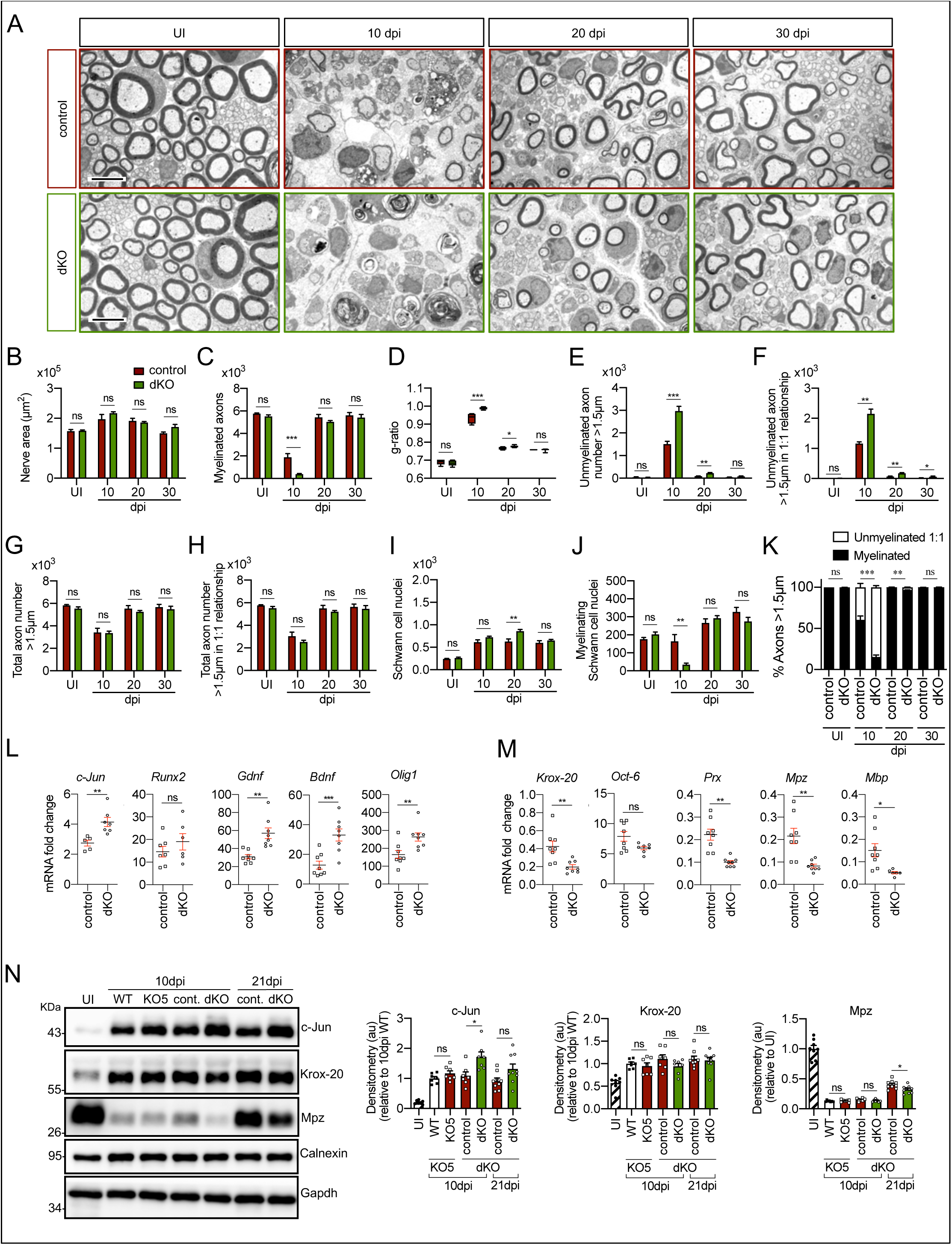
Remyelination is delayed in the nerves of the *dKO* mice. **A)** Representative transmission TEM images of P60 sciatic nerves uninjured (UI) and 10, 20 and 30 days post crush (dpi) of *dKO* (*P0-Cre^+/−^; HDAC4^flx/flx^; HDAC5^-/-^*) and the control (*P0-Cre^−/−^*; *HDAC4^flx/flx^; HDAC5^−/−^*) littermates are shown. Scale bars 5 μm. **B)** No statistically significant differences were observed in the area of the *dKO* nerves and control littermates (UI: P=0,804; 10 dpi: P=0,195; 20 dpi: P=0,559; 30 dpi: P=0,0594). **C)** The number of myelinated axons is notably decreased at 10 dpi (388 ± 55 in the *dKO* versus 1.889 ± 330 in the control; P=0,0005). **D)** *g-ratio* was increased at 10 dpi (0,989 ± 0,003 in the *dKO* versus 0,934 ± 0,015 in control (P=0,002)) and at P21 (0,776 ± 0,003 in the *dKO* versus 0,767 ± 0,003 in control (P=0,043). **E)** The number of unmyelinated axons in a 1:1 relationship with Schwann cells was notably increased at 10 dpi (2.969 ± 203 in the *dKO* versus 1.512 ± 119 in controls; P=0,0007) and at 20 dpi (224 ± 25 in the *dKO* versus 88 ± 14 in controls; P=0,0016). **F)** The total number of unmyelinated axons in a 1:1 relationship with Schwann cells is increased at 10 dpi (2.148 ± 155 in the *dK*O versus 1.158 ± 56 in the control; P=0011) at 20 dpi (175 ± 20 in the *dKO* versus 68 ± 12 in the control; P=002) and at 30 dpi (63 ± 17 in the *dKO* versus 22 ± 5 in the control; P=0,043).**G)** No changes in the total axon number was found (UI: P=0,157; 10 dpi: P=0,910; 20 dpi: P=0,349; 30 dpi: P=0,666). **H)** neither in the total sorted axon number (UI: P=0,193; 10 dpi: P=0,169; 20 dpi: P=0,294; 30 dpi: P=0,682). **I)** The total number of Schwann cells (counted as nuclei) was increased at 20 dpi (861 ± 34 in the *dKO* versus 630 ± 53 in controls; P=0,0041). **J)** In contrast, the number of myelinating Schwann cells was found decreased at 10 dpi (35 ± 8 in the *dKO* versus 164 ± 37 in controls; P=0,0032). **K)** The percentage of myelinated axons is decreased at 10 dpi (15,5 ± 2,3 % in the *dKO* versus 60,4 ± 4,8 % in controls; P<0,0001), 20 dpi (96,6 ± 0,4% in the *dKO* versus 98,8 ± 0,2 % in controls; P=0,0016) and, although much less, at P21 (98,9 ± 0,3 % in the *dKO* versus 99,6 ± 0,1 % in controls; P=0,0482). For these experiment 3 to 6 animals per genotype were used; Unpaired t-test was applied for statistical analysis. **L)** Expression of several negative regulators of myelination and repair Schwann cell markers is enhanced at 10 dpi in the sciatic nerves of the *dKO*: *c-Jun* (1,51 -fold; P=0,0056), *Gdnf* (1,85-fold; P=0,0025), *Bdnf* (2,60-fold; P=0,001) and *Olig 1* (1,60-fold; P=0,008). **M)** Expression of positive regulators and myelin genes is decreased at 10 dpi in the sciatic nerves of the *dKO*: *Krox20* (0,47-fold; P0,0068), *Prx* (0,45-fold; P=0,001), *Mpz* (0,33-fold; P=0,005) and *Mbp* (0,33-fold; P=0,012). RTqPCR with mouse-specific primers for the indicated genes was performed and normalized to 18S rRNA. The scatter plot, which include also the mean ± SE, shows the fold change of mRNA for each gene at 10 dpi normalized to the uninjured nerve. 5 to 8 mice per genotype were used. Data were analyzed with the unpaired t-test with Welch’s correlation. **N)** A representative WB of protein extracts from *dKO*, control, *KO5 ^-/-^* and wild type nerves is shown. In the quantification, c-Jun protein remains higher in the *dKO* at 10 dpi (1,72 ± 0,17-fold; P=0,012) and tend to equalize at 21 dpi. Mpz protein was found decreased by 0,32 ± 0,02-fold at 21 dpi (P=0,02), however we couldn’t find changes in *Krox20* (*KO5 ^-/-^* mice were used to compare with the wild types littermates). Densitometric analysis was done on 7 to 9 WB from the same number of mice and normalized to 10 dpi WT. Data were analyzed with the unpaired t-test. (* P<0,05; ** P<0,01; *** P<0,001; ns: no significant). See Source data section online (Graphs source data) for more details.

Since remyelination is only moderately delayed in the *dKO,* we asked whether HDAC7 was able to functionally compensate in a similar way as in development (previously, we could establish that *cKO7* mice have no defects in myelin clearance, injury induced gene expression or remyelination after an injury (Figure S10)). First, we found that *tKO* Schwann cells upregulate repair program genes after a cut injury (Figure S11) and interestingly some of them (*c-Jun*, *Olig-1* and *Shh*) appear to be overexpressed at some time points, suggesting that the class II a HDAC act as a brake on the initial induction of the Schwann cell repair phenotype (Figures S11A-C). In line with this observation, myelin was more rapidly cleared in these mutants (Figures 5A-C). On assessment of *tKO* nerves after a crush injury (Figures 5D-N), strikingly, we could not find any myelinated axon profile in the four *tKO* sciatic nerves analyzed at 10 dpi. This is in contrast to the controls which showed myelin profiles in 73 ± 3,1 % of axons (P≤0,0001) (Figure 5N). At 20 dpi, the *tKO* still had only 19,2 ± 5,5 % of the axons myelinated, whereas almost all large caliber axons were myelinated in the control (98,1 ± 0,2 %; P=0,0001). Moreover, at 30 dpi only 60,3 ± 6,1 % of the axons were myelinated (P=0,0031) in the *tKO* mice. At 60 dpi, myelination was only slightly delayed in the *tKO* (Figure 5N). Differences in *g-ratios* followed the same pattern (Figure 5G). In the same line we found a notable increase in the number of unmyelinated axons >1,5 μm at 10 dpi, that decreases slowly but progressively up to 60 dpi, when it approaches a similar number to the control (Figure 5H). Interestingly most of these unmyelinated axons are in a 1:1 relationship with Schwann cells (Figure 5I), suggesting that the delay is in the transition from the pro-myelinating to the myelinating Schwann cell stage. We also found a notable increase in the number of Schwann cells per nerve section that was maintained after 60 dpi (Figure 5L), and an increase in the nerve area (Figure 5E). The increased number of Schwann cells is probably consequence of over-proliferation, as suggested by Ki67 staining (Figure S12). We also observed a decrease in the number of axons >1,5 μm at 20 and 30 dpi (Figure 5J), probably because the smaller diameter of the unmyelinated axons. Finally, the delay in remyelination was substantiated by Western blot. As shown in Figure 5O, c-Jun protein is clearly more abundant in the nerves of the *tKO* than in either control littermates or wild types, both at 10 dpi and 21 dpi. Conversely, the amount of Mpz protein is lower at 10 and 21dpi. Krox20 is also lower at 10 dpi in the *tKO*, but levels had recovered by 21 dpi.

**Figure 5.**
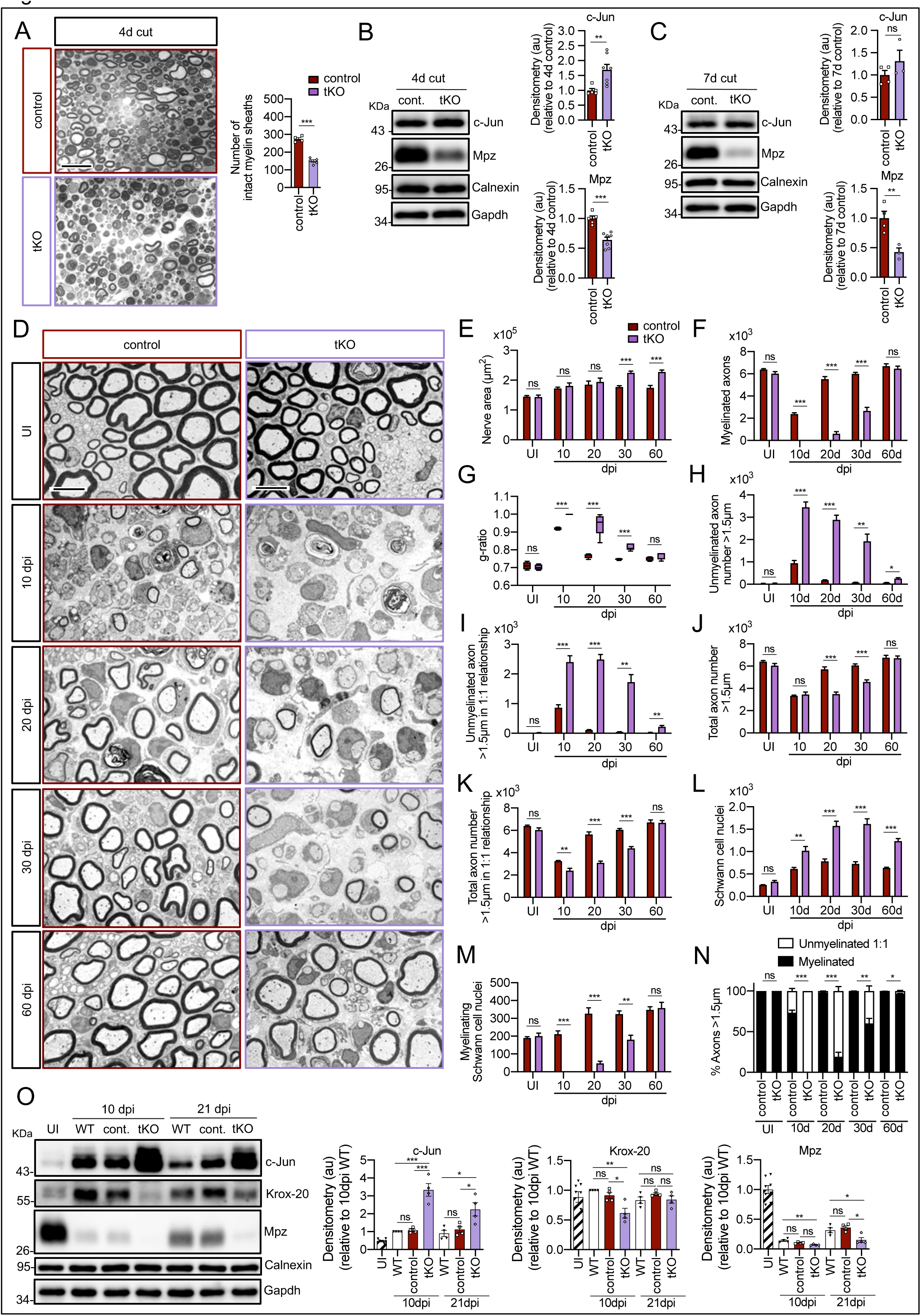
Remyelination is dramatically delayed in the *tKO*. **A)** Myelin clearance is accelerated in the sciatic nerves of the *tKO*. A representative toluidine blue staining image of 4 days-cut sciatic nerve of *tKO* and control mice is shown. The quantification of intact myelin sheaths shows a 0,55-fold change (P<0,0001) in the *tKO* (150 ± 7 intact myelin sheaths in the *dKO* versus 274 ± 8 in controls; P<0,0001). 7 to 8 animals per genotype were used for the experiment. Data were analyzed with the unpaired t-test. **B)** WB supports an accelerated myelin clearance in the *tKO*. As is shown, Mpz protein is decreased (0,7-fold; P=0,0062) in the *tKO* .As expected, c-Jun was increased. 4 mice per genotype were used. **C)** At 7 days post-cut the decrease in the Mpz protein was more marked (0,43-fold; P=0,012). 3 to 4 mice per genotype were used. Calnexin and Gapdh were used as housekeepings. Densitometric analysis was preformed and normalized to the controls. Data were analyzed with the unpaired t-test. **D)** Representative transmission TEM images of P60 sciatic nerves uninjured (UI) and 10, 20, 30 and 60 days post crush (dpi) of *tKO* and the control littermates are shown. Scale bars 5 μm. **E)** No statistically significant differences were observed for the area of the *tKO* nerves and control littermates in the UI (P=0,897), at 10 dpi (P=0,456) neither at 20 dpi (P=0,602). The nerve area of the *tKO* was increased at 30 dpi (1,26-fold; P=0,0002) and at 60 dpi (1,30-fold; P=0,0008). **F)** No myelinated axons in the *tKO* were found at 10 dpi whereas the control had 2.383 ± 112 axons myelin (P<0,0001). Myelinated axons were decreased at 20 dpi (606 ± 200 in the *tKO* versus 5.525 ± 222 in the control; P<0,0001), at 30 dpi (2.659 ± 323 in the *tKO* versus 6.003 ± 125 in the control; P<0,0001), to catch up at 60 dpi (6.458 ± 240 in the *tKO* versus 6.689 ± 212 in the control: P=0,491). **G)** *g-ratio* was increased in the *tKO* at 10 dpi (1 ± 0 in the *tKO* versus 0,92 ± 0,003 in control; P<0,0001) at 20 dpi (0,94 ± 0,027 in the *tKO* versus 0,77 ± 0,006 in control; P=0,0023) and at 30 dpi (0,82 ± 0,008 in the *tKO* versus 0,75 ± 0,002 in control; P=0,0009). **H)** The number of unmyelinated axons >1,5 was increased at 10 dpi (3.477 ± 236 in the *tKO* versus 950 ± 116 in controls; P<0,0001), at 20 dpi (2.885 ± 209 in the *tKO* versus 184 ± 15 in controls; P=0,0002), at 30 dpi (1.925 ± 319 in the *tKO* versus 76 ± 16 in controls; P=0,0044), and at 60 dpi (257 ± 43 in the *tKO* versus 69 ± 11 in controls; P=0,010). **I)** The number of unmyelinated axons >1,5 in a 1:1 relationship with Schwann was notably increased at 10 dpi (2.405 ± 209 in the *tKO* versus 864 ± 102 in controls; P=0,0006), at 20 dpi (2.487 ± 170 in the *tKO* versus 110 ± 13 in controls; P=0,0001), at 30 dpi (1.728 ± 250 in the *tKO* versus 43 ± 8 in controls; P=0,0025), and at 60 dpi (224 ± 43 in the *tKO* versus 28 ± 5 in controls; P=0,010). **J)** In contrast, the total number of axons >1,5 μm was decreased at 20 dpi (3.492 ± 184 in the *tK*O versus 5.790 ± 228 in the control; P<0,0001) and at 30 dpi (4.584 ± 184 in the *tKO* versus 6.080 ± 131 in the control; P=0,0002) to finally catch up at 60 dpi (6.716 ± 198 in the *tKO* versus 6.758 ± 221 in the control; P=0,889). **K)** The total number of axons >1,5 μm sorted (in a 1:1 relationship with Schwann cells) was decreased at 10 dpi (2.405 ± 209 in the *tK*O versus 3.247 ± 60 in the control; P<0,0082), at 20 dpi (3.093 ± 147 in the *tK*O versus 5.653 ± 233 in the control; P<0,0001) and at 30 dpi (4.387 ± 158 in the *tK*O versus 6.046 ± 127 in the control; P<0,0001). **L)** The number of Schwann cells (counted as nuclei) was increased at 10 dpi (1.017 ± 95 in the *tK*O versus 609 ± 33 in the control; P<0,0001), at 20 dpi (1.576 ± 100 in the *tK*O versus 779 ± 54 in the control; P<0,0001) and at 30 dpi (1.618 ± 116 in the *tK*O versus 723 ± 47 in the control; P<0,0001). **M)** The number of myelinating Schwann cells was also decreased at 10 dpi (0 ± 0 in the *tK*O versus 212 ± 18 in the control; P<0,0001), at 20 dpi (48 ± 12 in the *tK*O versus 326 ± 33 in the control; P<0,0001) and at 30 dpi (181 ± 24 in the *tK*O versus 325 ± 17 in the control; P=0,0011). **N)** The percentage of myelinated axons is decreased at 10 dpi (0 ± 0 % in the *tKO* versus 73,1 ± 3,1 % in controls; P<0,0001), 20 dpi (19,2 ± 5,5% in the *tKO* versus 98,1 ± 0,2 % in controls; P<0,0001), 30 dpi (60,3 ± 6,1% in the *tKO* versus 99,3 ± 0,1 % in controls; P=0,0031) and at 60 dpi (96,6 ± 0,7% in the *tKO* versus 99,6 ± 0,1 % in controls; P<0,0001). For these experiments 4 to 5 animals per genotype were used; Unpaired t-test was applied for statistical analysis. **O)** A representative WB of protein extracts from *tKO*, control and wild type nerves is shown. In the quantification, c-Jun protein remains higher in the *tKO* at 10 dpi (3,24 ± 0,35-fold; P<0,0001) and at 21 dpi (2,25 ± 0,38- fold;P=0,031). *Krox20* was found decreased at 10 dpi (0,61 ± 0,08-fold; P=0,011). Mpz protein was found decreased at 21 dpi by 0,15 ± 0,04-fold dpi (P=0’0091). Densitometric analysis was done for 7 to 9 WB from the same number of mice and normalized to 10 dpi WT. Data were analyzed with the unpaired t-test.( *P<0,05; ** P<0,01; *** P<0,001; ns: no significant). See Source data section online (Graphs source data) for more details.

To gain insight into the functional consequences of the remyelination delay in the *tKO* we performed nerve impulse conduction studies of the sciatic nerves after crush injury (see Material and methods). In uninjured nerves, we found no differences in voltage amplitude or nerve conduction velocity between *tKO* and controls (Figure 6A-D) (curiously we observed a smaller amplitude and slower nerve conduction velocity for all genotypes when compared with wild type nerves (Figure S13), probably due to the absence of HDAC5 in neurons). By contrast, at 40 dpi, whereas 6 of 9 sciatic nerves of control mice showed a response when electrically stimulated at 8V, only 1 of 8 *tKO* responded (Figures 6E and 6F). The same distribution was found at 10 V. At 15V, 8 of 9 control mice responded while only 4 of 8 *tKO* responded. In the same line, the amplitude of the A-fiber component of the compound action potential was decreased for 8, 10 and 15V stimuli (Figure 6G). Regarding the component corresponding to C fibers, amplitude was also decreased for 8 and 10V stimulation (Figure 6H). Moreover, nerve conduction velocity showed a statistically significant decrease when using 15V stimuli (Figure 6I). All together our data demonstrate that the kinetics of remyelination after nerve injury is directly correlated with class IIa HDAC gene dose.

**Figure 6.**
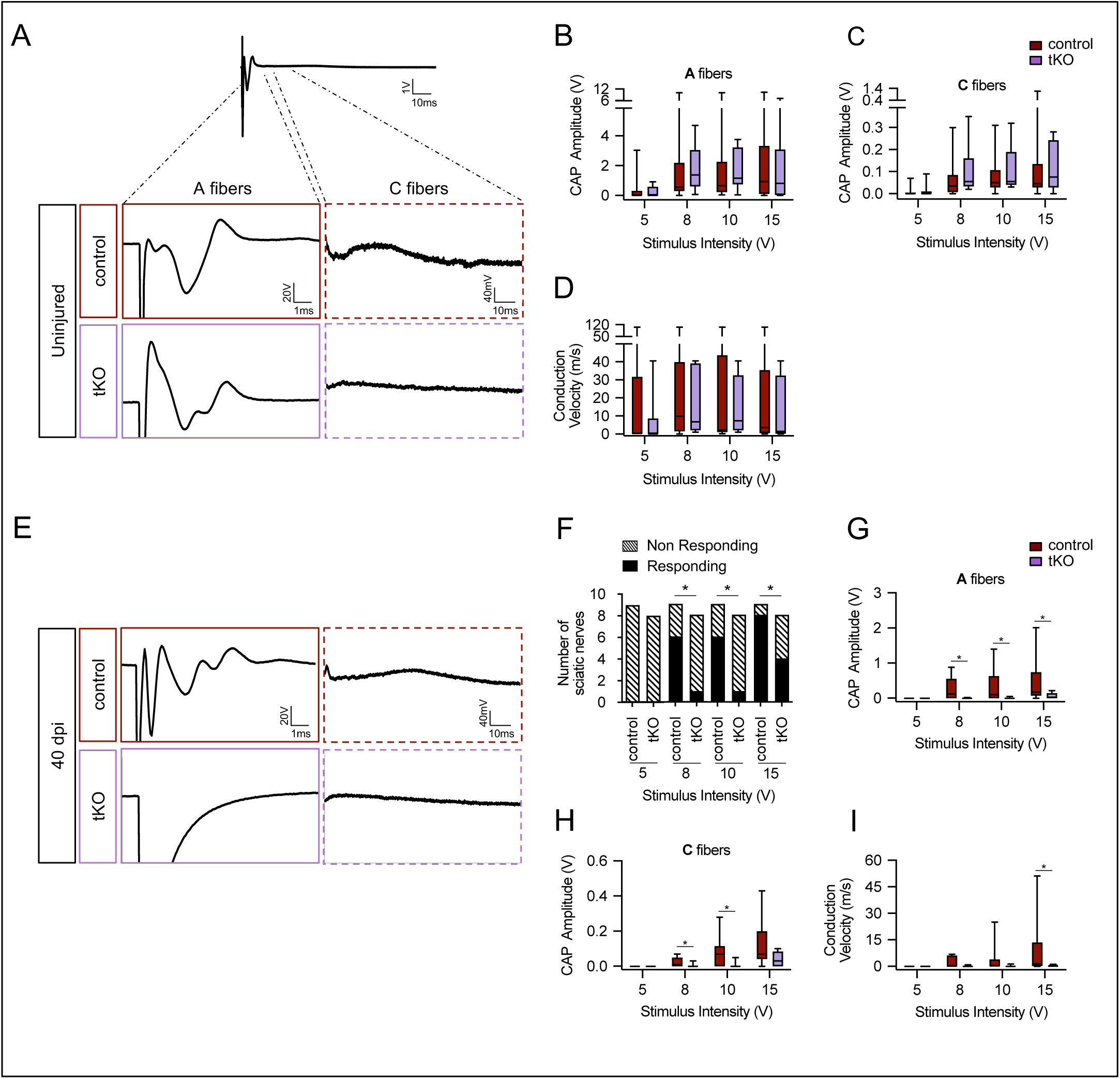
Remyelination failure in the *tKO* hampered nerve impulse conduction. **A)** Sample recordings of compound action potential (CAP) in uninjured sciatic nerves of control and *tKO* mice showing the waveform components corresponding to myelinated (A fibers) and unmyelinated (C) fibers. **B-C)** Waveform component of A fibers (**B**) and C-fibers (**C**) showed similar amplitude in both genotypes for stimulation with pulses of increasing intensity (5, 8, 10 and 15 V). **D)** Nerve conduction velocity was also preserved in the *tKO* mice. **E)** Sample recordings of CAPs obtained in a control and a *tKO* sciatic nerve 40 days after nerve crushing. **F)** Number of nerves that responded with a detectable CAP after the stimulation with increasing intensity. **G)** Amplitude of CAP corresponding to A fibers was significantly smaller in *tKO* than in control nerves for stimulation with 8, 10 and 15 V. **H)** Amplitude of C fiber component was significantly smaller in *tKO* than in control nerves for stimulation with 8 and 10 V. **I)** Nerve conduction velocity was significantly reduced after injury in both genotypes, being significantly lower in *tKO* than in control nerves for stimulation at 15 V. In this set of experiments, the whole length of a sciatic nerve was exposed from its proximal projection (L4 spinal cord) to its distal branches in deeply anesthetized mice. Compound action potentials (CAPs) were evoked by electrical stimulation of increasing intensity (5, 8, 10 and 15 V, 0.03 ms pulse duration). The maximum amplitude of the A-fiber and C-fiber components of CAP electrical signal, and their mean nerve conduction velocity were measured. Seven to 18 animals per genotype and condition were used. Mann-Whitney’s U were used for non-parametric paired comparisons and Chi-Squared test were used for statistical comparations. (* P<0,05; ** P<0,01; *** P<0,001; ns: no significant). See Source data section online (Graphs source data) for more details.

### Targets of class IIa HDACs in Schwann cells

To try to identify the genes regulated by class IIa HDACs we first performed a genome-wide transcriptomic analysis of the *tKO* remyelinating sciatic nerves and control littermates (Source data section online (RNAseq source data)). At 1 dpi, 395 genes were up regulated and 274 downregulated in the *tKO* (Figures S14A and S15A). Similar to the uninjured nerve analysis, the 10 most robustly changed genes were upregulated. Interestingly Tyrp1 and Mcam were also among the most upregulated genes. At 10 dpi the number of dysregulated genes was notably increased, with 1227 transcripts upregulated and 1550 downregulated (Figures S14B and S15A). Among the most robustly upregulated genes was the repair cell marker *Bdnf* (Arthur-Farraj et al., 2017; Kristjan R. Jessen & Mirsky, 2019), possibly reflecting impairment of the *tKO* repair Schwann cells to fully re-differentiate into myelin and non-myelin forming Schwann cells. Interestingly, in contrast with previous time points, 8 of the 10 most robustly changed genes were downregulated, including many myelin-related genes (*Mal*, *Prx*, *Mag*, *Ncmap*, *Pmp22*, *Mbp* and *Mpz*), which is expected given the dramatic delay in myelin sheath development in the *tKO* at 10 dpi (Figure 5N). At 20 dpi, we identified 1895 transcripts upregulated and 2450 downregulated in the tKO (Figure S14C and S15A). As before, the 10 most robustly changed genes were upregulated (Figure S14C) and among them we found again *Tyrp1*, *Mcam* and *Ednrb*. The negative regulator of myelination *c-Jun* was upregulated in the *tKO* from 1 dpi and remained increased up to 20 dpi (Figure S15B). A similar profile was shown for *Runx2, Gdnf*, *p75Ngfr* and *Sox2* (Figures S15C-F). *Oct-6* was induced up to 10 dpi in both control and *tKO* nerves, to later (20 dpi) downregulated in the control (20 dpi) but not in the *tKO* nerves (Figure S15G). By contrast, the master transcriptional regulator of myelination *Krox20,* was downregulated at 10 and 20 dpi in the *tKO*, when remyelination was highly active in the controls (Figure S15H). Probably as a consequence, early myelin genes such as *Drp2* and *Prx* were downregulated at different time points (Figures S15I and 15J). Other myelin protein genes that are expressed later, such as *Mpz*, *Mbp*, *Mag*, *Pmp22* and *Plp1* were also consistently downregulated (Figures S15K-O). Myelin is a specialized plasma membrane with a distinctive lipid composition particularly rich in cholesterol (Poitelon et al., 2020). During remyelination of the *tKO* nerves we found several genes of the sterol branch of the mevalonate pathway downregulated (*Hmgcs*, *Lss*, *Dhcr24*) (Figures S15P-R). We also found downregulated genes encoding for enzymes involved in the elongation (*Elov1*), transport (*Pmp2*) and insertion of double bonds (*Scd2, and Fads1*) into fatty acids (Figures S15S-V). Interestingly, *Cers2* and *Ugt8a* were also downregulated. These genes are involved in the synthesis of sphingomyelin and galactosyl- ceramide respectively (Figure S15W and 15X), both abundant lipids in myelin (Poitelon et al., 2020). Together our data shows that class IIa HDACs are necessary to block negative regulators of myelination and induce the expression of genes encoding for myelin proteins and key enzymes for the biosynthesis of myelin lipids.

To learn which genes are direct targets of class IIa HDACs and which are regulated indirectly, we performed a chromatin immunoprecipitation assay with anti-HDAC4 coupled to massive sequencing (ChIPseq) in dbcAMP differentiated Schwann cells. We found 3.932 peaks, 67,27% of which were located in the proximal promoter regions of genes (≤ 1Kb from the transcription start site (TSS)) (Figure 7A). The localization of these peaks in the rat genome is shown in the Source data section online (ChiPseq peaks source data). Importantly, ChIPseq analysis confirmed our previous results (Gomis-Coloma et al., 2018) showing that HDAC4 binds to the promoters of *c-Jun*, *Gdnf* and *Runx2* (Figure 7B-E). Interestingly, HDAC4 also binds to the promoter region of *Sox2* (Figure 7B and 7F), another negative regulator of myelination. We found also peaks for *Id2* and *Hey2* (Figure 7B and ChiPseq peaks source data). Here we show that *Oct-6* is overexpressed in the PNS of the *tKO* during development (Figure 1K-M), and that it is not properly downregulated during remyelination (Figure S15G). Interestingly, we found three peaks of HDAC4 bound near the TSS of *Oct-6* (Figures 7B and 7G), a result that was confirmed by ChIPqPCR (Figure 7H). Regarding the melanocyte lineage, we found a clear peak of HDAC4 close the TSS of *Mcam* (Figure 7B). However, we did not detect peaks in *Tyrp1* and *Ednrb*, suggesting that HDAC4 does not directly repress its expression, or alternatively it does it by using alternative promoters or enhancers. Surprisingly, we also found peaks in the promoter regions of *Mbp* and *Hmgcr* (Fig 7B), two genes highly expressed during myelination. Although it could seem contradictory (as HDACs have mainly a gene repressor activity), it is worth mentioning that HDACs are also bound to the promoter regions of highly expressed genes in other tissues (Wang et al., 2009) (see Discussion). Thus, our data shows that class IIa HDACs bind to and repress the expression of melanocyte lineage genes and negative regulators of myelination allowing myelination and remyelination proceed in a timely fashion.

**Figure 7.**
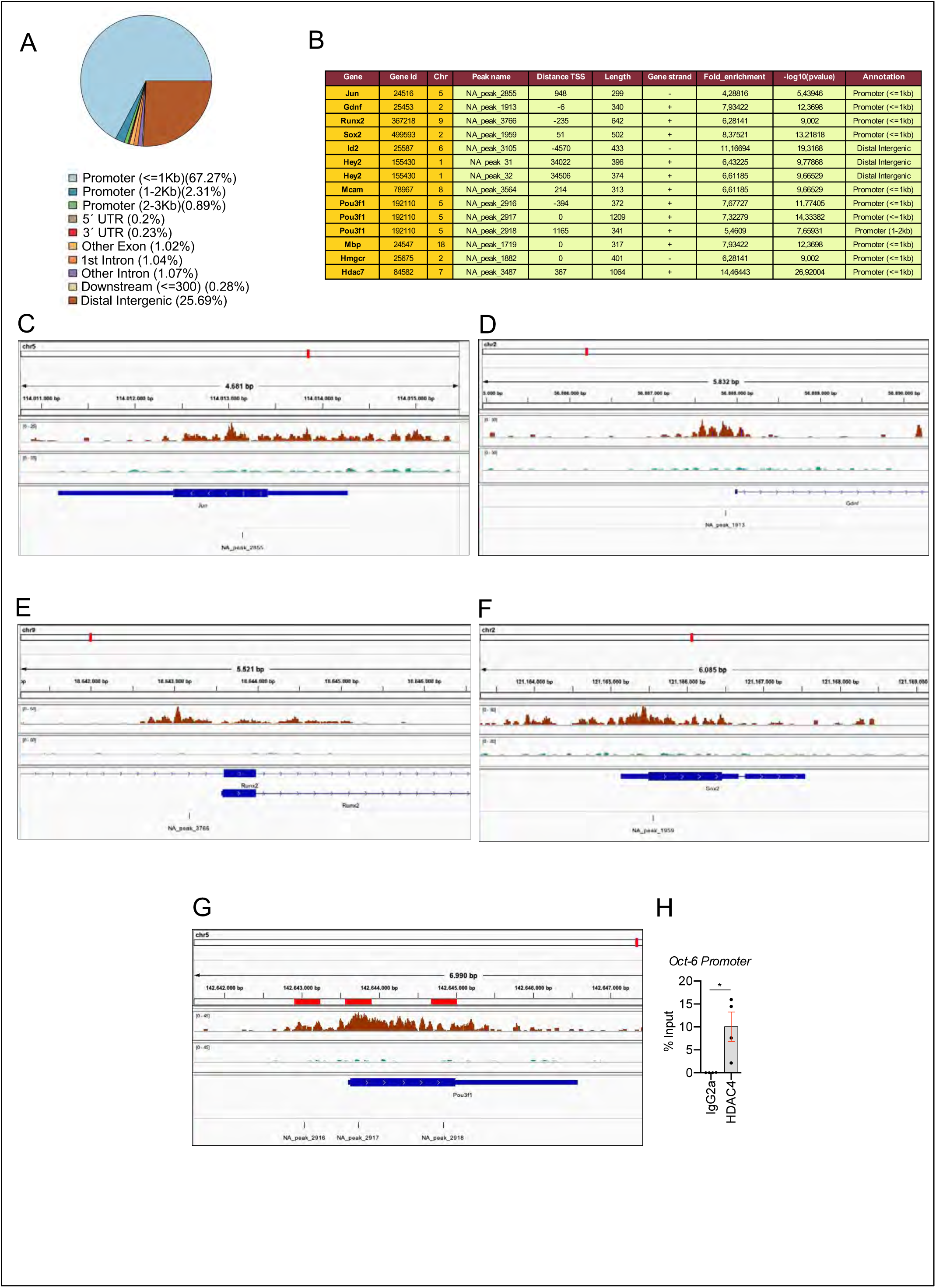
HDAC4 binds to the promoter of pivotal genes for myelin development. Cultured rat Schwann cells were incubated with 1 mM dbcAMP to shuttle HDAC4 into the nucleus, and ChIPseq analysis performed on the crosslinked chromatin, with anti-HADC4 antibody. **A)** 3.932 HDAC4 peaks were found in the rat genome. Most of these peaks (67,27%) were located in the promoter regions (≤ 1Kb from the TSS). **B)** A table with the localization of some of these peaks (a complete list can be found in See Source data section online (ChIPseq peaks source data)). **C-E**), ChIPseq signal analysis confirmed our previous results (Gomis-Coloma et al., 2018) showing that HDAC4 binds to the promoters of *c-Jun*, *Gdnf* and *Runx2.* **F)** HDAC4 was also found bound to the promoter region of *Sox2* . **G)** Three peaks (NA_peaks 2916,2917 and 2918) near the TSS of *Oct-6* were also found. **H)** The binding of HDAC4 to *Oct-6* gene was confirmed by ChIP-qPCR. 4 different experiments from 4 distinct cultures were used. Data were analyzed with the Mann-Whitney test.(* P<0,05; ** P<0,01; *** P<0,001; ns: no significant). See Source data section online (Graphs source data) for more details.

### c-Jun binds to the promoter and induces the expression of *HDAC7* gene in the PNS

As we have shown, the simultaneous elimination of HDAC4 and HDAC5 activates a mechanism to compensate for the drop in class IIa HDAC gene-dose in Schwann cells. This mechanism multiplies by 3-fold the expression of HDAC7, the other member of this family expressed in these cells (Figure 1A). But what mechanism is involved? We have shown before that c-Jun is highly expressed in the developing PNS of the *dKO* mice (Gomis-Coloma et al., 2018). Here we show that c-Jun expression remains high during remyelination in the *dKO* sciatic nerves (Figures 4L,4N and S15B). Interestingly, we found in ENCODE (https://www.encodeproject.org/) that c-Jun binds to the promoter region of *HDAC7* in A549 cells. Thus, the increased c-Jun might bind to the promoter of *HDAC7* inducing its compensatory overexpression in the *dKO* nerves. To test this, we checked whether c-Jun binds to the HDAC7 promoter in Schwann cells by ChIPqPCR. We found that in cultured Schwann cells c-Jun binds to the *HDAC7* promoter (Figure 8A). We then identified an evolutionarily conserved c-Jun consensus binding motif in the proximal promoter region of the mouse HDAC7 gene (Figure S16). To test this functionally, a fragment of 1.189 bp containing this region was amplified by PCR and cloned into the pGL3- luciferase reporter vector (see Materials and methods). The 1.189-promoter- HDAC7-pGL3 luciferase construct was transfected into HEK293 cells together with a pcDNA3 plasmid encoding for c-Jun, and luciferase activity measured 12h post-transfection. As shown in Figure 8B, this promoter fragment responded to c-Jun by increasing the luciferase activity by 3,4 fold over the control, which supports the idea that *HDAC7* expression is regulated by c-Jun. To further test this hypothesis *in vivo*, we utilized mouse transgenic lines (*P0Cre^+/-^ /R26 c-Jun ^stopflox/+^* mice) that either overexpresses *c-Jun* in Schwann cells, referred to as c-Jun OE mice (Fazal et al., 2017) or lack *c-Jun* expression in Schwann cells (*P0- Cre^+/-^ c-Jun^flox/flox^* mice ;Parkinson et al., 2008) (Figure 8C). c-Jun overexpression in Schwann cells induced *HDAC7* expression by almost two-fold (Figure 8D). This was a specific effect as no changes were found for *HDAC4* and *HDAC5* mRNA (Figures 8E and F). By contrast, *c-Jun* removal produced no changes in the expression of any class IIa HDACs, suggesting it is not necessary for the basal expression of these HDACs (Figures 8D-F). Thus, our data clearly show that c-Jun induces the expression of HDAC7 by Schwann cells *in vivo.* Interestingly, we detected a peak of HDAC4 bound to the promoter of HDAC7 (Figures 7B and 8G) in the ChIPseq experiment, a result that we confirmed by ChIPqPCR (Figure 8H). This suggests that, in differentiated Schwann cells, other class IIa HDACs contribute to maintaining HDAC7 expression at the basal level. The simultaneous loss of HDAC4 and HDAC5 allows the expression of c-Jun, which can bind to the promoter of HDAC7, now free of the repression by class IIa HDACs, increasing the transcription of this deacetylase.

**Figure 8.**
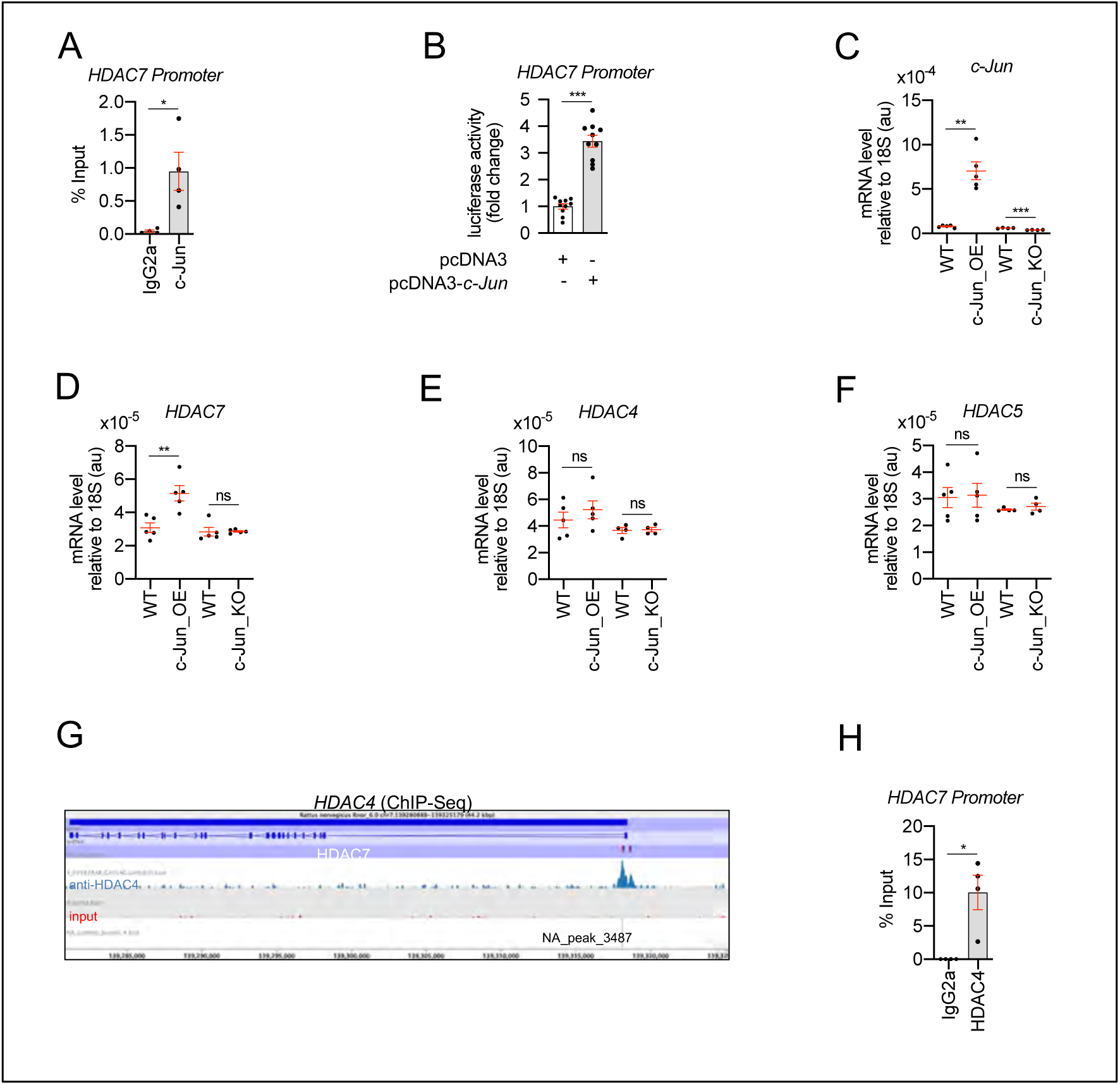
HDAC7 compensatory overexpression is induced by c-Jun. **A)** ChIPqPCR of dbcAMP treated rat Schwann cells with anti-c-Jun antibody showed that this transcription factor is bound to the promoter of HDAC7. 4 different experiments from 4 distinct cultures were used. Data were analyzed with the Mann-Whitney test. **B)** A 1.189 fragment of the mouse HDAC7 promoter containing a conserved c-Jun binding consensus sequence was PCR amplified and cloned into the pGL3 luciferase reporter vector. HEK293 cells were transfected with this construct and the pcDNA3 (empty vector) or pcDNA3 c-Jun. As is shown c-Jun induced the luciferase activity by 3,4 ± 0,22 fold (P<0,0001; n=10). Unpaired t-test with Welch’s correlation was used for statistical comparison. **C)** Levels of mRNA for c-Jun in the nerves of c-Jun_OE and c-Jun_ KO mice. **D)** HDAC7 expression is enhanced in the sciatic nerves of the c- Jun_OE mice (5,16 ± 0,46 x10^-5^ in the c-Jun_OE versus 3,09 ± 0,29 x10^-5^ in the WT; P=0,005) but doesn’t change in the c-Jun_ KO mice. **E)** HDAC4 expression in sciatic nerves doesn’t change in c-Jun_OE and c-Jun_ KO mice. **F)** HDAC5 expression in sciatic nerves doesn’t change in c-Jun_OE and c-Jun_ KO mice. RTqPCR with mouse-specific primers for the indicated genes was performed. The scatter plot, which include also the mean ± SE, shows the expression of each gene normalized to the housekeeping 18S. 4 to 5 mice per genotype were used. Data were analyzed with the unpaired t-test with Welch’s correlation. **G)** A peak of HDAC4 (NA_peak 3487) was found on the HDAC7 promoter in the ChIPseq experiemt. **H)** ChIPqPCR confirmed that HDAC4 is bound to the promoter of HDAC7. 4 different experiments from 4 distinct cultures were used. Data were analyzed with the Mann-Whitney test. (* P<0,05; ** P<0,01; *** P<0,001; ns: no significant). See Source data section online (Graphs source data) for more details.

### HDAC9 is expressed *de novo* in the sciatic nerve of the *tKO*

Although developmental myelination and remyelination after injury were notably delayed in the *tKO* nerves, Schwann cells in these mice are still able to eventually myelinate axons. One possibility could be that HDAC9, the final class IIa HDAC left to be tested, was responsible for the rescue of this phenotype. However we have consistently found extremely low or undetectable levels of the mRNA for this protein in the C57B6 mouse sciatic nerves and cultured rat Schwann cells (Gomis-Coloma et al., 2018), a result that is in agreement with the recent Sciatic Nerve ATlas (SNAT: https://www.snat.ethz.ch) (Gerber et al., 2021). Surprisingly, we found HDAC9 among the most robustly upregulated genes by the *tKO* nerves in the RNAseq analysis (Figure 2H). To confirm this result, we measured the mRNA for this gene in the sciatic nerves of the *tKO* and control adult mice (P60) by RTqPCR. As is shown in Figure 9A, the mRNA for HDAC9 was increased by 4,4-fold in the *tKO* mice (P<0,0001). This increase was not found in the *dKO* and was smaller in the *cKO7* (2,4 fold; P=0,0057) and *cKO4* (1,3 fold; P=0,01) mice. These results suggest that HDAC9 is strongly expressed in the sciatic nerves of the *tKO* mice to compensate for the absence of other class IIa HDACs. Indeed, we found that HDAC9 is already induced early in development in the *tKO* (P2) whereas HDAC9 mRNA was practically undetectable in the nerves of control animals (Figure 9B). At this time, myelin sheaths are present, although sparse, in mutant nerves (Figure 1B-J). At P8, HDAC9 expression remains extremely low in control nerves, but it is robustly expressed in the *tKO* mice (an increase of 5,3- fold; P=0,0117), correlating with an increase in the number of myelinated axons (Figure 1B-J). Robust HDAC9 gene expression was maintained in the sciatic nerve of the *tKO* at P60 (Figure 9A), when myelination catches up with levels in control nerves. Interestingly, we found that in the *tKO* nerves, the lysine 9 of histone 3 on the HDAC9 promoter is more acetylated (Figure 9C), suggesting that the absence of class IIa HDACs allows the expression of this gene. Importantly, HDAC9 gene is furtherly upregulated in the injured sciatic nerves of *tKO* mice, as is shown in Figure 9D, where we show the alignment of the reads of the RNAseq from three individual sciatic nerves from control and *tKO* mice at 20 days post crush (20 dpi), and in Figure 9E, where followed the mRNA levels of HDAC9 (as Fragments Per Kilobase Million, FPKMs) at 0, 1, 10 and 20 days post crush in the RNAseq experiment. Thus far, our data suggest that in the absence of class IIa HDACs, repression is lost and Schwann cells start to express *de novo* HDAC9.

**Figure 9.**
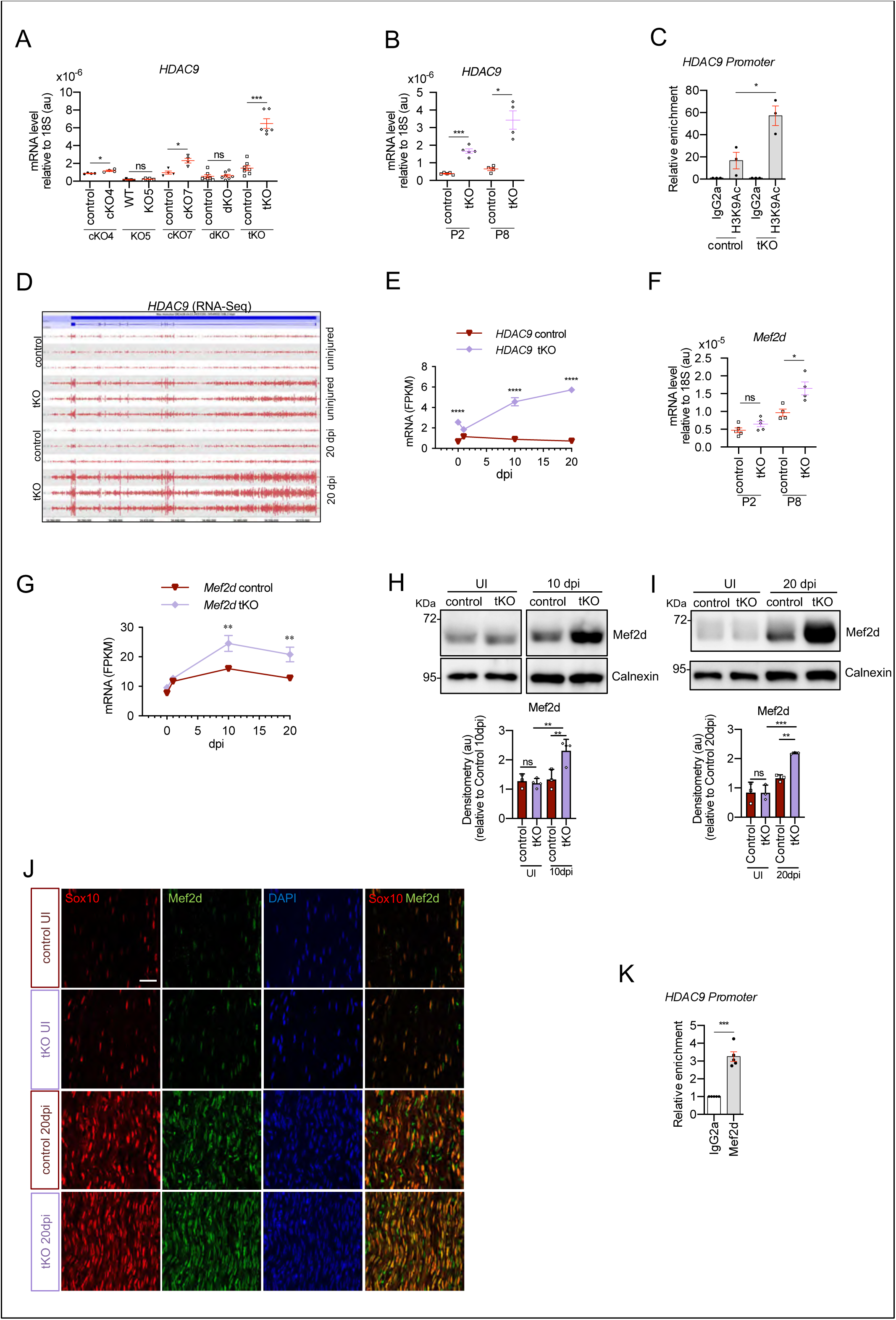
Mef2d mediates HDAC9 *de novo* expression in the tKO. **A)** HDAC9 expression is notably induced in the sciatic nerves of the adult (P60) *tKO* mice (6,48 ± 0,53 x10^-6^ au the *tKO* versus 1,46 ± 0,28 x10^-6^ in the control; P<0,0001). Only minor changes were observed in the *cKO4* and *cKO7*. 4 to 8 mice per genotype were used. Unpaired t-test was used for comparations. **B)** HDAC9 expression is increased from early postnatal development of the nerve. At P2 we found 1,67 ± 0,13 x10^-6^ au in the *tKO* versus 0,39 ± 0,03 x10^-6^ in the controls (P<0,0001) and at P8 we found 3,43 ± 0,52 x10^-6^ au in the *tKO* versus 0,65 ± 0,09 x10^-6^ in the controls (P<0,0001). RTqPCR with mouse-specific primers for HDAC9 was performed. The scatter plot, which include also the mean ± SE, shows the expression of HDAC9 normalized to the housekeeping 18S. 4 to 5 mice per genotype were used. Data were analyzed with the unpaired t-test with Welch’s correlation. **C)** ChIPqPCR with anti-H3K9Ac of adult (P60) sciatic nerves of *tKO* and control mice. Three different experiments of 4-5 animals per genotype is shown. Data were normalized to the IgG value as shown as relative enrichment. Unpaired t-test was used for comparations. **D)** Alignment of the reads of the RNAseq from 3 individual sciatic nerves of control and 3 *tKO* mice, both uninjured and at 20 days post crush (20 dpi). HDAC9 gene is transcribed at detectable levels in the sciatic nerve of the uninjured *tKO* mice, whereas it is almost nondetectable in the control sciatic nerves. The *tKO* mice (but not the controls) increase additionally the expression of HDAC9 gene during remyelination (20 dpi). **E)** mRNA levels of HDAC9 (as FPKMs) at 0, 1, 10 and 20 days post crush (dpi) in the RNAseq experiment. Two-way ANOVA was used for statistical comparation. **F)** Mef2d expression is increased early in development (P8) in *tKO* nerve (1,65 ± 0,18 in the *tKO* versus 0,97 ± 0,10 in controls; P=0,025). RTqPCR with mouse-specific primers for Mef2d was performed. The scatter plot, which include also the mean ± SE, shows the expression normalized to the housekeeping 18S. 4 to 5 mice per genotype were used. Data were analyzed with the unpaired t-test with Welch’s correlation. **G)** mRNA levels of Mef2d (as FPKMs) at 0, 1, 10 and 20 days post crush (dpi) in the RNAseq experiment. **H)** A representative WB of protein extracts from *tKO*, control and wild type nerves at 10 dpi is shown. In the quantification, Mef2d protein was increased in the *tKO* nerves (2,31 ± 0,19 au in the *tKO* versus 1,33 ± 0,19 in controls; P<0,0069) (**I**) Same for 20 dpi (2,19 ± 0,03 au in the *tKO* versus 1,33 ± 0,11 in controls; P<0,0073). Densitometric analysis was done for 3 to 4 WB from the same number of mice and normalized to the control 20 dpi. Data were analyzed with the unpaired t-test. **J)** Mef2d colocalizes with the transcription factor Sox10^+^, suggesting that it is expressed by Schwann cells. P60 sciatic nerves were fixed and submitted to immunofluorescence with the indicated antibodies. Nuclei were counterstained with Hoechst. Representative confocal images of sections obtained from the sciatic nerves of control and *tKO* mice are shown. Scale bars= 20 μm. **K)** Mef2d binds to the HDAC9 promoter in the *tKO*. ChIPqPCR of 20 dpi nerves of *tKO* mice was performed using an antiMef2d specific antibody. 5 different experiments from 4 to 5 mice per genotype were performed. Data were analyzed with the unpaired t-test. (* P<0,05; ** P<0,01; *** P<0,001; ns: no significant). See Source data section online (Graphs source data) for more details.

To investigate which transcription factor is responsible of HDAC9 expression in the *tKO* we again looked at c-Jun, however we couldn’t detect c-Jun bound to the promoter region of HDAC9 in A549 cells in the ENCODE database. Also, HDAC9 mRNA levels were not increased in the sciatic nerves of the c-Jun OE mice (Figure S17). Interestingly, it has been described that Mef2d binds to the promoter and regulates the expression of HDAC9 during muscle differentiation, and in leiomyosarcoma cells (Di Giorgio et al., 2020; Haberland et al., 2007). We therefore reasoned that Mef2d might be involved in the upregulation of HDAC9 in Schwann cells. Indeed, we have previously shown that Mef2 family members are expressed in cultured Schwann cells and adult sciatic nerves (Gomis-Coloma et al., 2018). Importantly, RTqPCR shows that *Mef2d* mRNA is induced in the nerves of the *tKO* during development (Figure 9F). Also, RNAseq data showed that *Mef2d* is upregulated in the sciatic nerves of the *tKO* mice at 10 and 20 dpi (Figure 9G). Western blot supported this idea (Figures 9H and 9I) and immunofluorescence studies (Figure 9J) showed that Mef2d is expressed by Schwann cells (Sox10^+^). Importantly, we detected Mef2d bound to the promoter of HDAC9 in the *tKO* nerves at 20 dpi (Figure 9K). Altogether these data support the view that in the *tKO* nerve, Mef2d is induced to activate the *de novo* expression of *HDAC9* and keep a sufficient class IIa HDAC gene dose to allow myelin formation during development and after nerve injury.

## DISCUSSION

Functional redundancies, consequence of gene duplications, are found in most genomes, and are postulated to give a robustness to organisms against mutations (Kafri et al., 2006; Peng, 2019). It has been predicted that redundancy is evolutionarily unstable and has only a transient lifetime. However, numerous examples exist of gene redundancies that have been conserved throughout dilated evolutionary periods. Interestingly, many of these redundant genes are transcriptionally regulated in response to the intactness of their partners, being up-regulated if the latter are inactivated, and it has been proposed that they are evolutionary stable because they reduce the harmful effects of gene expression noise (Kafri et al., 2006). Redundancies have been shown to be particularly relevant during development. Although in many cases redundant genes have different spatial and/or temporal expression patterns, some degree of overlap is usually observed. One example of developmental regulators that have preserved redundancy in spite of the evolutionary distance is the couple *MyoD/Myf-5*, which are master regulators of skeletal muscle development (Sabourin & Rudnicki, 2000). Similar to what happens with other redundant couples, *Myf-5* expression has a linear response that strictly dependents on the *MyoD* gene expression dosage, which can likely contribute to reduced gene expression noise and thus allow skeletal muscle differentiation (Kafri et al., 2006). Similarly, slow oxidative fiber gene expression in skeletal muscles depends on class IIa HDACs gene dosage (Potthoff, Wu, et al., 2007). Here we show that the activation of the myelin gene expression program by Schwann cells is also directly related to class IIa HDACs gene dosage. Thus, differentiation is directly related to class IIa HDAC gene dose in at least two different tissues. But how is this process regulated? Despite genetic compensation being documented many times in different organisms, our understanding of the underlying molecular mechanisms that control this process still remains limited (El-Brolosy & Stainier, 2017). Thus, genetic compensation of class I HDACs has been previously described during myelin development (Jacob et al., 2011), however the mechanisms regulating this process have not been investigated. Here we show that removal of HDAC4 and HDAC5 upregulates the compensatory expression of HDAC7 in Schwann cells allowing, although with delay, myelin formation both during development and after nerve injury. Our data suggest that it is mediated by the transcription factor *c-Jun* which binds to and induce the expression of HDAC7 gene. Interestingly, we found HDAC4 is bound to the promoter of HDAC7 in differentiated Schwann cells, suggesting other class IIa HADCs contribute to maintain the expression of this gene at basal level. In the *dKO*, the absence of HDAC4 and HADC5 might allow c*-Jun* to bind and stimulate the compensatory expression of HDAC7 (Figure 10).

**Figure 10.**
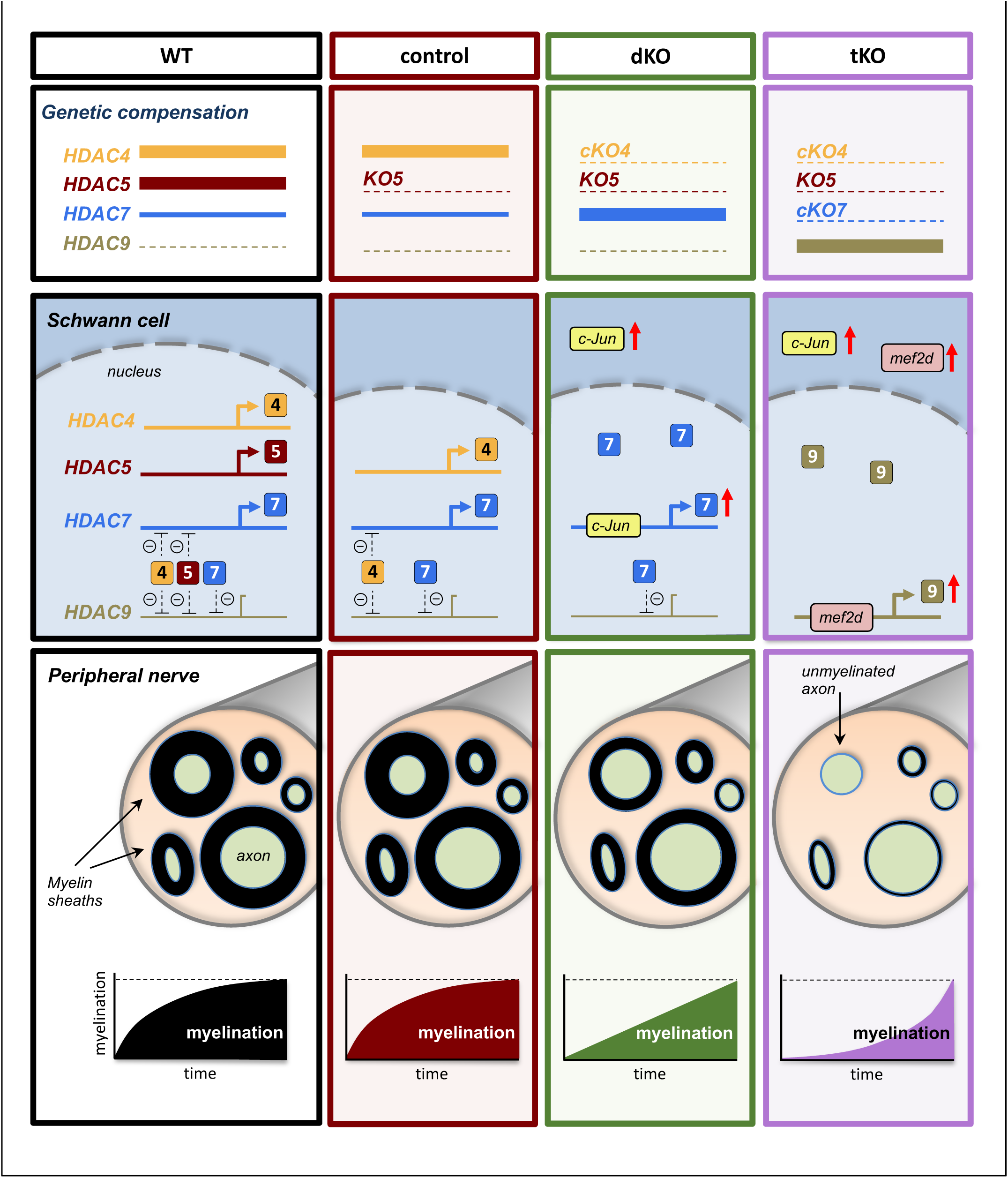
A graphical summary of the proposed model: In wild type nerves (WT) the expression of HDAC4, HDAC5 and HDAC7 allows myelin formation and blocks the expression of HDAC9. The removal of HDAC5 (control) has no effects on myelination neither HDAC9 expression. In the nerves of the *dKO,* c-Jun induces the compensatory overexpression of HDAC7, allowing delayed myelination but having no effect on HDAC9 gene expression. The simultaneous elimination of HDAC4, HDAC5 and HDAC7 (*tKO*) induces the overexpression of Mef2d, which binds to the promoter and induce the compensatory expression of HDAC9, which after a long delay induces myelination.

Developmental myelination and remyelination in *tKO* nerves are notably delayed, although not completely blocked. One possibility is that HDAC9, another deacetylase of the same family, could compensate during myelination. However our data and data from other labs (Gerber et al., 2021), show that this gene is not expressed in peripheral nerves. Strikingly, we found HDAC9 is one of the most robustly upregulated genes in the nerves of the *tKO*. This supports that HDAC9 gene is *de novo* expressed in response to the drop of class IIa HDAC gene dose. To investigate mechanisms that activate *de novo* expression of *HDAC9* in the *tKO* nerves we focused our attention in the Mef2 family of transcription factors, as it is known they regulate *HDAC9* expression in other tissues (Di Giorgio et al., 2020; Haberland et al., 2007). Interestingly, we found *Mef2d* overexpressed in the *tKO* Schwann cells, both during development as well as during remyelination after nerve crush. Class IIa HDACs can associate with Mef2 and suppress its transcriptional activity (Haberland et al., 2007). Thus, repressor activity of HDAC4, HDAC5 and/or HDAC7 could theoretically block the expression of HDAC9 in control nerves. By contrast, in the *tKO* Schwann cells, HDAC9 promoter will be free of repression as no class IIa HDAC is expressed in these cells. In support of this, we found much more H3K9Ac associated with HDAC9 in *tKO* nerves. Importantly, we also detect Mef2d bound to the promoter of HDAC9 in the nerves of the *tKO* at 20 dpi. Taken together, our data suggest that the overexpressed and non-repressed Mef2d transcription factor is able to bind to the promoter and induce the *de novo* expression of HDAC9 in the Schwann cells of the *tKO* mice (Figure 10).

Although adult *tKO* nerves have morphologically normal myelin, RNAseq analysis showed 1270 genes differently expressed, the most robustly changed genes being upregulated. This suggests a predominantly gene repressive function for class IIa HDACs in Schwann cells, as is the case for other cell types (Parra, 2015; Parra & Verdin, 2010). The most robustly upregulated gene in these mice was *Tyrp1*, a gene involved in the stabilization of tyrosinase and the synthesis of melanin. Strikingly, we did not find expression of the tyrosinase gene in these nerves, suggesting a different role for Tyrp1. In fact, our data shows that Tyrp1 mRNA is not translated into protein (Figure S6D). It has been shown that Tyrp1 mRNA indirectly promotes melanoma cell proliferation by sequestering miR-16 (Gautron et al., 2020; Gilot et al., 2017). Whether *Tyrp1* mRNA is also responsible for the increased cell proliferation of Schwann cells in the *tKO* nerve needs to be clarified. Because a subgroup of melanocytes are formed from Schwann cell precursor (SCP) cells (Adameyko et al., 2009) and *P0-Cre* is already expressed by SCPs, our data strongly suggest that class IIa HDACs are necessary to repress genes involved in SCP proliferation and differentiation into the melanocytic lineage.

We show that class IIa HDACs removal delays remyelination after nerve crush. Importantly, myelin clearance is not hampered which rules out a problem in the removal of debris after nerve injury as the cause for the remyelination delay. Indeed, *tKO* Schwann cells are efficiently reprogramed into the repair phenotype and myelin clearance is even more efficient than in control nerves. Although we don’t know why myelin clearance is accelerated, it is worth to mention that the expression of repair Schwann cell markers is increased in *tKO* nerves, what could improve reprograming into the repair phenotype. Future experiments are needed to clarify this point.

Genome-wide transcriptomic analysis showed that the number of differentially expressed genes increases after crush, and is maximum at 20 dpi. Many genes are robustly upregulated, particularly at 1 and 20 dpi, supporting further the idea that the main role of class IIa HDACs in Schwann cells is to repress gene expression. This agrees with the role of this family of deacetylases in other contexts (Chang et al., 2004, 2006; Parra & Verdin, 2010) where they work mainly as co-repressors of transcription factors such as Mef2 and Runx2 (Bialek et al., 2004; Potthoff & Olson, 2007; Vega et al., 2004). Interestingly, among the upregulated genes in the *tKO* injured nerves we found *c-Jun*, *Runx2*, *Gdnf*, *p75Ngfr* and *Sox2*, all expressed by non-myelinating and repair Schwann cells. We also found *Oct-6* overexpressed in the *tKO*. It has been shown that *Oct-6* overexpression blocks the transition of Schwann cells from the promyelinating into the myelinating stage (Jaegle et al., 1996). Thus, *Oct-6* mis-expression may contribute to the delayed myelination of the *tKO* nerve both during development and after injury. In a simplistic model, the failure of class IIa HDACs to downregulate *c-Jun* could indirectly induce the expression of other negative regulators of myelination controlled by this transcription factor. However, class IIa HDACs could also directly repress the expression of other negative regulators of myelination. To explore this idea, we performed a genome-wide mapping of genes that are direct targets of HDAC4. Importantly, we confirmed our previous results showing that HDAC4 binds to the promoters of *c-Jun*, *Runx*2 and *Gdnf* in Schwann cells (Gomis-Coloma et al., 2018). We found that HDAC4 also binds to the promoters of many other genes including other negative regulators of myelination such as *Sox2*, *Id2* and *Hey2*. Interestingly, HDAC4 also binds to the promoter of *Oct-6*. Thus, the direct repressive effect of class IIa HDACs is not circumscribed to *c-Jun* but is much wider, supporting the view that they work as a cAMP-regulated blocking hub for repressors of myelination.

Surprisingly, we found that HDAC4 is also bound to the promoter of *Mbp* and *Hmgcr*, two genes that are actively expressed during myelination. HDACs are usually bound to repressed genes and replaced by histone acetyl transferases (HATs) upon gene activation (Lan et al., 1999). However, it has been shown that class I HDACs are also bound, together with HAT, to the promoter regions of actively transcribed genes (Wang et al., 2009). Interestingly, the histones associated with these promoters are heavily acetylated, what makes these authors to propose that one function of HDACs is to remove the acetyl groups added by HATs in active genes to reset chromatin modification after gene activation. Although class IIa HDACs have no deacetylase activity, they recruit class I HDACs by forming a complex with NcoR1 and SMRT. Thus, the possibility exists that class IIa and class I HADCs have a similar role when bound to the promoter of highly active genes in myelinating Schwann cells.

In summary, the data presented in this manuscript unveil responsive back-up circuits mediated by the transcription factors c-Jun and Mef2d, that coordinate genetic compensatory mechanisms within class IIa HDACs, aimed at repressing the expression of negative regulators of myelination to ensure differentiation of Schwann cells in response to cAMP, and the generation of the myelin sheath during development and after nerve injury.

## MATERIALS AND METHODS

### Animal studies

All animal work was conducted according to European Union guidelines and with protocols approved by the Comité de Bioética y Bioseguridad del Instituto de Neurociencias de Alicante, Universidad Hernández de Elche and Consejo Superior de Investigaciones Científicas (http://in.umh.es/). To avoid suffering, animals were sacrificed by cervical dislocation. The *P0-cre* mouse line is described in Feltri et al., 1999. *P0-cre*^-/-^ littermates were used as controls. HDAC4 floxed mice are described in Lehmann et al., 2018 and Potthoff et al., 2007. The HDAC5 KO mouse line is described in Chang et al., 2004. HDAC7 floxed mice are described in (Chang et al., 2006). The c-Jun_OE and c-Jun_KO mouse lines are described in Fazal et al., 2017 and Parkinson et al., 2008. MGI ID can be found in Source data section online (Key Resources Table). Experiments used mice of either sex on the C57BL/6 background.

### Plasmids

The luciferase reporter plasmid was generated by cloning the mouse HDAC7 promoter into the NheI site of pGL4 Luciferase reporter plasmid (Promega). The mouse HDAC7 promoter was amplified using Platinum SuperFi II DNA Polymerase (12361010, Thermo Fisher Scientific) and primers described in Source data section online (Key Resources Table). The plasmid pCMV-c-Jun was a gift of Dr Marta Giralt (Universitat de Barcelona).

### Reporter activity assays

HEK293 cells were transfected with the indicated constructs and then lysed. Their luciferase activity was determined with the Luciferase Assay System (Promega) using the manufacturer’s recommendations.

### Cell cultures

Schwann cells were cultured from sciatic nerves of neonatal rats as described previously (Brockes et al., 1979) with minor modifications. We used P3-P4 Wistar rat pups. The sciatic nerves were cut out from just below the dorsal root ganglia and at the knee area. During the extraction and cleaning, the nerves were introduced into a 35-mm cell culture dish containing 2 ml of cold Leibovitz’s F-15 medium (Gibco) placed on ice. The nerves were cleaned, desheathed, and placed in a new 35-mm cell culture dish containing DMEM GlutaMAX, 4.5 g/l glucose (Gibco) with 1 mg/ml of collagenase A (Roche). Subsequently they were cut into very small pieces using a scalpel and left in the incubator for 2 h. Nerve pieces were homogenized using a 1 ml pipette, digestion reaction stopped with complete medium, and the homogenate poured through a 40-μm Falcon Cell Strainer (Thermo Fisher Scientific). We then centrifuged the homogenate at 210 g for 10 min at room temperature and resuspended the pellet in complete medium supplemented with 10 μM of cytosine- β-D-arabinofuranoside (Sigma-Aldrich) to prevent fibroblast growth. The resuspended cells were then introduced into the polyl-lysine–coated 35-mm cell culture dishes. After 72 h, the medium was removed and cell cultures expanded in DMEM supplemented with 3% FBS, 5 μM forskolin, and 10 ng/ml recombinant NRG1 (R&D Systems). Where indicated, cells were incubated in SATO medium (composed of a 1:1 mixture of DMEM and Ham’s F12 medium (Gibco) supplemented with ITS (1:100; Gibco), 0.1 mM putrescine, and 20 nM of progesterone (Bottenstein & Sato, 1979). HEK 293 cells were obtained from Sigma-Aldrich. The cells were grown in non-coated flasks with DMEM GlutaMAX, 4.5 g/l glucose (Gibco) supplemented with 100 U/ml penicillin, 100 U/ml streptomycin, and 10% bovine fetal serum. Cells were transfected with plasmid DNA using Lipofectamine 2000 (Thermo Fisher Scientific) following the manufacturer’s recommendations.

### Nerve injury

Mice were anesthetized with 2 % isoflurane, and the sciatic nerve was exposed and cut or crushed (3 x 15 s at 3 rotation angles) at the sciatic notch using angled forceps. The wound was closed using veterinary autoclips (AutoClip® System). The nerve distal to the cut or crush was excised for analysis at various time points. Contralateral uninjured sciatic nerves were used as controls. For Western blotting and mRNA extraction we used the first 8 mm of the distal stump of injured nerves, and 1 cm or 3 cm of the control nerve, respectively. For electron microscopy we used the first 5,5 mm of the distal stump of injured nerves.

### mRNA detection and quantification by RT-qPCR

Total mRNA from uninjured or injured sciatic nerves was extracted using used TRI reagent / chloroform (Sigma-Aldrich) and the mRNA was purified using a NucleoSpin RNA mini kit (Macherey-Nagel), following manufacturer’s recommendations. RNA quality and concentration were determined using a nanodrop 2000 machine (Thermo). Genomic DNA was removed by incubation with RNase free DNase I (Thermo Fisher Scientific), and 500 ng RNA was primed with random hexamers (Invitrogen) and retrotranscribed to cDNA with Super Script II Reverse transcriptase (Invitrogen). Control reactions were performed omitting retrotranscriptase. qPCR was performed using an Applied Biosystems QuantStudio 3 Real Time PCR System and 5× PyroTaq EvaGreen qPCR Mix Plus (CMB). To avoid genomic amplification, PCR primers were designed to fall into separate exons flanking a large intron wherever possible. A list of the primers used can be found in Source data section online (Key Resources Table). Reactions were performed in duplicates of three different dilutions, and threshold cycle values were normalized to the housekeeping gene 18S. The specificity of the products was determined by melting curve analysis. The ratio of the relative expression for each gene to 18S was calculated by using the 2^-ΔΔCT^ formula. Amplicons were of similar size (≈100 bp) and melting points (≈85°C). Amplification efficiency for each product was confirmed by using duplicates of three dilutions for each sample.

### RNA sequencing analysis

Total RNA was isolated using the NucleoSpin RNA, Mini kit for RNA purification (Macherey-Nagel). The purified mRNA was fragmented and primed with random hexamers. Strand-specific first-strand cDNA was generated using reverse transcriptase in the presence of actinomycin D. The second cDNA strand was synthesized using dUTP in place of dTTP to mark the second strand. The resultant cDNA was then “A-tailed” at the 3end to prevent self-ligation and adapter dimerization. Truncated adaptors containing a T overhang were ligated to the A-tailed cDNA. Successfully ligated cDNA molecules were then enriched with limited cycle PCR (10–14 cycles). Libraries to be multiplexed in the same run were pooled in equimolar quantities. Samples were sequenced on the NextSeq 500 instrument (Illumina). Run data were demultiplexed and converted to fastq files using Illumina’s bcl2fastq Conversion Software version 2.18 on BaseSpace. Fastq files were aligned to the reference genome (Mouse (GRCm38/Ensembl release 95) and analyzed with Artificial Intelligence RNA-SEQ (A.I.R) software from Sequentia Biotech (https://www.sequentiabiotech.com/).

### Antibodies

Immunofluorescence antibodies: c-Jun (Cell Signaling Technology, rabbit 1:800), Ki67 (Abcam, rabbit 1:100), L1 (Chemicon International, rat 1:50), Mcam (Origene, rabbit 1:200), Mpz (AvesLab, chicken 1:1000), p75NGFR (Thermo Fisher Scientific, mouse 1:100), Sox10 (R and D Systems, goat 1:100), donkey anti-goat IgG (H+L) Alexa Fluor 555 conjugate (Molecular Probes, 1:1000), donkey anti-rabbit IgG (H+L) Alexa Fluor 488 conjugate (Molecular Probes, 1:1000), donkey anti-chicken IgY (H+L) Alexa Fluor 488 conjugate (Jackson ImmunoResearch Labs, 1:1000), goat anti-rat IgG (H+L) Alexa Fluor 555 conjugate (Molecular Probes, 1:1000), Cy3 donkey anti-mouse IgG (H+L) (Jackson Immunoresearch, 1:500) and Cy3 donkey anti-rabbit IgG (H+L) (Jackson Immunoresearch, 1:500).

Antibodies used for western blotting: calnexin (Enzo Life Sciences, rabbit 1:1000), c-Jun (Cell Signaling Technology, rabbit 1:1000), GAPDH (Sigma-Aldrich, rabbit 1:5000), HDAC5 (Santa Cruz, mouse 1:500), Krox-20 (Millipore, rabbit 1:500), Mcam (Origene, rabbit 1:1000), Mpz (AvesLab, chicken 1:1000), p75NGFR (Covance, rabbit 1:1000), Tyrp1 (Sigma-Aldrich, rabbit 1:1000), IgY anti-chicken HRP-linked (Sigma-Aldrich, 1:2000), IgG anti-mouse and IgG anti-rabbit HRP-linked (Cell Signaling Technology, 1:2000). A list of the antibodies used can be found in Source data section online (Key Resources Table).

### Immunofluorescence

For immunofluorescence, mice were sacrificed by cervical dislocation and fresh frozen tissue was embedded in OCT (Sakura, 4583). Cryosections were cut at 10 μm on Superfrost Plus slides (Thermo Scientific, J1800AMNZ). Sections were thawed and fixed with 4% PFA for 5 min at room temperature. Then, samples were washed 3x in PBS 1x and immersed in 50% acetone, 100% acetone and 50% acetone for 2 min each. Then samples were washed 3x in PBS 1x and blocked in 5% donkey serum 0.1% BSA in PBS for 1h. Samples were incubated with the appropriate primary antibodies diluted in blocking solution overnight at room temperature. A list of the antibodies used can be found in Source data section online (Key Resources Table). Samples were washed and incubated with secondary antibodies and DAPI in blocking solution for 1h at room temperature. Samples were mounted in Fluoromont G. Images were obtained at room temperature using a confocal ultraspectral microscope (Vertical Confocal Microscope Leica SPEII) with a 63× Leica objective and using Leica LAS X software. Images were analyzed with ImageJ software.

### EM studies

Mice were sacrificed by cervical dislocation and sciatic nerve were exposed and fixed by adding fixative solution (2% PFA (15710, Electron Microscopy Sciences), 2.5% Glutaraldehyde (16220, Electron Microscopy Sciences), 0.1M Cacodylate buffer pH=7.3 (12300, Electron Microscopy Sciences) for 15 minutes. Afterwards, the nerve was removed and placed in same fixative solution overnight at 4°C. Then, the nerve was washed in 0.1M cacodylate buffer 3x for 15 minutes each. Then, the nerve was osmicated by adding 1% osmium tetroxide, 0.1M cacodylate buffer pH=7.3 for 1 and half hour at 4°C. Then, the nerve was washed 2x with dd H_2_O for 15 minutes each. Samples were dehydrated by washing progressively in: 25% ethanol for 5 minutes, 50% ethanol for 5 minutes, 70% ethanol for 5 minutes, 90% ethanol for 10 minutes, 100% ethanol for 10 minutes (x4), propylene oxide for 10 minutes (x3). They were then changed into a 50:50 mixture of Agar 100 resin:propylene oxide for 1 hour at RT. The final change was into a 75:25 mixture of Agar 100 resin:propylene for 2 hours at RT. Nerves were blocked in resin and left shaking O/N at RT. These nerves were re-blocked the following day with fresh resin for 2 hours at RT. The nerves were finally embedded in fresh resin and left in the oven for 24 hours at 65°C. Transverse ultrathin sections from nerves were taken 5 mm from the sciatic notch and mounted on film (no grid bars). Images were taken using a Jeol 1010 electron microscope with a Gatan camera analyzed with ImageJ software.

### SDS-PAGE and immunoblotting

Sciatic nerves were homogenized at 4°C in RIPA buffer (PBS, 1% Nonidet P-40, 0.5% sodium deoxycholate, 0.1% SDS, and 5 mM EGTA) containing protease inhibitors (Mini Protease Inhibitor Cocktail; Sigma-Aldrich) and phosphatase inhibitors (Phosphatase Inhibitor Mini Tablets; Fisher Scientific). We homogenized the tissue using Bullet Blender Homogenizer BBX24-CE (Next Advance) and then sonicated for 4 minutes (30 sec on / off) using a Biorruptor Pico (Diagenode). Protein concentrations were determined by the BCA method (Thermo Scientific). 10 μg of total protein was subjected to SDS-PAGE and blotted on to Protran nitrocellulose membrane (Amersham Biosciences). Membranes were blocked using 5 % milk (Sigma-Aldrich) in TBS 1 % and incubated for 16 hours at 4°C with the indicated primary antibody, washed and incubated with secondary antibodies, and developed with ECL Prime (Amersham). Antibodies used can be found in Source data section online (Key Resources Table). We used an Amersham Imager 680 machine (Amersham) for visualization.

### Chromatin immunoprecipitation assays (ChIP)

Chromatin immunoprecipitation assays (ChIP): The ChIP assay was a modification of the method described by Jang et al (Jang et al., 2006). Schwann cell cultures were incubated in PBS/1% PFA for 10 min at room temperature, harvested by centrifugation (1,000 g, 5min, 4°C) and washed with PBS. The pellet was resuspended in 1 ml of buffer A (50 mM Hepes-KOH, pH 8.1, 1 mM EDTA, 0.5 mM EGTA, 140 mM NaCl, 10 % glycerol, 0.5 % NP40, 0.25 % Triton X-100 and protease inhibitors), homogenized, and sonicated (15 pulses of 30 s separated) in a Biorruptor Pico (diagenode). Chromatin was clarified by centrifugation at 17,000 g for 3 min at room temperature. Protein concentration in the supernatant was quantified by the BCA method (Thermo Scientific). An aliquot was saved as input. The volume corresponding to 60–100 μg of protein was incubated with the corresponding antibody and Dynabeads Protein G (Life Technologies) overnight at 4°C to form immunocomplexes. For in vivo ChIP, freshly dissected uninjured and injured nerves were incubated in PBS/1% PFA for 10 min at room temperature and then quenched for 5 min with glycine 0.125 M. Nerves were washed in PBS for 10 min at 4°C and then lysed in 200 µl of buffer A, using Bullet Blender Homogenizer BBX24-CE (Next Advance). Nuclei were harvested by centrifugation (10,000 g, 5min, 4°C) and washed with 1 ml of buffer B (10 mM Tris-HCl, pH 8.0, 1 mM EDTA, 0.5 mM EGTA, 200 mM NaCl and protease inhibitors), and sonicated (15 pulses of 30 s separated) in a Biorruptor Pico (diagenode). Chromatin was clarified by centrifugation at 17,000 g for 3 min at room temperature. Protein concentration in the supernatant was quantified by the BCA method (Thermo Scientific). AlAn aliquot was saved as input. The volume corresponding to 200–300 μg of protein was incubated with the corresponding antibody and Dynabeads Protein G (Life Technologies) overnight at 4°C to form immunocomplexes. In both cases, immune complexes were centrifuged (500 g, 3 min) and washed twice with 1 ml of “low-salt buffer” (0.1% SDS, 1% Triton X- 100, 2 mM EDTA, 20 mM Tris, pH 8.1, 150 mM NaCl, and protease inhibitors; Roche), and then washed once with 1 ml of “high-salt buffer” (the same but with 500 mM NaCl) and washed three times with 1 ml of LiCl buffer (0.25 M LiCl, 1% IGEPAL, 1% sodium deoxycholate, 1 mM EDTA, 10 mM Tris, pH 8.1, and protease inhibitors). Chromatin from immunocomplexes and input was eluted with 200 μl of 1% SDS, 0.1 M NaHCO3, and 200 mM NaCl and incubated at 65°C for 6 h (to break the DNA–protein complexes). DNA was purified using a column purification kit (GE Healthcare) and submitted to 5× PyroTaq EvaGreen qPCR Mix Plus (CMB) qPCR with the indicated primers.

The ChIP-Seq experiment was performed following a single-end sequencing strategy. High quality reads were aligned against the reference genome (Rattus norvegicus (Rnor_6.0)) with Minimap2 (https://github.com/lh3/minimap2). Read duplicates from the PCR amplification step in the sequencing process were removed with Picard tools (https://broadinstitute.github.io/picard/) and only uniquely mapped reads were kept in the alignments. The uniquely mapped reads were obtained with SAMtools (http://www.htslib.org/) filtering by mapping quality >= 30. MACS2 (https://github.com/macs3-project/MACS, version 2.2.4) was used for peak calling and the results were filtered by -log10 FDR >3. To enable a more informative functional interpretation of experimental data, we identified genes close to or having ChIP-Seq tags on their sequence. ChIPseeker (http://bioconductor.org/packages/release/bioc/html/ChIPseeker.html) was used for this step. Rattus norvegicus (Rnor_6.0) was selected as annotation database (https://bioconductor.org/packages/release/data/annotation/html/TxDb.Rnorvegicus.UCSC.rn6.refGene.html).

### *In vivo* recording of compound action potential from mouse sciatic nerves

Mice were deeply anesthetized by intraperitoneal injection of 40 mg/kg ketamine and 30 mg/kg xylazine. The whole length of the right sciatic nerve was then exposed from its proximal projection into the L4 spinal cord to its distal branches innervating gastrocnemius muscles: tibial, sural and common peroneal. Extracellular recording of compound action potentials (CAPs) was carried out by placing the proximal part of the sciatic nerve on an Ag/AgCl recording electrode with respect to a reference electrode (Ag/AgCl) placed inside the contralateral paw of the animal. For electrical stimulation, another electrode was placed in the distal part of the sciatic nerve just before its trifurcation. To avoid nerve desiccation and the consequent axonal death, the nerve was continuously lubricated with paraffin oil (Panreac). To selectively activate Aβ-, Aδ-, and C-fibers, we recorded CAPs evoked by graded electrical stimulations (5, 8, 10 and 15 V intensity, 0.03 ms pulse duration, Grass Instruments S88, A-M Systems) using both normal and inverse polarity. CAPs were amplified (x1000) and filtered (high pass 0.1 Hz, low pass 10kHz) with an AC amplifier (DAM 50, World Precision Instruments) and digitalized and stored at 25 kHz in a computer using a CED micro-1401interface and Spike2 v.7.01 software (both from Cambridge Electronic Design). In the CAPs, different waveform components corresponding to A- (β and δ), and C-fiber activation were easily distinguished by latency. The amplitude of the different components and the mean nerve conduction velocity (NCV) were measured. The amplitude of each component was measured from the maximum negative to the maximum positive deflection (Sdrulla et al., 2015). For NCV measurement, we divided the distance between the stimulating and recording electrodes by the latency to the CAP component with the biggest amplitude (Vleggeert-Lankamp et al., 2004). The distance between the stimulating and recording electrodes was measured for each experiment using an 8/0 suture thread.

### Statistics

Values are given as means ± standard error (SE). Statistical significance was estimated with the Student’s t test with or without Welch’s correction, one-way ANOVA with Tukey’s multiple comparisons test, mixed ANOVA with Bonferronís multiple comparisons test, Chi-Squared test and the Mann-Whitney U test. A p-value <0.05 was considered statistically significant. For the parametric tests (t test and ANOVA), data distribution was assumed to be normal (Gaussian), but this was not formally tested. Analysis was performed using GraphPad software (version 6.0). Statistics for each experiment are described in more detail in the legends to figures.

## ACKNOWLEDGMENTS

We would like to thank C. Morenilla-Palao for advice in ChIP and other molecular biology experiments. We thank L. Wrabetz and L. Feltri for *P0-Cre* mice and E. Olson for HDAC mice. We also thank P. Morenilla-Ayala for technical assistance. We thank Prof Rhona Mirsky, University College London and Shaline Fazal, University of Cambridge, for insightful comments on the manuscript This work has been funded by grants from the Ministerio de Economía y Competitividad (BFU2016-75864R and PID2019-109762RB-I00), ISABIAL (UGP18-257 and UGP-2019-128) to H. Cabedo and Conselleria Educació Generalitat Valenciana (PROMETEO 2018/114) to J. Gallar and H. Cabedo. Predoctoral fellowships ACIF/2 017/169 from Generalitat Valenciana (to L. Frutos-Rincón) and FPU16/00283from Ministerio de Universidades are also acknowledged. The Instituto de Neurociencias is a ‘‘Center of Excellence Severo Ochoa” (Ministerio de Economía y Competitividad SEV-2013-0317). The authors declare no competing financial interests.

## Author contributions

S. Velasco-Aviles, N. Patel, A. Casillas-Bajo J.A. Gomez Sanchez and L. Rincón-Frutos performed the experiments. S. Velasco-Aviles, N. Patel, L. Rincón-Frutos, E. Velasco, J. Gallar, P. Arthur-Farraj, J.A. Gomez-Sanchez, and H. Cabedo conceived and designed the experiments. H. Cabedo and J.A. Gomez-Sanchez wrote the paper.

---

**Figure S1:**
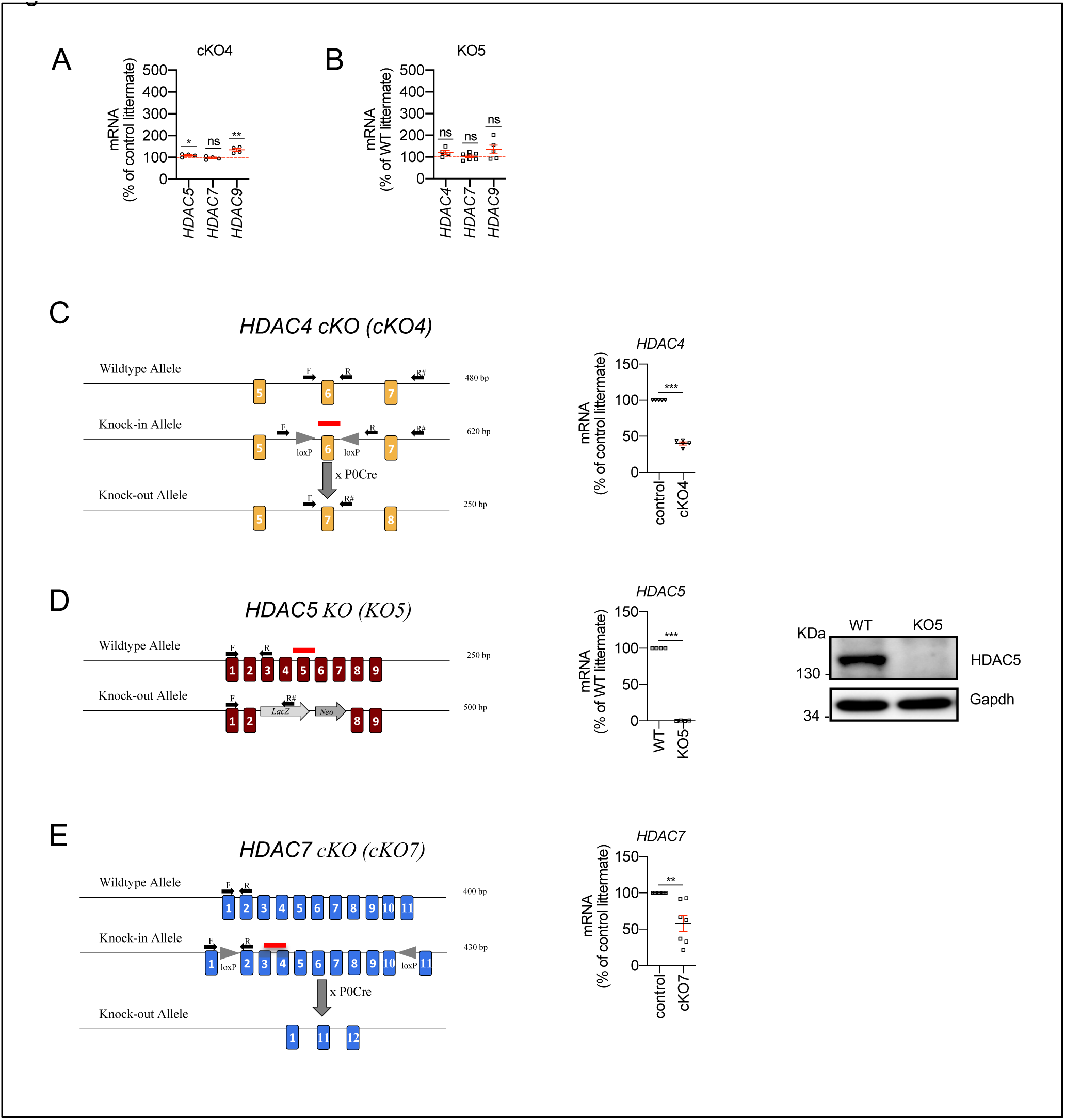
Class IIa HDAC gene expression and removal from Schwann cells. **A)** Only minor or no changes were found for in the expression of HDAC5, HDAC7 or HDAC9 in the nerves of the *cKO4* . **B)** No changes in the expression of HDAC4, HDAC5 and HDAC9 were found in the nerves of the *KO5*. RTqPCR with mouse-specific primers for the indicated genes was performed and normalized to 18S rRNA. The scatter plot, which includes also the mean ± SE, shows the fold change in mRNA normalized to control littermates. 4 to 8 mice per genotype were used. Data were analyzed with the unpaired t-test. **C)** To generate a Schwann cell specific cKO4 mice, the *P0-cre^+/-^* mouse was crossed with a knock-in mouse with the exon 6 of HDAC4 floxed. RTqPCR with specific primers for exon 6 (F and R) demonstrates a decrease by more than 50% in the amount of mRNA in the sciatic nerve. The absence of a complete elimination is probably due to the contribution of mRNA from other cell types. **D)** For HDAC5 we used a complete HDAC5 KO mouse that has no apparent PNS phenotype. In the sciatic nerves of these mice HDAC5 was undetectable both with RT-qPCR and WB. **E)** To generate a Schwann cell specific cKO7 mice, the *P0-cre^+/-^* mouse was crossed with a knock-in mouse with the exons 2 to 10 of HDAC7 floxed. RTqPCR with specific primers (F and R) demonstrates a decrease of about 50% in the amount of mRNA for this HDAC in the sciatic nerve. As in the case of HDAC4, the absence of a complete elimination is probably due to the contribution of mRNA from the axons and other cell types. Primer sequences are described in Source data section online (Key Resources Table). (* P<0,05; ** P<0,01; *** P<0,001; ns: no significant). See Source data section online (Graphs source data) for more details.

**Figure S2:**
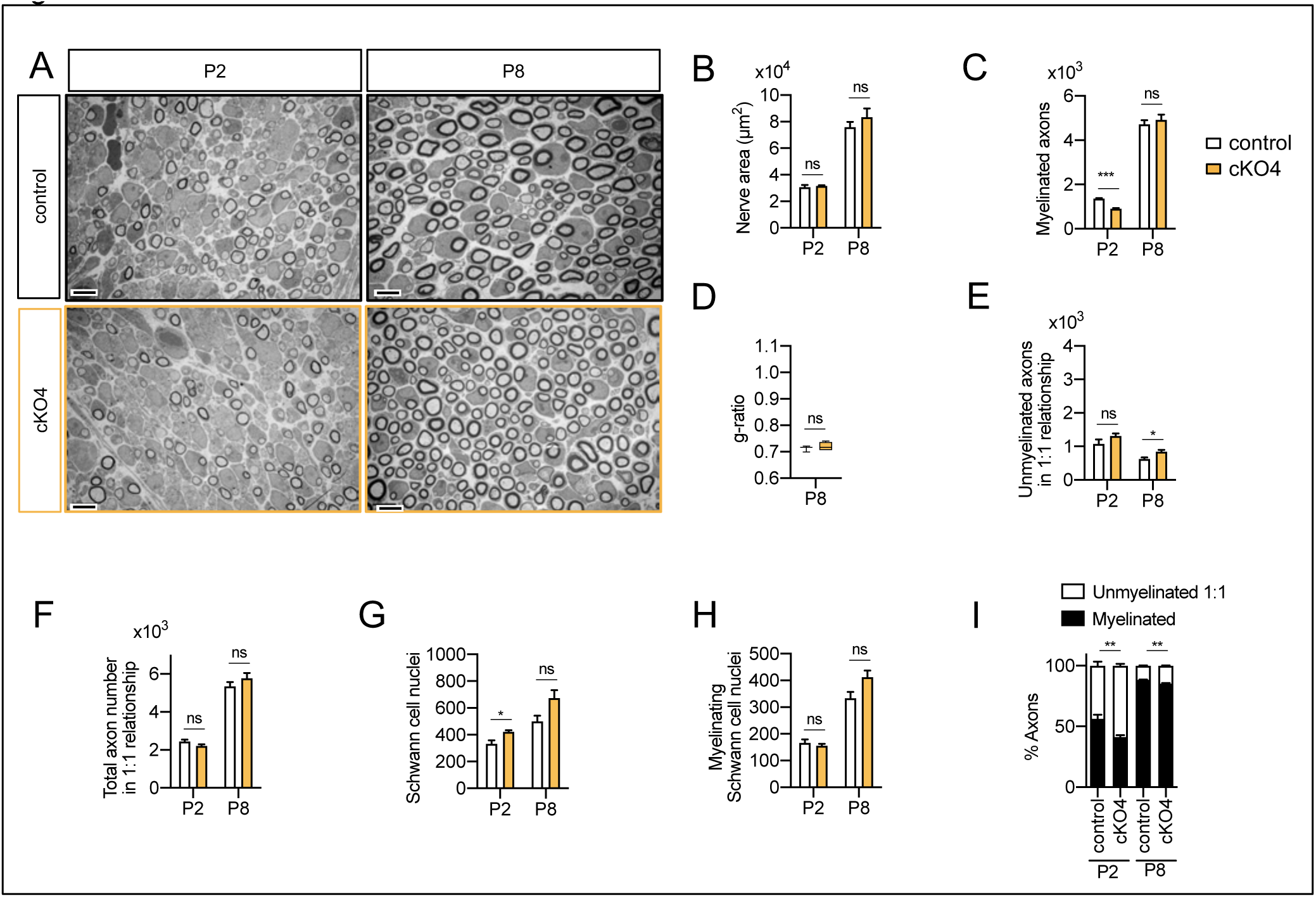
Myelin development in the *cKO4* mice sciatic nerves: Representative transmission TEM images of P2 and P8 sciatic nerves of *cKO4 mice* (*P0-Cre^+/−^; HDAC4^flx/flx^*) and the control (*P0-Cre^−/−^*;*HDAC4^flx/flx^*) littermates. Scale bars 5 μm. **B)** No statistically significant differences were observed between the area of the *cKO4* nerves and control littermates **C)** The number of myelinated axons was slightly decreased at P2 (912 ± 25 in th *cKO4* versus 1363 ± 29 in the control; P<0,0001) but not at P8. **D)** *g-ratio* was not changed at P8. **E)** The number of unmyelinated axons in a 1:1 relationship with Schwann cells was slightly increased at P8 (845 ± 57 in the *cKO4* versus 625 ± 45 in controls; P=0,023). **F)** The total number of sorted axons in a 1:1 relationship with Schwann cells was not changed. **G)** The total number of Schwann cells (counted as nucleus) is slightly increased at P2 (421 ± 12 in the *cKO4* versus 333 ± 25 in controls; P=0,013). **H)** The number of myelinating Schwann cells was not changed. **I)** The percentage of myelinated axons is slightly decreased at P2 (41,17 ± 1,68 % in the *cKO4* versus 56,29 ± 3,39 % in controls; P=0,0037) and P8 (85,37 ± 0,4% in the *cKO4* versus 88,35 ± 0,41 % in controls; P=0,002). For these experiments 4 to 5 animals per genotype were used; Unpaired t-test was used for statistical analysis. (* P<0,05; ** P<0,01; *** P<0,001; ns: no significant). See Source data section online (Graphs source data) for more details. Primer sequences and antibodies are listed in Source data section online (Key Resources Table).

**Figure S3:**
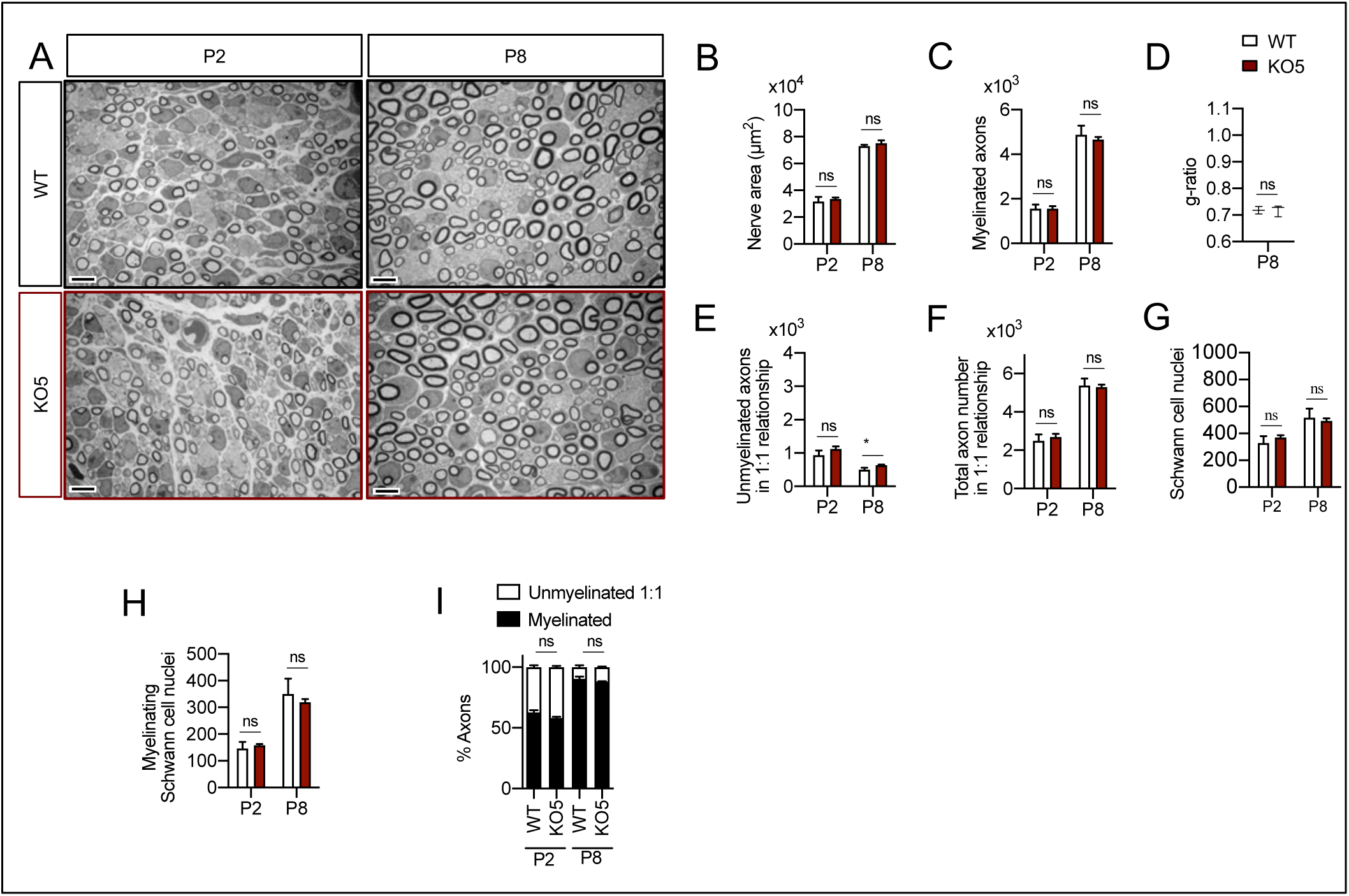
Myelin development in the *KO5* mice sciatic nerves: Representative transmission TEM images of P2 and P8 sciatic nerves of *KO5 mice* (*HDAC5 ^- /-^*) and the control (*HDAC5 ^+/+^*) littermates. Scale bars 5 μm **B)** No statistically significant differences were observed between the area of the *cKO4* nerves and control littermates **C)** The number of myelinated axons was not changed at P2 neither at P8. **D)** *g-ratio* was not changed at P2 neither at P8. **E)** The number of unmyelinated axons in a 1:1 relationship with Schwann cells was slightly increased at P8 (632 ± 19 in the *KO5* versus 503 ± 57 in controls; P=0,044). **F)** The total number of sorted axons in a 1:1 relationship with Schwann cells was not changed. **G)** The total number of Schwann cells (counted as nucleus) was not changed. **H)** The number of myelinating Schwann cells was not changed **I)** The percentage of myelinated axons was not changed. For these experiments 3 animals per genotype were used; Unpaired t-test was used for statistical analysis. (* P<0,05; ** P<0,01; *** P<0,001; ns: no significant). See Source data section online (Graphs source data) for more details.

**Figure S4:**
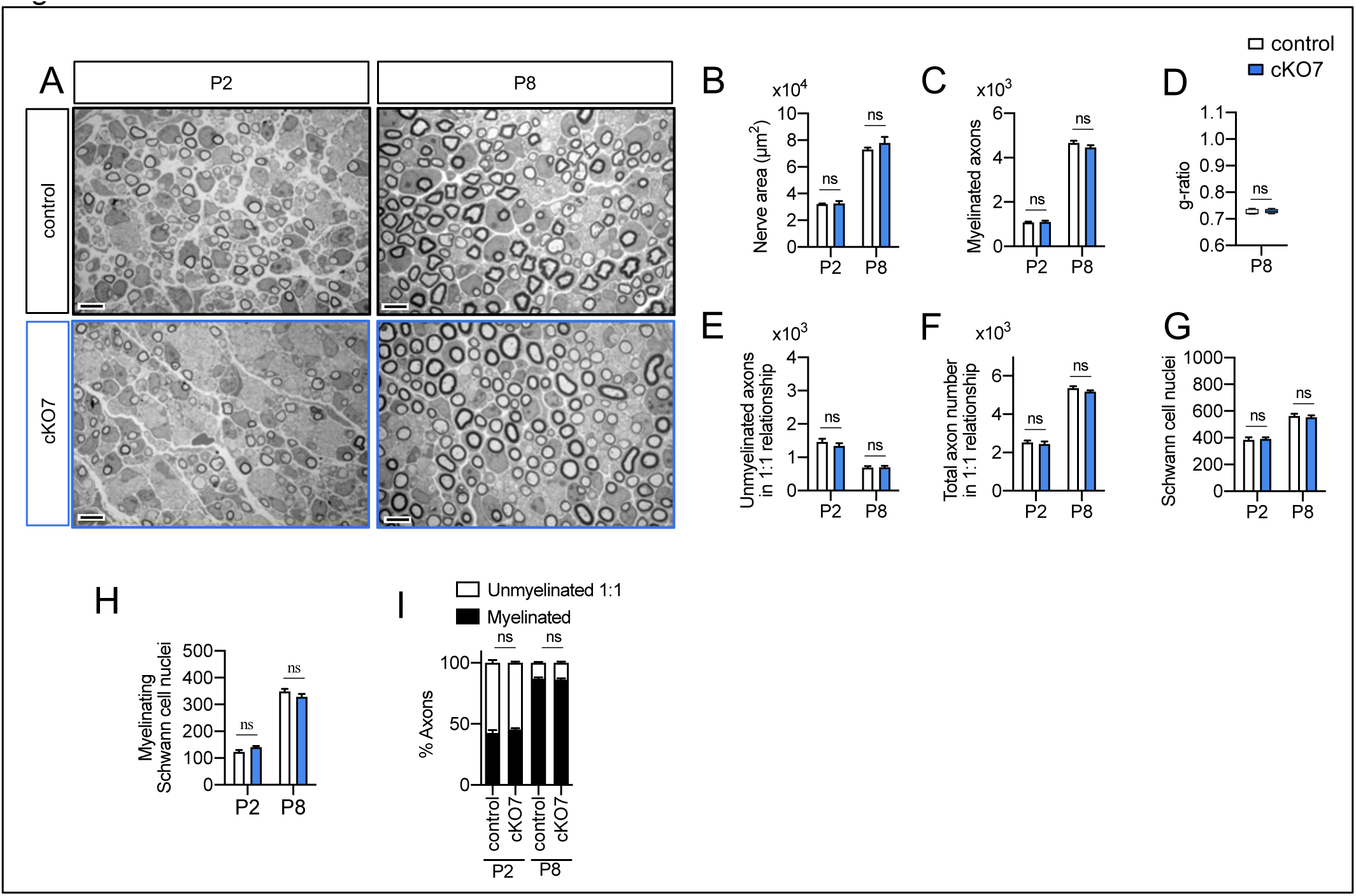
Myelin development in the *cKO7* mice sciatic nerves: Representative transmission TEM images of P2 and P8 sciatic nerves of *cKO7 mice* (*P0-Cre^+/−^;HDAC7 ^flx/flx^)* and the control littermates (*P0-Cre^-/-^;HDAC7^flx/flx^)*. Scale bars 5 μm. No statistically significant differences were observed between the area of the *cKO4* nerves and control littermates **(B)**, number of myelinated axons **(C)**, *g-ratio* **(D)**, number of unmyelinated axons in a 1:1 relationship with Schwann cells **(E),** the total number of sorted axons in a 1:1 relationship with Schwann cells **(F)**, the total number of Schwann cells (counted as nucleus) **(G)**, and number of myelinating Schwann **(H)**. The percentage of myelinated axons was not changed **(I).** For these experiments 3 to 5 animals per genotype were used; Unpaired t-test was used for statistical analysis. (* P<0,05; ** P<0,01; *** P<0,001; ns: no significant). See Source data section online (Graphs source data) for more details.

**Figure S5:**
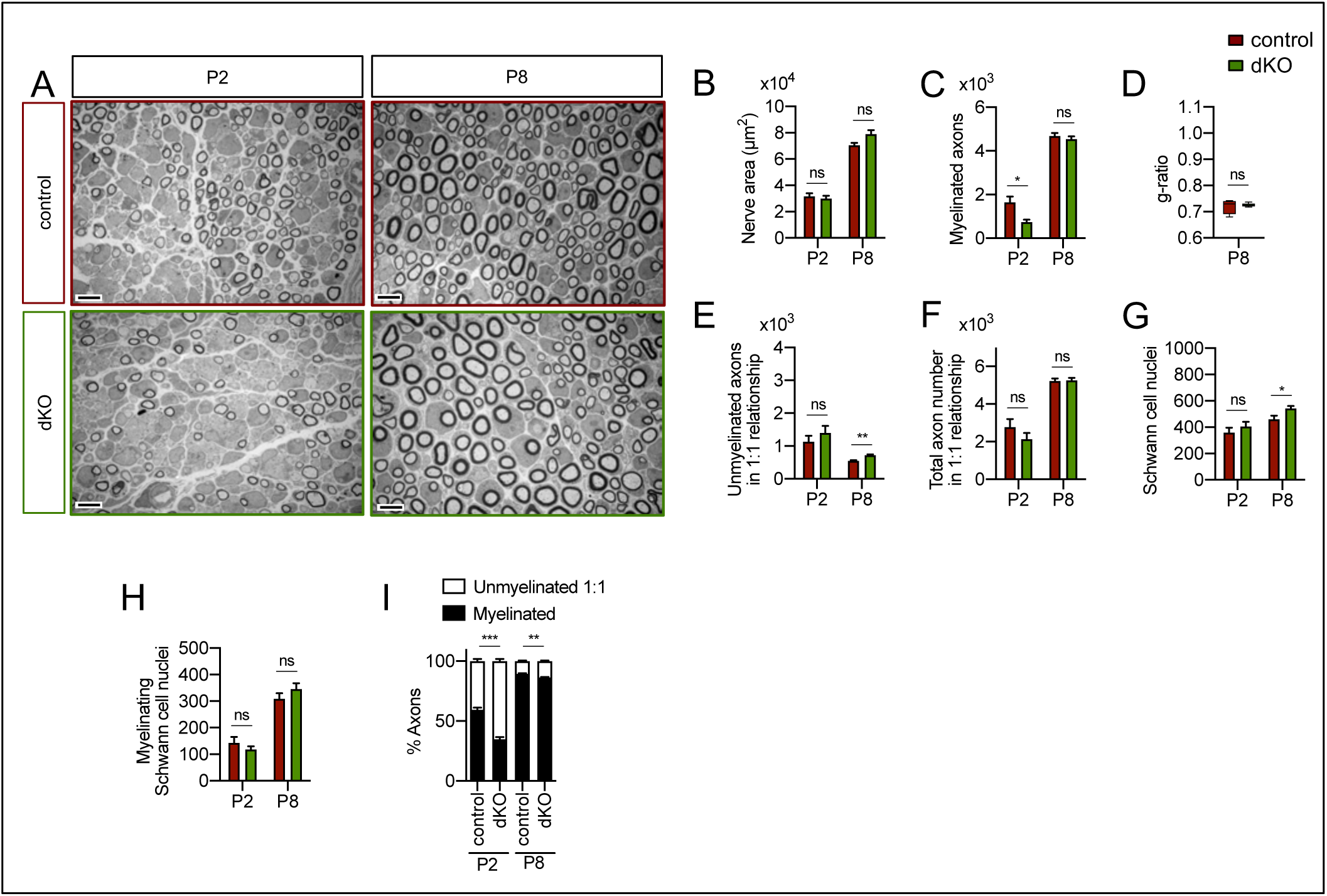
Myelin development in the *dKO* mice sciatic nerves: Representative transmission TEM images of P2 and P8 sciatic nerves of *dKO mice* (*P0-Cre^+/−^;HDAC4^flx/flx^;HDAC5^-/-^)* and the control littermates (*P0-Cre^-/-^;HDAC4^flx/flx^;HDAC5^-/-^)*. Scale bar 5 μm **B)** No statistically significant differences were observed between the area of the *cKO4* nerves and control littermates **C)** The number of myelinated axons is decreased at P2 (743 ± 115 in the *dKO* versus 1645 ± 269 in the control; P=0,0012) but not at P8. **D)** *g-ratio* was not changed at P8. **E)** The number of unmyelinated axons in a 1:1 relationship with Schwann cells was slightly increased at P8 (721 ± 32 in the *dKO* versus 550 ± 15 in controls; P=0,003). **F)** The total number of sorted axons in a 1:1 relationship with Schwann cells was not changed. **G)** The total number of Schwann cells (counted as nucleus) was slightly increased at P8 (543 ± 18 in the *dKO* versus 461 ± 29 in controls; P=0,038). **H)** The number of myelinating Schwann cells was not changed. **I)** The percentage of myelinated axons is decreased at P2 (34,89 ± 1,75% in the *dKO* versus 59,29 ± 1,97 % in controls; P<0,0001) and P8 (86,27 ± 0,62% in the *dKO* versus 89,47 ± 0,49 % in controls; P=0,0061). For these experiments 4 to 5 animals per genotype were used; Unpaired t-test was used for statistical analysis. (* P<0,05; ** P<0,01; *** P<0,001; ns: no significant). See Source data section online (Graphs source data) for more details.

**Figure S6:**
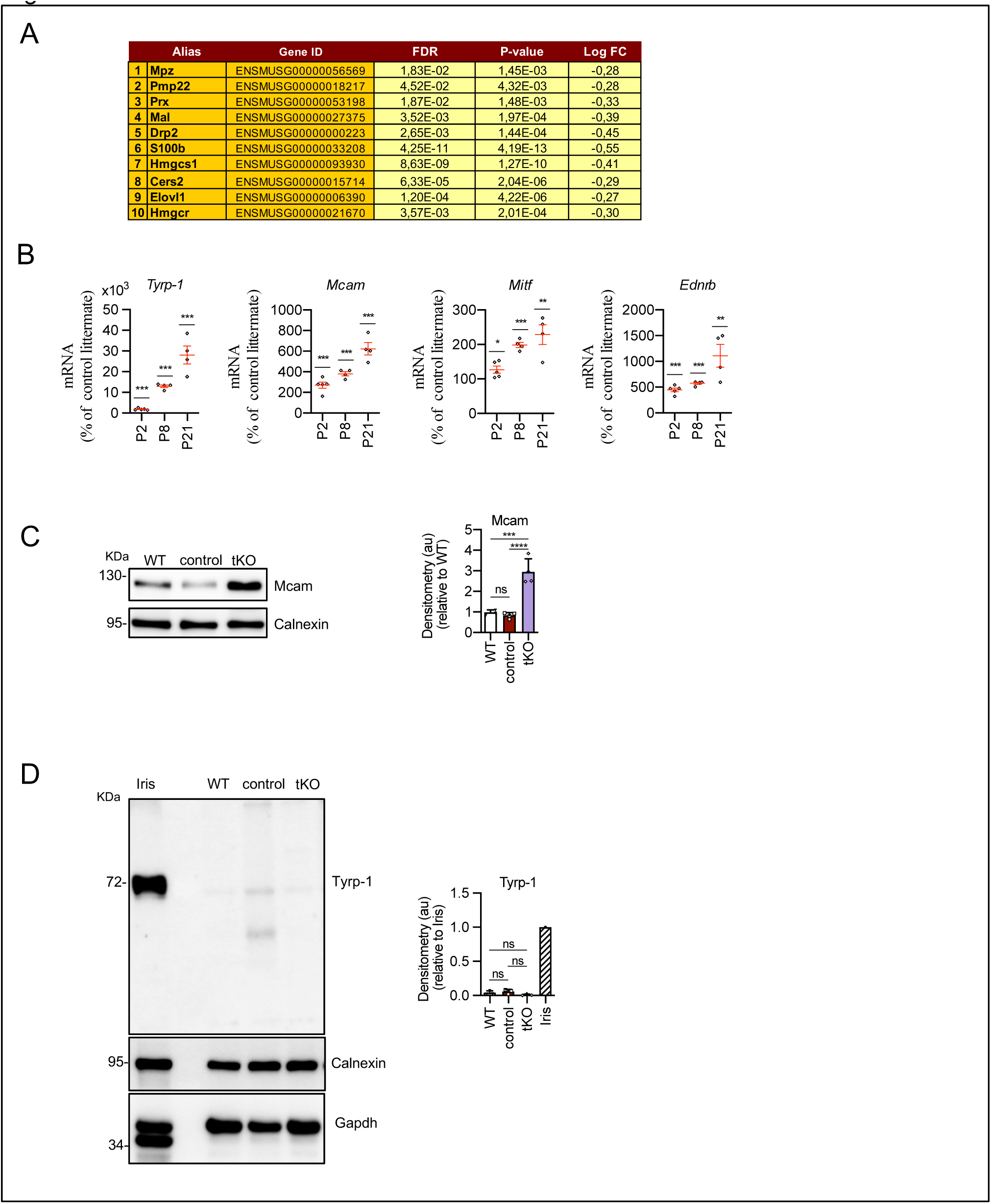
Gene profiling in the *tKO* mice: **A)** RNAseq data shows a slight decrease in the expression of genes encoding for myelin proteins and myelin lipids biosynthesis. FDR, P-value and Log FC is shown. **B)** Melanocyte lineage genes are upregulated during early development. P2, P8 and P21 mouse sciatic nerves were removed and total RNA extracted. RTqPCR with mouse-specific primers for the indicated genes was performed and normalized to 18S rRNA. Graph shows the percentage of mRNA for each gene in the *tKO* normalized to the control littermates. A scatter plot is shown with the results obtained, which include also the mean ± SE. 4/5 mice per genotype and age were used. Data were analyzed with the unpaired t-test. **C)** A representative Western blot of protein extracts obtained from the sciatic nerves of P8 WT, control and *tKO* mice is shown. Calnexin was used as a protein loading control. Densitometric analysis was done for 4 WB and normalized the WT. Data were analyzed with the One-way ANOVA Tukey’s test. **D)** The mRNA for *Tyrp-1* is not translated to protein. A representative Western blot of protein extracts obtained from the sciatic nerves of adult WT, control and *tKO* mice is shown. Iris was used as a positive control. Calnexin and Gapdh were used as protein loading controls. Primer sequences and antibodies are listed in Source data section online (Key Resources Table). (* P<0,05; ** P<0,01; *** P<0,001; ns: no significant). See Source data section online (Graphs source data) for more details.

**Figure S7:**
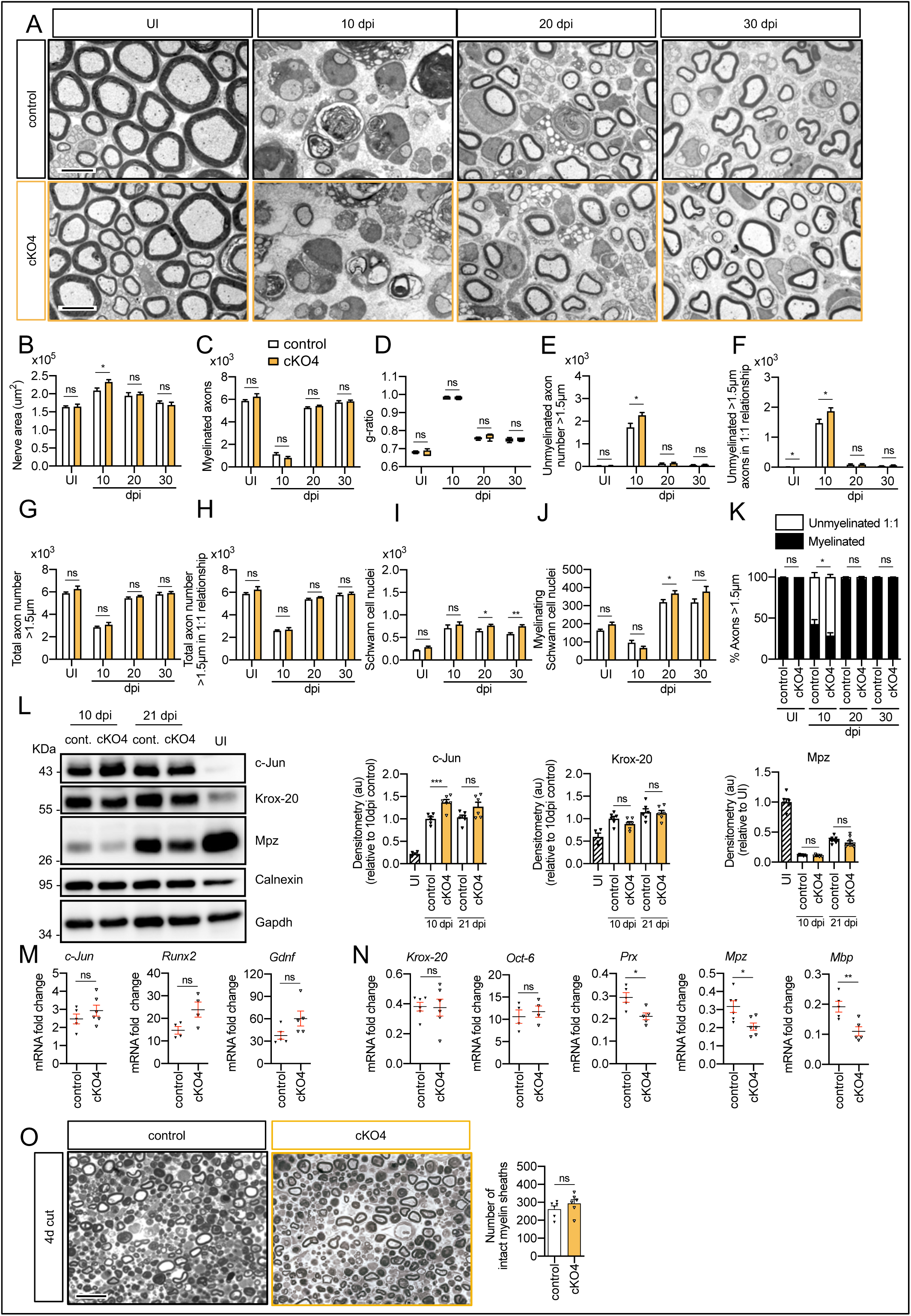
Remyelination in the *cKO4* mice: **A)** Representative transmission TEM images of P60 sciatic nerves uninjured (UI) and 10, 20 and 30 days post crush (dpi) of *dKO* (*P0-Cre^+/−^; HDAC4^flx/flx^*) and the control (*P0-Cre^−/−^*;*HDAC4^flx/flx^*) littermates are shown. Scale bars 5 μm. **B)** Only slight significant statistically differences were observed in the area of the *cKO4* at 10 dpi (P=0.0372). **C)** The number of myelinated axons was not changed at any point. **D)** the same for *g-ratio*. **E)** The number of unmyelinated axons >1,5 μm was increased at 10 dpi in the *cKO4* (1.730 ± 175 in the control versus 2.266 ± 116 in the *cKO4*; P=0,024). **F)** The total number of unmyelinated axons in a 1:1 relationship with Schwann cells is increased at 10 dpi (1.869 ± 115 in the *cK*O*4* versus 1.472 ± 130 in the control; P=0047). No changes in the total axon number was found (**G)** neither in the sorted total axon number **(H)**. **I)** The total number of Schwann cells (counted as nuclei) was increased at 20 dpi (765 ± 29 in the *cKO4* versus 643 ± 41 in controls; P=0,0041) and at 30 dpi (752 ± 34 in the *cKO4* versus 575 ± 32 in controls; P=0,0066). **J)** The number of myelinating Schwann cells was slightly increased at 20 dpi (368 ± 15 in the *cKO4* versus 319 ± 14 in controls; P=0,0045). **K)** The percentage of myelinated axons was decreased at 10 dpi (29,0 ± 3,2 % in the *cKO4* versus 42,8 ± 5,4 % in controls; P=0,0474). For these experiments 4 to 7 animals per genotype were used; Unpaired t-test was applied for statistical analysis. **L)** A representative WB of protein extracts from *cKO4* and control nerves is shown. In the quantification, c-Jun protein was higher in the *cKO4* at 10 dpi and tended to equalize at 21 dpi. No changes were found in Krox20 or Mpz. Densitometric analysis was done on 4 to 6 WB from the same number of mice. Data were analyzed with the unpaired t-test. **M)** Expression of *c-Jun*, *Runx2* and *Gdnf* at 10 dpi was not changed in the sciatic nerves of the *cKO4*. **N)** Expression of *Krox20* and *Oct6* was not changed, whereas *Prx*, *Mpz* and *Mbp* expression was decreased in the *cKO4* sciatic nerve at 10 dpi. RTqPCR with mouse-specific primers for the indicated genes was performed and normalized to 18S rRNA. The scatter plot, which include also the mean ± SE, shows the fold change of mRNA for each gene at 10 dpi normalized to the uninjured nerve. 4 to 6 mice per genotype were used. Data were analyzed with the unpaired t-test. **O)** A representative toluidine blue staining image of 4 days-cut sciatic nerve of *cKO4* and control mice is shown. The quantification of intact myelin sheaths showed no changes in the *cKO4*. 6 animals were used for the experiment. Data were analyzed with the unpaired t-test. Primer sequences and antibodies are listed in Source data section online (Key Resources Table). (* P<0,05; ** P<0,01; *** P<0,001; ns: no significant). See Source data section online (Graphs source data) for more details.

**Figure S8:**
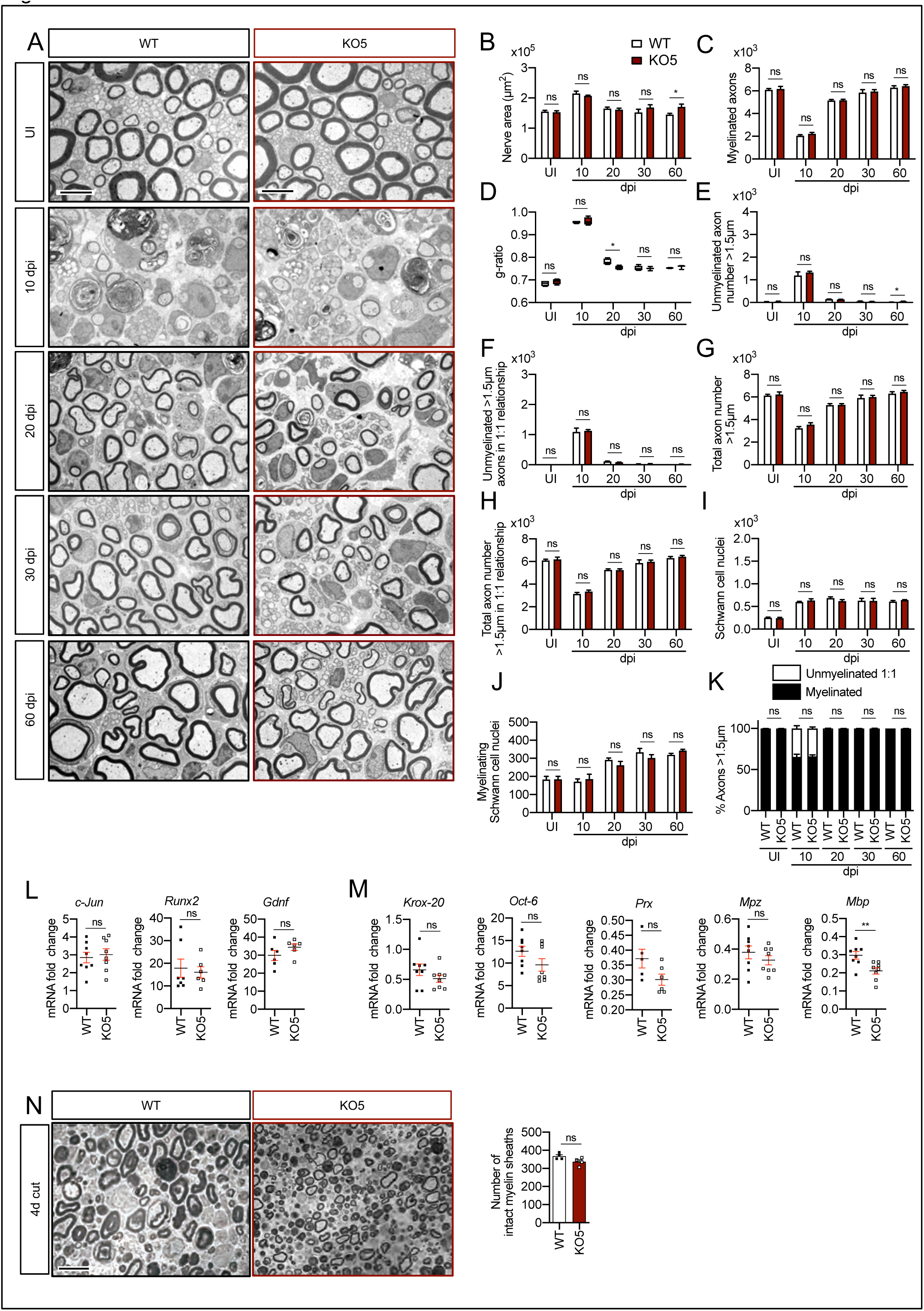
Remyelination in the *KO5* mice: **A)** Representative transmission TEM images of P60 sciatic nerves uninjured (UI) and 10, 20,30 and 60 days post crush (dpi) of *KO5 (HDAC5 ^-/-^*) and the WT control littermates are shown. Scale bars 5 μm. **B)** A slight increase in the area of the *KO5* at 60 dpi (P=0.0419) was found. **C)** The number of myelinated axons was not changed at any point. **D)** The *g-ratio* was found slightly decreased in at 20 dpi in the *KO5* (0,75 ± 0,004 in the *KO5* versus 0,78 ± 0,007 in the control; P=0,0154). **E)** The number of unmyelinated axons >1,5 μm was slightly increased at 60 dpi in the *KO5* (52 ± 10 in the control versus 22 ± 6 in the *KO5*; P=0,0352). The total number of unmyelinated axons in a 1:1 relationship with Schwann cells **(F)**, the total axon number (**G),** the total number of sorted axons **(H**), the total number of Schwann cells (counted as nuclei) **(I)**, the number of myelinating Schwann cells **(J)** neither the percentage of myelinated axons **(K)** were found changed at any point. For these experiments 3 to 5 animals per genotype were used. Unpaired t-test was applied for statistical analysis. **L)** Expression of *c-Jun*, *Runx2* and *Gdnf* at 10 dpi was not changed in the sciatic nerves of the *KO5*. **M)** Expression of *Krox20*, *Oct6, Prx and Mpz* was not changed, whereas *Mbp* expression was slightly decreased in the *KO5* sciatic nerve at 10 dpi. RTqPCR with mouse-specific primers for the indicated genes was performed and normalized to 18S rRNA. The scatter plot, which include also the mean ± SE, shows the fold change of mRNA for each gene at 10 dpi normalized to the uninjured nerve. 4 to 6 mice per genotype were used. Data were analyzed with the unpaired t-test. **O)** A representative toluidine blue staining image of 4 days-cut sciatic nerve of *KO5* and control mice is shown. The quantification of intact myelin sheaths showed no changes in the *KO5*. 6 animals were used for the experiment. Data were analyzed with the unpaired t-test. Primer sequences are listed in Source data section online (Key Resources Table). (* P<0,05; ** P<0,01; *** P<0,001; ns: no significant). See Source data section online (Graphs source data) for more details.

**Figure S9:**
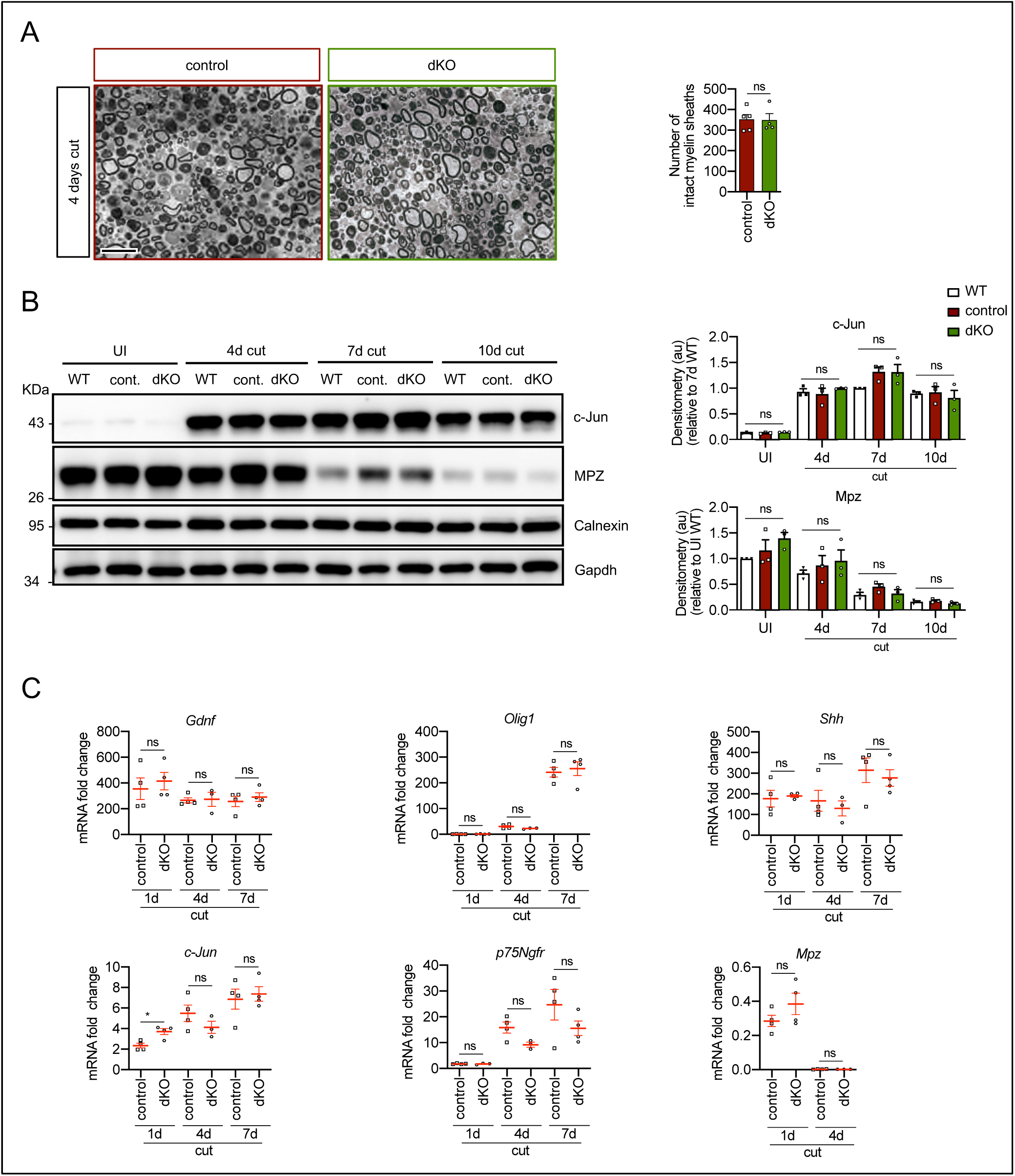
Myelin clearance and repair phenotype activation in the *dKO* mice. **A)** A representative toluidine blue staining image of 4 days-cut sciatic nerve of *dKO* and control mice is shown. The quantification of intact myelin sheaths showed no changes. 4 to 5 animals were used for the experiment. Data were analyzed with the unpaired t-test. **B)** WB against c-Jun and Mpz supports that myelin clearance is normal in the *dKO* nerves. Calnexin and Gapdh were used as housekeeping. Three mice per genotype were analyzed independently by densitometry. Data were analyzed with the unpaired t-test. **C)** Repair phenotype activation was determined by measuring the expression of marker genes and comparing with the uninjured control nerve. As is shown only a slight increase in the expression of c-Jun at 1 day after cut in the *dKO* was found (3,71 ± 0,24 in the *dKO* versus 2,35 ± 0,22 in the control; P=0,0102). RTqPCR with mouse-specific primers for the indicated genes was performed and normalized to 18S rRNA. Graph shows the percentage of mRNA for each gene in the injured nerve normalized to the uninjured controls. A scatter plot is shown with the results obtained, which include also the mean ± SE. 3 to 4 mice per genotype were used. Data were analyzed with the unpaired t-test. Primer sequences and antibodies are listed in Source data section online (Key Resources Table). (* P<0,05 ** P<0’01 *** P<0’001; ns: not significant). See Source data section online (Graphs source data) for more details.

**Figure S10:**
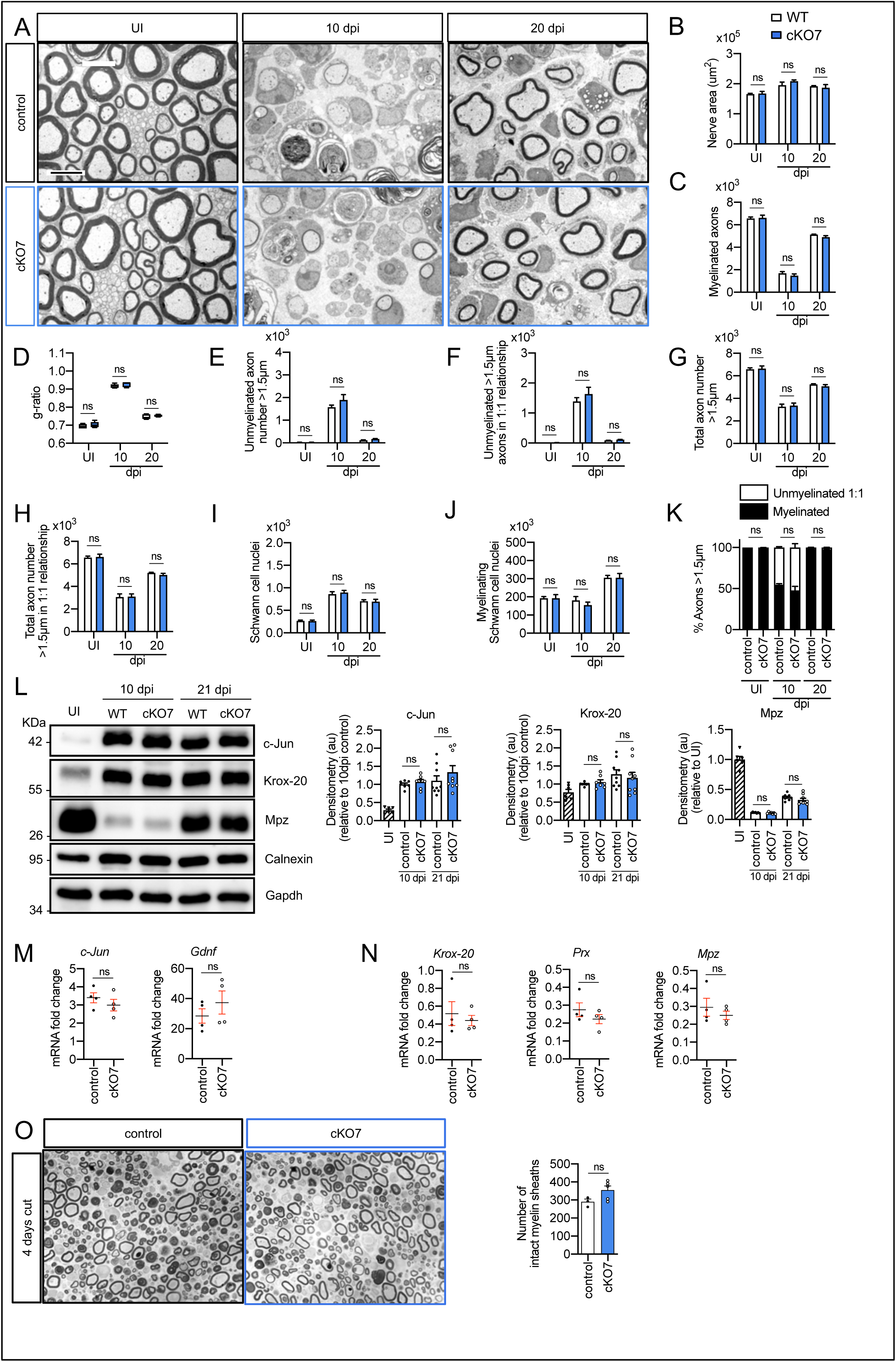
Remyelination in the *cKO7* mice: **A)** Representative transmission TEM images of P60 sciatic nerves uninjured (UI) and 10 and 20 days post crush (dpi) of c*KO7 (P0-Cre^+/−^;HDAC7 ^flx/flx^*) and the WT control littermates *(P0-Cre^-/−^;HDAC7 ^flx/flx^*) are shown. Scale bars 5 μm. No changes were found at any point in the nerve area (**B**), the number of myelinated axons(**C**), *g-ratios* (**D**), number of unmyelinated axons >1,5 μm (**E**),the total number of unmyelinated axons >1,5 μm in a 1:1 relationship with Schwann cells **(F)**, the total axon number (>1,5 μm) (**G**), the total number of sorted axons **(H**), the total number of Schwann cells (counted as nuclei) **(I)**, the number of myelinating Schwann cells **(J)** neither the percentage of myelinated axons **(K)**. For these experiments 3 to 5 animals per genotype were used. Unpaired t-test was applied for statistical analysis. **L**) A representative Western blot of protein extracts obtained from sciatic nerves UI, 10 and 21 days post crush (pdi) is shown. Densitometric quantification shows no differences between phenotypes.6 to 9 mice were used for quantification. **M**) No changes were found in the mRNA for *c-Jun* and *Gdnf* at 10 dpi. **N)** No changes were found for *Krox20*, *Prx* and *Mpz* at 10 dpi. RTqPCR with mouse-specific primers for the indicated genes was performed and normalized to 18S rRNA. Graph shows the percentage of mRNA for each gene in the crush injured (10 dpi) nerve normalized to the uninjured controls. A scatter plot is shown with the results obtained, which include also the mean ± SE. 4 mice per genotype were used. Data were analyzed with the unpaired t-test. **O**) A representative toluidine blue staining image of 4 days-cut sciatic nerve of *cKO7* and control mice is shown. The quantification of intact myelin sheaths shows no changes in the *cKO7*. 3 to 5 animals were used for the experiment. Data were analyzed with the unpaired t-test. Primer sequences and antibodies are listed in Source data section online (Key Resources Table). (* P<0,05 ** P<0’01 *** P<0’001; ns: not significant). See Source data section online (Graphs source data) for more details.

**Figure S11:**
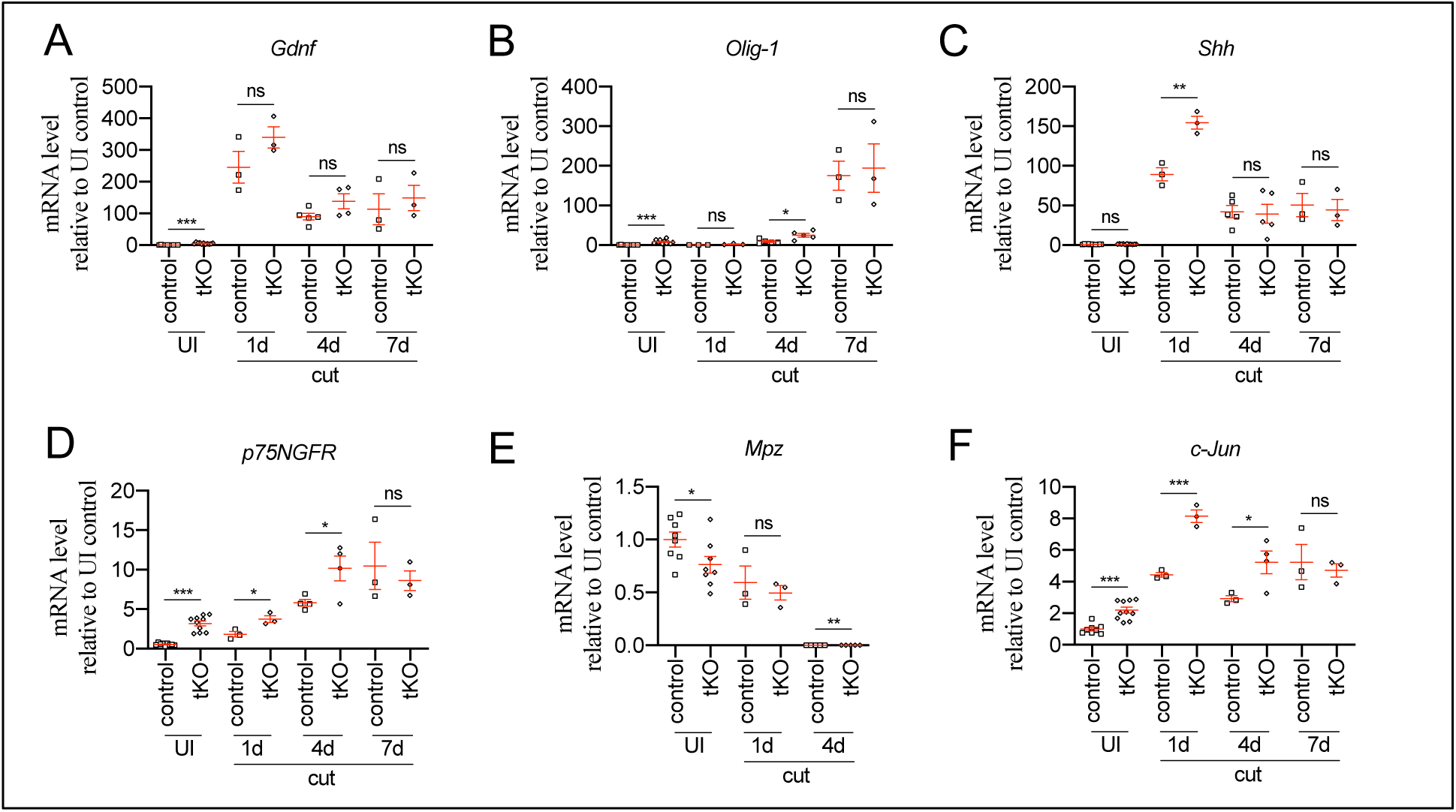
Repair Schwann cell gene expression in the *tKO*. *tKO* Schwann cells express efficiently markers of the repair phenotype after a cut experiment. In some time-points, c-Jun (**A**), *Olig1* (**B**), *Shh* (**C**) and *p75Ngfr* (**D**) were even more expressed in the *tKO* than in the control nerves. Mpz was decreased in the UI and at 4 days post cut (**E**). No changes were found in *Gdnf* (**F**). A cut experiment was performed (P60) and sciatic nerves removed at 1, 4 and 7 days postinjury. RTqPCR with mouse-specific primers for the indicated genes was performed and normalized to 18S rRNA. Graph shows the percentage of mRNA for each gene in the crush injured (10 dpi) nerve normalized to the uninjured controls. A scatter plot is shown with the results obtained, which include also the mean ± SE. 3 to 5 animals per time point and genotype were used, except for uninjured (10-11 nerves). Data were analyzed with the unpaired t-test. Primer sequences are listed in Source data section online (Key Resources Table). (* P<0,05 ** P<0’01 *** P<0’001; ns: not significant). See Source data section online (Graphs source data) for more details.

**Figure S12:**
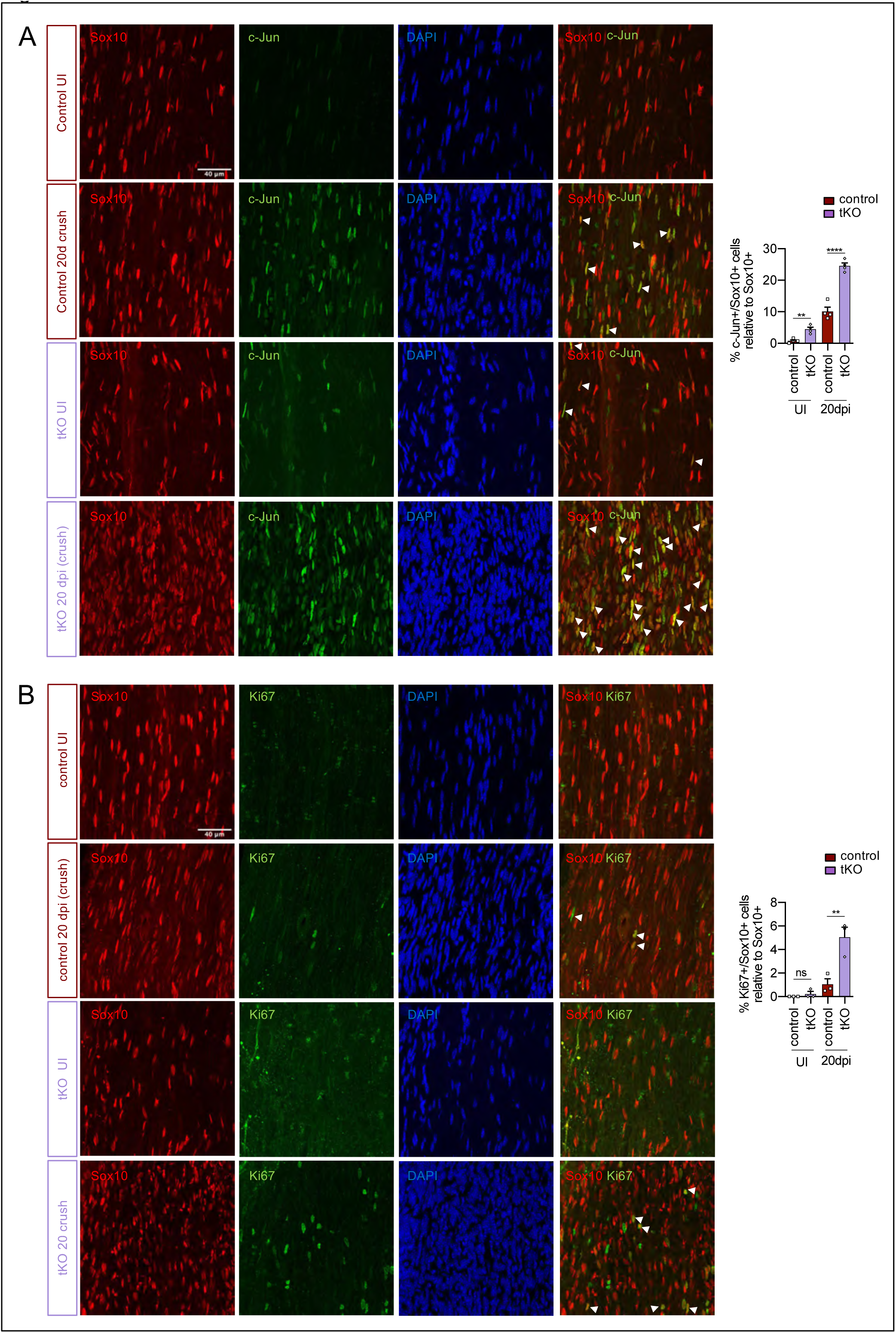
Increased c-Jun and Schwann cell proliferation in the *tKO* sciatic nerve. **A)** *tKO* mice sciatic nerves showed increased numbers of Schwann cells (Sox10^+^) expressing c-Jun both before and 20 days post crush. **B)** *tKO* mice sciatic nerves show increased numbers of Schwann cell (Sox10^+^) proliferating (Ki67^+^) at 20 dpi. Uninjured and 20 dpi crushed sciatic nerves (P60) were fixed and submitted to immunoflorescence with the indicated antibodies. Nuclei were counterstained with Hoechst. Representative confocal images of sections obtained from the sciatic nerves of wild type (WT), control and *tKO* mice are shown. Scale bars= 20 μm. For the quantification 4 animals per genotype were used. Data were analyzed with the unpaired t-test. Antibodies used are listed in Source data section online (Key Resources Table).(* P<0,05 ** P<0’01 *** P<0’001; ns: not significant). See Source data section online (Graphs source data) for more details.

**Figure S13:**
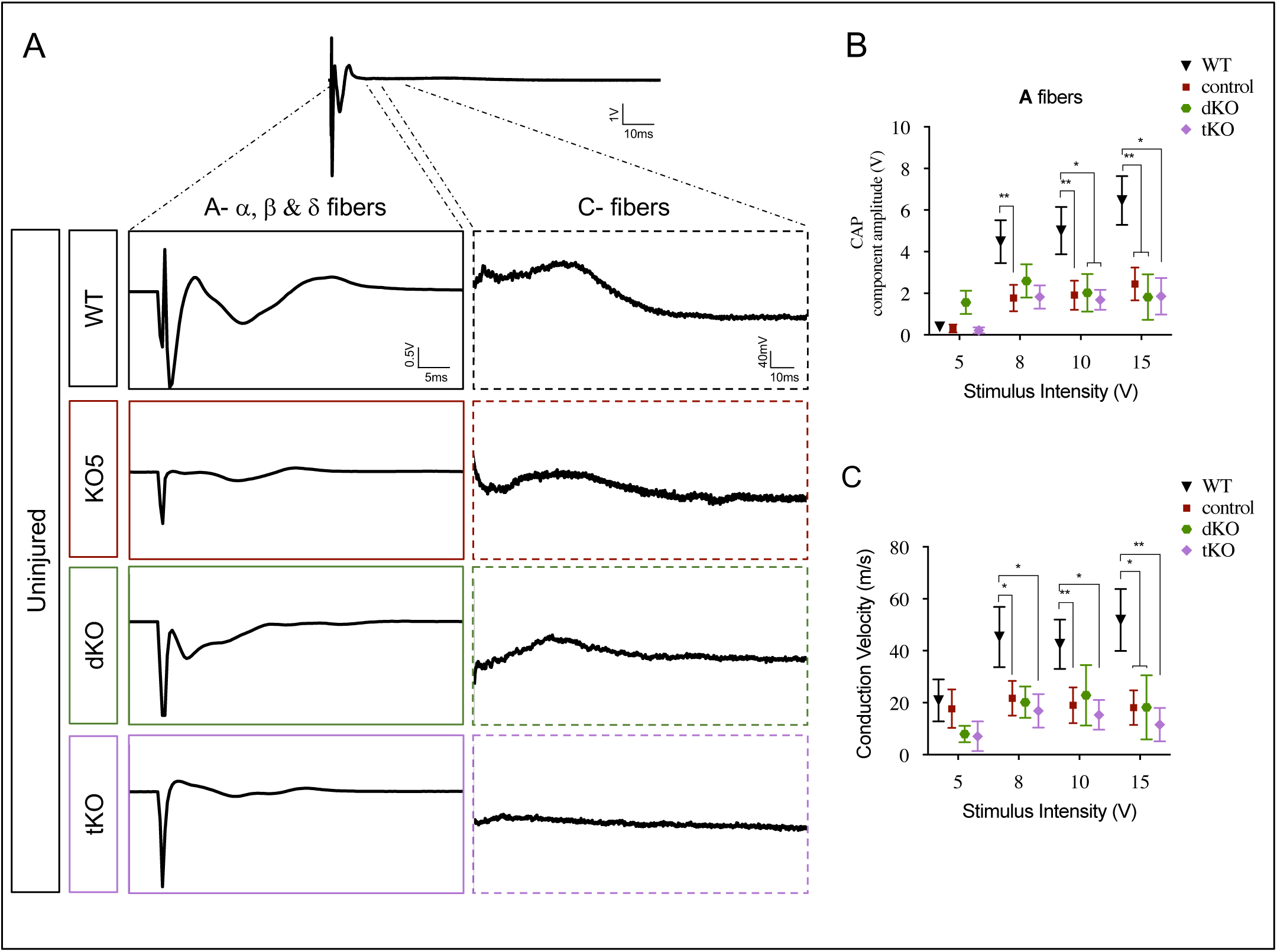
HDAC5 elimination decreases nerve conduction velocity and voltage amplitude. **A)** Sample recordings of compound action potential (CAP) in uninjured sciatic nerves of WT, *KO*5,*dKO* and *tKO* mice showing the waveform components corresponding to myelinated (A fibers) and unmyelinated (C) fibers. **B)** Waveform component of A fibers showed a decreased amplitude for all genotypes (compared to WTs) at pulses of 8, 10 and 15 V. **C)** The same for nerve conduction velocity. In this set of experiments, the whole length of a sciatic nerve was exposed from its proximal projection (L4 spinal cord) to its distal branches in deeply anesthetized mice. Compound action potentials (CAPs) were evoked by electrical stimulation of increasing intensity (5, 8, 10 and 15 V, 0.03 ms pulse duration). The maximum amplitude of the A-fiber components of CAP electrical signal, and their mean nerve conduction velocity were measured. Seven to 18 animals per genotype were used. Mann-Whitney’s U were used for non-parametric paired comparisons.(* P<0,05; ** P<0,01; *** P<0,001; ns: no significant). See Source data section online (Graphs source data) for more details.

**Figure S14:**
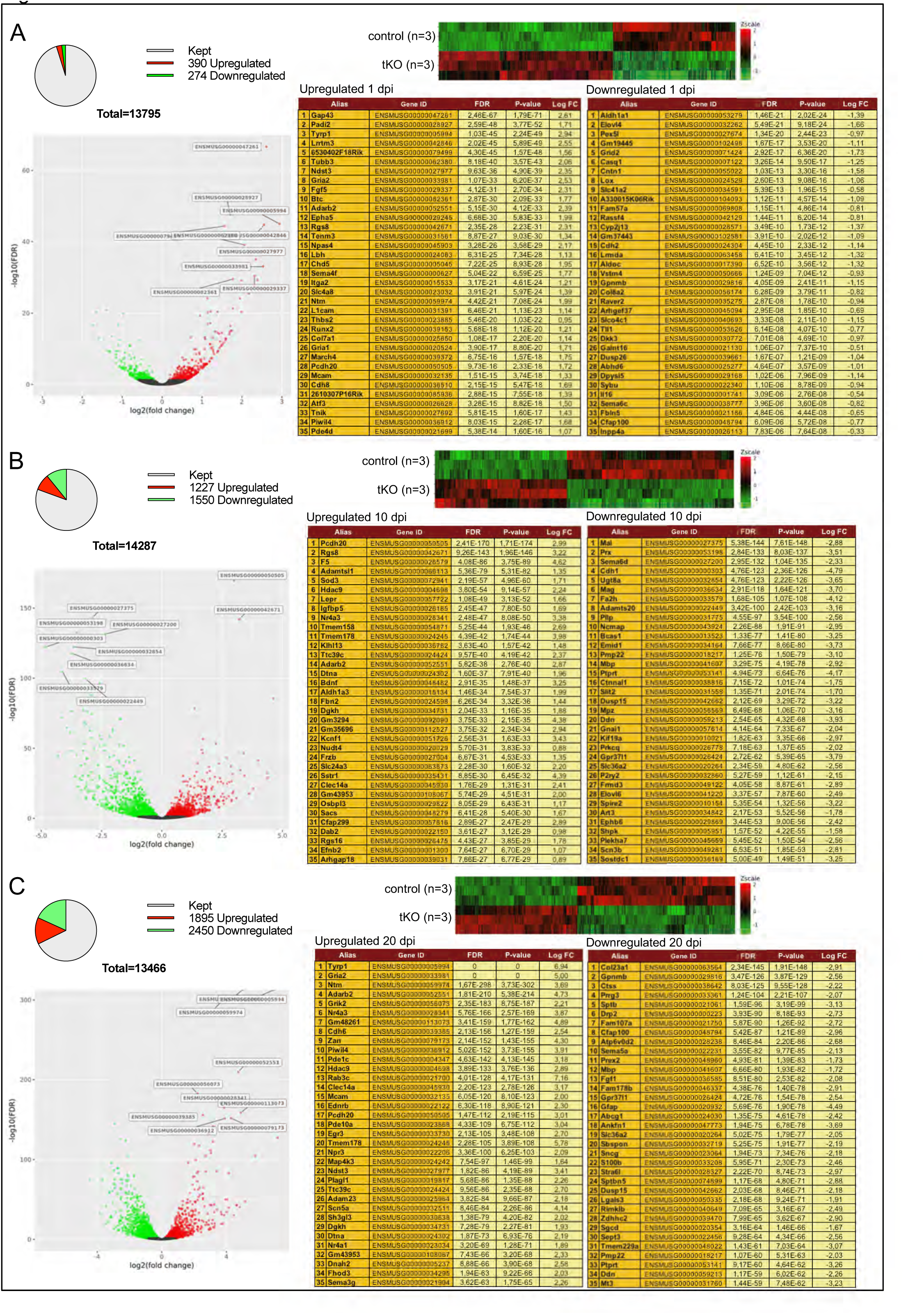
Evolution of sciatic nerve gene expression profile at 1, 10 and 20 dpi. The Pie chart, DEG heatmap, volcano plot and list of the 35 most upregulated and downregulated genes in the *tKO* classified by FDR at 1 dpi (**A**), 10 dpi (**B**) and 20 dpi (**C**) is shown. Data obtained from the RNAseq analysis of 3 animals per condition. See Source data section online (RNAseq source data) for more details.

**Figure S15.**
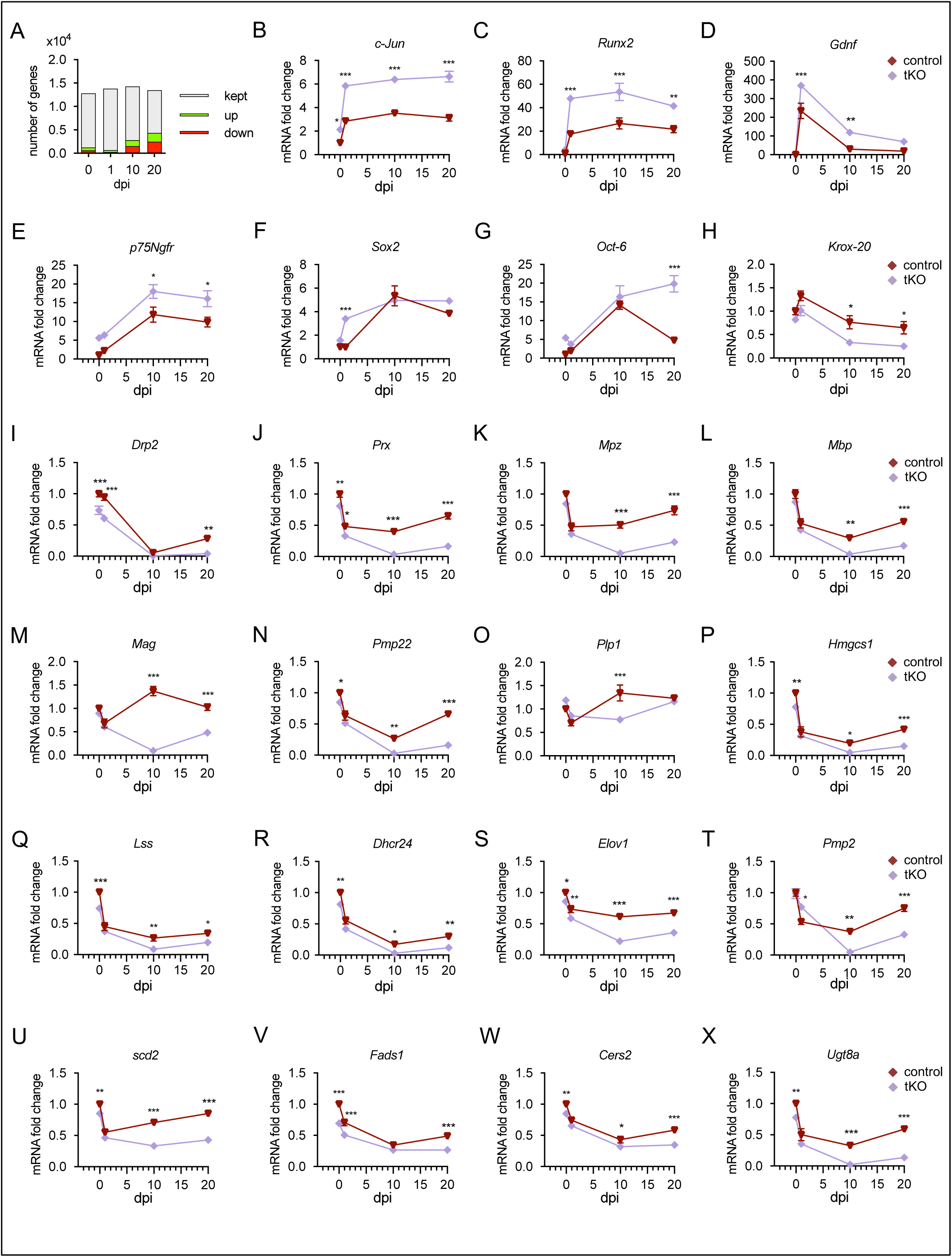
Time course analysis of myelination-relevant gene expression during nerve regeneration from RNAseq. **A)** Bar chart showing an increase in the number of DE genes at different time points after nerve crush. In the injured *tKO* nerve 1.270 genes are DE being 390 upregulated and 274 downregulated. At 1 dpi, the number of genes DE genes decreased to 664, being 390 upregulated and 274 downregulated. At 10 dpi, there were 2.777 genes DE, being 1.227 upregulated and 1.550 downregulated. The bigger differences were found at 20 dpi, with up to 4.345 DE genes, being 1.895 upregulated and 2.450 downregulated. **B-F)** The expression of markers of non-myelin forming Schwann cells, such as *c-Jun*, *Runx2, Gdnf*, *p75Ngfr* and *Sox2*, is enhanced in the sciatic nerves of the *tKO* mice **G)** *Oct-6* is induced in the control and *tKO* animals by more than 10-fold at 10 dpi. Then, at 20 dpi it drops in the control nerves but still grows up in the *tKO* (up to 15-fold). **H-O)** *Krox20* and myelin protein genes are downregulated in the *tKO* nerves **P-R)** Different genes of the sterol branch of the mevalonate pathway (*Hmgcs*, *Lss*, *Dhcr24*) are downregulated in the *tKO*. We also found downregulated genes encoding for enzymes involved in the elongation, (*Elov1*) (**S**), transport (*Pmp2*) (**T**) and the insertion of double bonds (*Scd2, and Fads1*) (**U** and **V**) into fatty acids. Finally, we found downregulated *Cers2* and *Ugt8a*, genes involved in the synthesis of sphingomyelin and galactosyl-ceramide respectively (**W** and **X**). Level of mRNA for each gene (as FPKM) and genotype was normalized to the level of the uninjured control nerve and expressed as fold change. The sciatic nerve of 3 mice per genotype and timepoint were used to perform the RNAseq (see Material and methods). Two-way ANOVA test was used for the analysis. (* P<0,05; ** P<0,01; *** P<0,001; ns: no significant). See Source data section online (Graphs source data) for more details.

**Figure S16:**
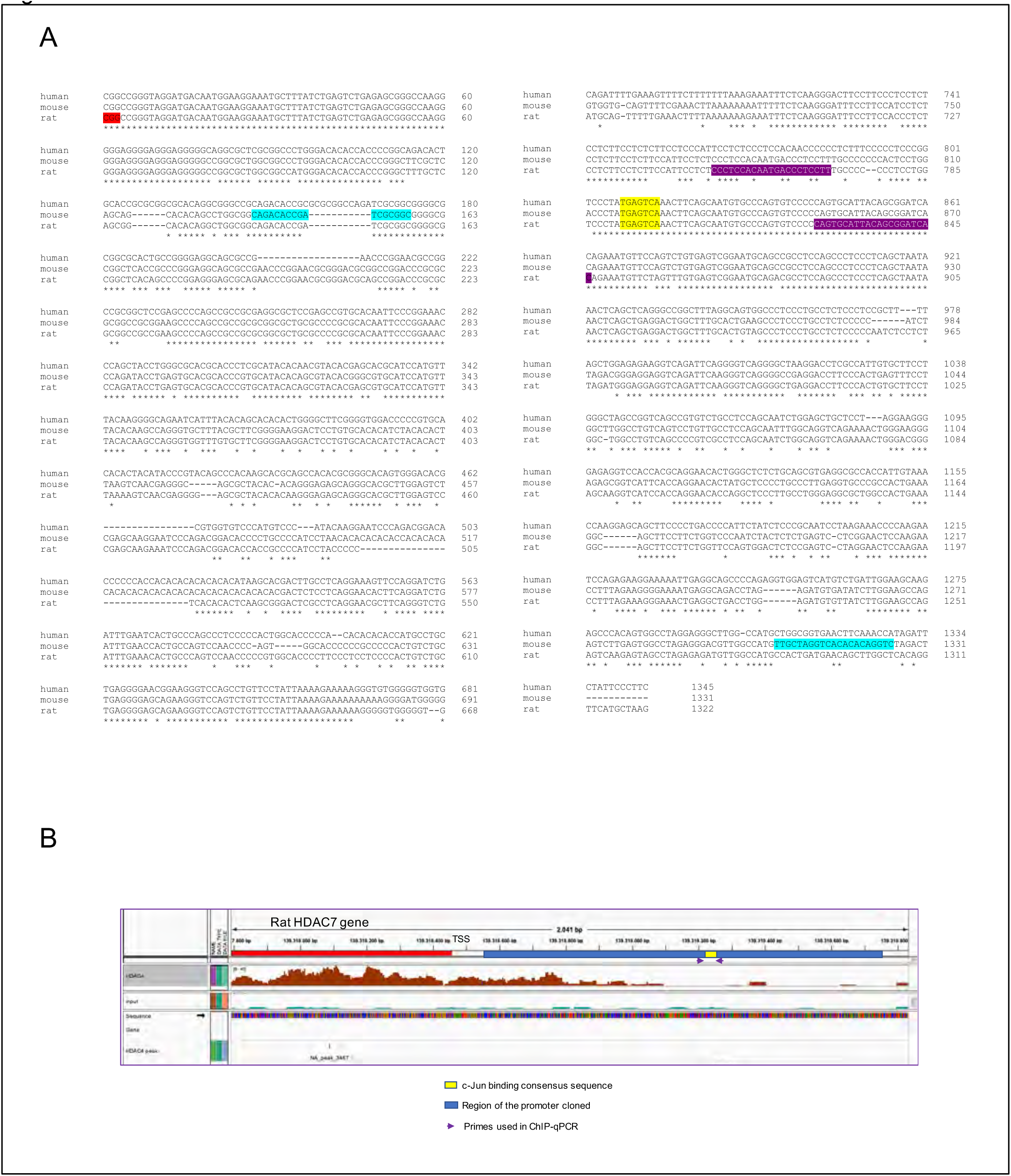
The proximal promoter region of the HDAC7 has a c-Jun consensus binding sequence. **A)** The alignment of the proximal promoter region for human, mouse and rat is shown. A c-Jun consensus binding sequence in the promoter of the mouse HDAC7 gene was identified (yellow). The consensus sequence is conserved in rat and humans. The transcription starting site defined for rat is shown in a red box. In blue, the sequence of the primers used to amplify and clone the mouse HDAC7 promoter region are shown. In purple are the sequences of the primers used for the ChIPqPCR with anti-c-Jun and anti-HDAC4. **B)** Density of reads in the ChIP-seq and mapping of the c-Jun consensus sequence and primer position in the HDAC7 gene.

**Figure S17:**
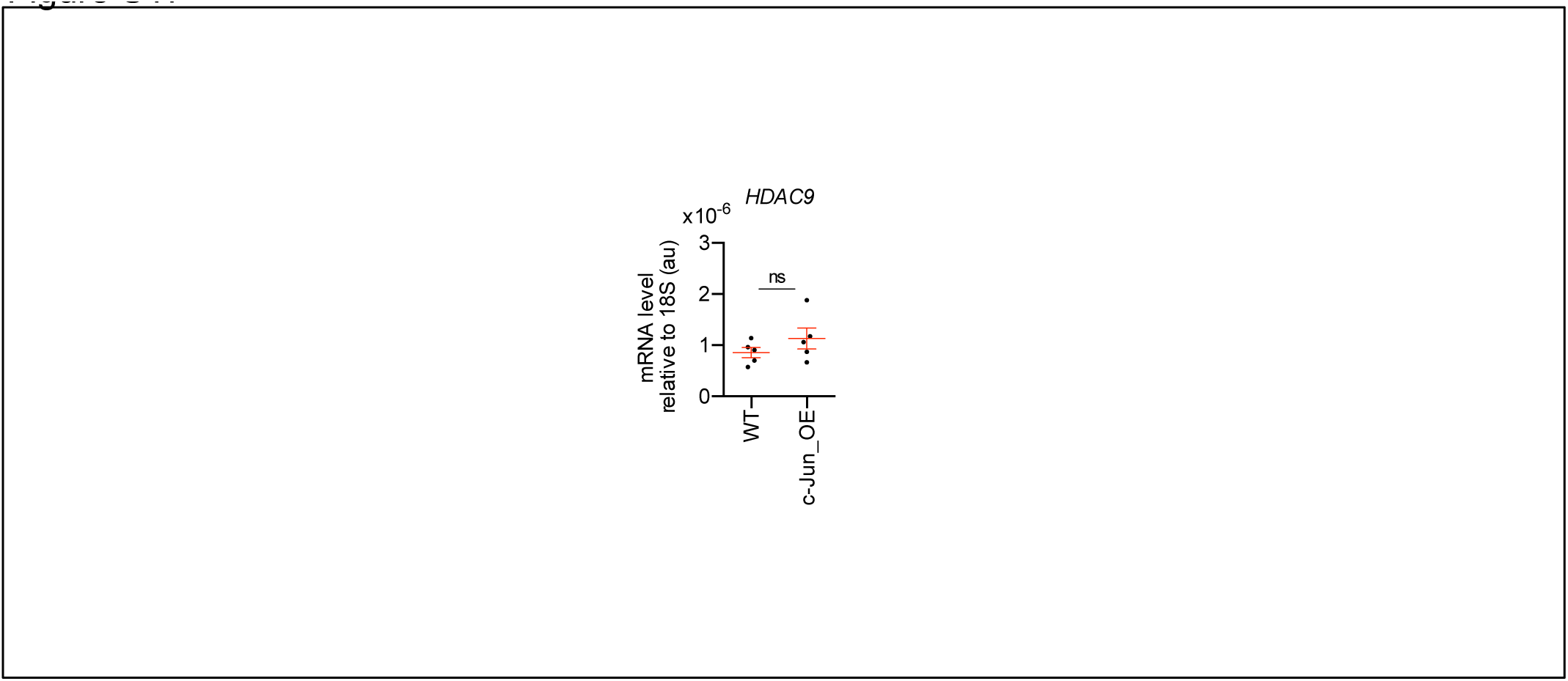
HDAC9 gene expression regulation. **A)** The *in vivo* overexpression of c-Jun in Schwann cells (c-Jun OE mice) did not modify the expression of HDAC9 gene in the PNS. RTqPCR with mouse-specific primers for HDAC9 was performed and normalized to 18S rRNA. Graph shows the relative expression of the mRNA normalized to the expression of 18S. A scatter plot is shown with the results obtained, which include also the mean ± SE. 5 mice per genotype were used. Data were analyzed with the unpaired t-test. Primer sequences are listed in Source data section online (Key Resources Table). (* P<0,05; ** P<0,01; *** P<0,001; ns: no significant). See Source data section online (Graphs source data) for more details.

